# STAT: A multi-agent framework for integrated and interactive spatial transcriptomics analysis

**DOI:** 10.64898/2026.05.01.722244

**Authors:** Yuheng Chen, Shi Han, Zitong Chao, Yuyao Liu, Fan Zhang, Hao Chen, Jiguang Wang, Jiashun Xiao, Can Yang

## Abstract

Spatial transcriptomics analysis often involves a myriad of computational methods across diverse platforms, leading analysts to spend excessive time on data assembly rather than deriving biological insights. Current AI solutions tend to either oversimplify spatial data into generic single-cell tables or operate autonomously without opportunities for intermediate review, thus hindering the visual and iterative analyses essential for spatial biology. In response to these challenges, we introduce STAT, a multi-agent framework designed to make spatial analysis more conversational and user-friendly while maintaining transparency and control. STAT integrates a persistent session, a shared interactive tissue viewer, and a staged skill-aware pipeline, enabling a more intuitive analytical experience. In a comprehensive benchmark evaluation encompassing eleven analytical task categories across three spatial platforms and both cell- and spot-resolution data, STAT demonstrates superior performance compared to a vanilla large language model and existing autonomous spatial analysis agents, excelling in task completion, analytical quality, and token efficiency. Notably, STAT enables multi-task spatial analysis of a mixed-resolution breast cancer cohort and successfully reproduces key findings from a published Visium HD colorectal cancer study using natural language prompts alone. STAT thus facilitates trustworthy and scientifically rigorous spatial transcriptomics analysis, allowing researchers to focus more on biological interpretation.

## Introduction

Spatial transcriptomics (ST) measures gene expression at single-cell or spot-level resolution while retaining the spatial context of each measurement within intact tissue sections [1], [2]. Its use has grown in cancer biology, neuroscience, immunology, and developmental biology [3]. However, biologists, bioinformaticians, and clinicians face a fragmented workflow. Each platform comes with its own conventions, and numerous incompatible methods exist for various analytical steps, including tissue-domain identification [4], [5], [6], cell-type deconvolution and annotation [7], [8], [9], [10], [11], [12], cell-cell communication [13], [14], [15], spatially variable gene detection [16], [17], [18], niche detection [19], [20], batch integration [21], [22], [23], and spatial registration [24], [25]. Analysts must select appropriate methods, prepare data in the required format, set platform-specific parameters, and visualize results in tissue space to guide subsequent steps.

Consequently, much of the analyst’s time is spent on assembly and visualization rather than on biological interpretation.

The development of biomedical AI agents aims to fulfill the diverse needs of biologists, bioinformaticians, and clinicians. However, existing agents exhibit significant limitations in integration and interactivity, which hinder researchers’ abilities to focus on specific regions, refine their inquiries, and adapt analyses in real time. General-purpose systems like Biomni [26], CellAgent [27], CellVoyager [28], and scAgent [29] can generate and execute code but oversimplify ST data as generic single-cell tables, overlooking crucial spatial coordinates and failing to distinguish between cell-resolution and spot-level analyses. In contrast, systems specifically designed for ST, such as ChatSpatial [30], connect large language models (LLMs) with existing tools through a command-line interface, while SpatialAgent [31] and STAgent [32] autonomously generate end-to-end reports. However, these systems frequently regenerate datasets with each query, lack adequate tissue visualizations, fragment analyses into isolated snippets, and restrict users to their existing tools. In particular, autonomous systems like Biomni and SpatialAgent incur significant token and time costs due to lengthy tool chains, limiting user oversight of intermediate processes. What is critically required for effective ST analysis is an integrated and interactive system. Integration maintains a persistent analytic state that enables the agent to access both the dataset and the tissue displayed in the graphical user interface (GUI) coherently. Interactivity should support multi-turn exchanges, allowing researchers to specify regions of interest, clarify queries, and inspect or rerun generated code and visualizations. Without this cohesive framework, researchers face challenges in engaging in a productive exploration cycle characterized by hypothesis formulation, real-time verification, and iterative method refinement.

In this study, we introduce STAT, a multi-agent framework specifically designed for ST that effectively integrates and enhances user interactions throughout the analytical process. STAT optimizes the workflow by loading the dataset once at the beginning of each session, organizing it as a single object whose platform, data level, available annotations, and other properties can be queried at every stage of analysis. This integration facilitates a GUI between the researcher and the agent, enabling real-time spatial visualization of gene expression, cell types, and deconvolution proportions, all of which dynamically update in response to modifications made by the agent. Moreover, STAT prioritizes interactivity, allowing users to convey their analytical intent without needing to repeatedly specify data attributes. As users delineate regions on the canvas, these areas are designated as regions of interest (ROIs), which the agent subsequently references in future queries. In instances of ambiguity, the agent seeks clarification rather than making assumptions, ensuring a more intuitive interaction. Additionally, all generated code and visualizations are compiled into a self-contained, executable notebook, empowering users to review, rerun, or refine their analyses. Rather than replacing the analyst, STAT streamlines the assembly and visualization processes that typically consume a significant portion of analysis time, while also providing essential support for the biological interpretations that ensue. Thus, comprehensive ST analyses can be conducted as a conversational exchange, maintaining the transparency and extensibility characteristic of traditional scripted pipelines. This innovative approach marks a significant step forward in the integration of AI within the realm of ST, enabling researchers to achieve their analytical goals more efficiently and effectively.

To assess whether STAT generalizes across platforms, tissues, and analytical tasks, we evaluate it in three complementary ways. First, we benchmark STAT against a vanilla LLM, Biomni, and SpatialAgent on 40 queries spanning eleven analytical task categories — for example, tissue-domain identification, cell-type annotation and deconvolution, cell-cell communication, and spatially variable gene detection — evaluated across three datasets drawn from three spatial platforms and both data resolutions: the spot-level Visium dorsolateral prefrontal cortex (DLPFC) dataset, and the cell-resolution MERFISH mouse-brain and Xenium breast cancer datasets. STAT achieves the highest task-completion rate and analysis quality under both LLM-based and ground-truth evaluations, and its competitors’ recurrent failure modes (silent wrong-tool selection, looped trial-and-error) illustrate how STAT’s architectural decisions contribute to this advantage.

Second, we dissect STAT’s multi-agent architecture — the persistent session, the staged skill-aware pipeline (planner, programmatic skill filter, semantic skill matcher, prerequisite verifier, code executor, and biological analyzer), and the classified error reflector — and show that its reliability survives a change of language-model backbone across seven frontier models, isolating the contribution of architecture from that of any particular model. Third, we apply STAT end-to-end to two disease cohorts: a mixed-resolution breast cancer cohort [33] on which the agent runs a diverse suite of spatial analyses through natural language dialogue and drawn-on-tissue ROIs, and a Visium HD colorectal cancer (CRC) cohort [34] on which we reproduce the central analysis of the original study from natural language prompts alone. Together, these evaluations show that STAT not only excels on standardized benchmarks but also supports the integrated, interactive spatial analysis that prior agent systems have not made possible.

## Results

### STAT enables integrated and interactive spatial transcriptomics analysis

STAT is a multi-agent framework designed to make ST analysis conversational. It combines a persistent session that holds the dataset, a shared interactive tissue viewer, and a staged multi-agent pipeline that plans and executes analyses, integrating three elements that are usually kept separate: the data object, the visual analytic workspace, and the reasoning layer. In contrast to general-purpose biomedical agents that flatten spatial data into generic single-cell tables, and to autonomous spatial agents that deliver an end-to-end report before the user can intervene, STAT integrates these three layers so that they reinforce one another: the session exposes explicit data attributes that the pipeline checks before choosing a method, the shared viewer lets the researcher steer analysis without restating the data, and the turn-by-turn dialogue keeps biology, code, and figures inside a single audit trail. STAT thereby shifts analyst time away from assembly and visualization toward biological interpretation, while producing an executable notebook that preserves the reproducibility of a scripted pipeline.

A STAT session is built around a single in-memory data object, holding the loaded slices, images, and their metadata, that both the researcher and the agent refer to throughout the conversation (Fig. 1, Methods). Starting from one or more ST slices, associated staining images, and an LLM API key, STAT auto-validates and casts each input into a persistent session in which platform, modality, data level (cell or spot), cell or spot count, and existing annotations are machine-readable attributes that later agent roles query without re-inspecting the raw data. The same session is surfaced to the researcher as a three-panel web interface (Fig. 3a, described in detail in a later section): an interactive tissue canvas with live overlays for cell types, gene expression, and deconvolution proportions; a side panel of slice, modality, cell-type, and region-of-interest controls; and a chat widget. Because the canvas and the session are two views of one state, regions the user draws on the tissue become named references (e.g., “ROI_1 on slice 1”) that the agent can read, and analyses the agent performs become overlays the user can inspect. A single button exports the full trace as a self-contained, executable Jupyter notebook, so each conversational session remains reproducible as a scripted workflow.

**Figure 1.**
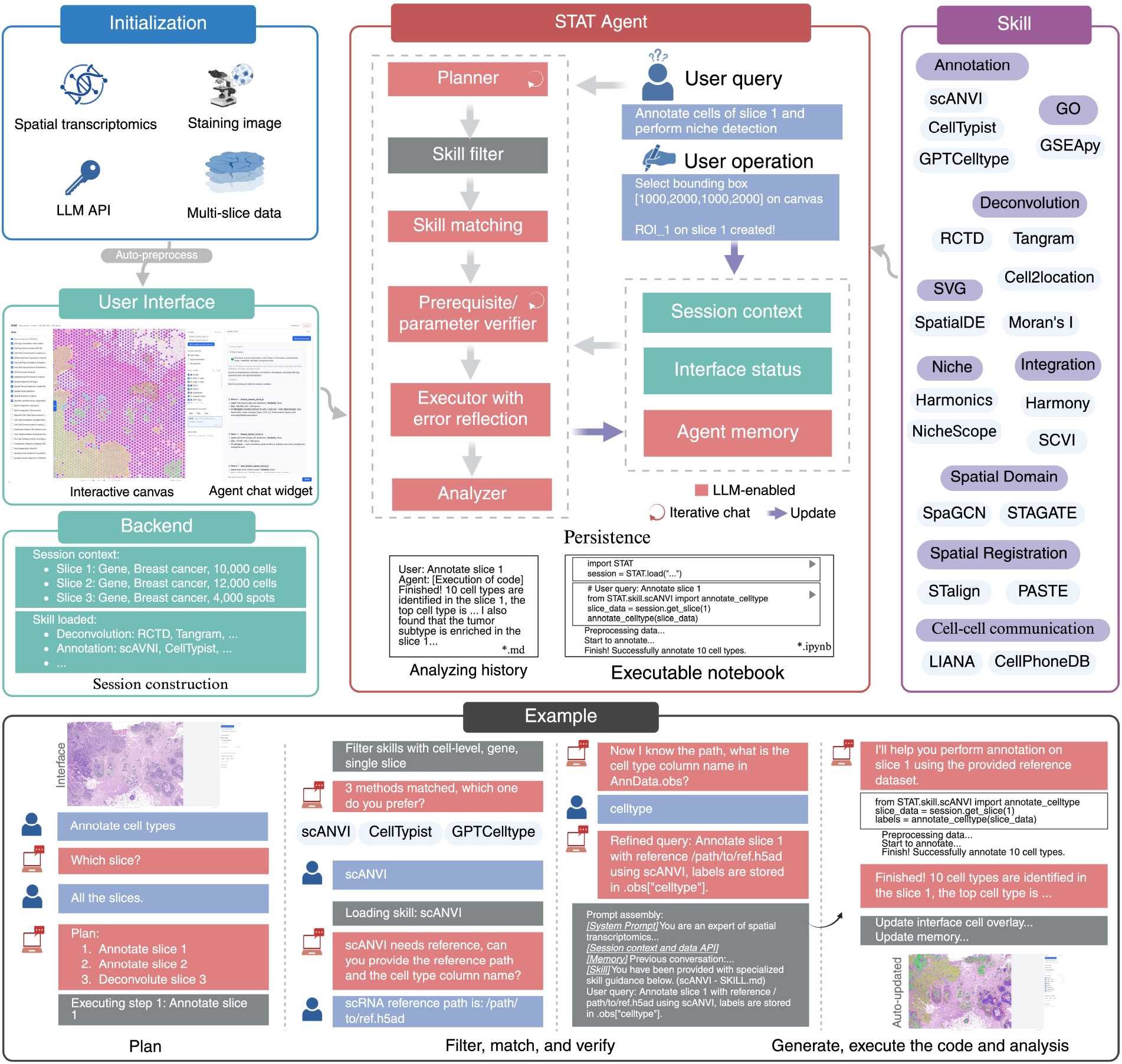
Overview of STAT. STAT loads one or more spatial transcriptomics slices, paired staining images, and an LLM API key (Initialization, top-left), auto-preprocesses them, and casts each input into a Backend session (bottom-left) in which every slice carries explicit modality, data level (cell or spot), cell or spot count, and existing annotations. The Skill registry (right) contains 27 self-describing analysis methods grouped by category (cell-type annotation and deconvolution, GO and pathway enrichment, spatially variable gene detection, niche detection, batch integration, spatial domain identification, spatial registration, and cell-cell communication, among others). The same session is exposed to the user through a three-column web interface (User Interface) and to the agent through a multi-agent pipeline (centre) in which a planner, skill filter, skill matcher, prerequisite/parameter verifier, executor with error reflector, and analyzer share a persistent context (Session context, Interface status, Agent memory). LLM-driven roles are coloured red; the circular-arrow icons mark roles that may surface clarification turns to the user; the purple update arrow marks the executor’s write-back to the agent memory. Every session is exported as a Markdown analysis history (`*.md`) and an executable Jupyter notebook (`*.ipynb`) (Persistence). The bottom strip walks one query end-to-end on a Xenium breast cancer slice (Example): the planner expands “Annotate cell types” into per-slice steps; the filter narrows the registry to skills compatible with the loaded data (cell-level, gene, single slice); the matcher offers three compatible methods (scANVI, CellTypist, GPTCelltype) and the user selects scANVI; the verifier collects the reference path and the cell-type column name; the executor runs the chosen method and updates the canvas overlay automatically. The four sources read by the assembled prompt at code-generation time — system prompt, session context and data API, conversation memory, and the loaded skill — are shown in the prompt-assembly inset.

This design is realized through a multi-agent architecture in which specialized roles (planning, skill filtering and matching, prerequisite verification, code generation, error reflection, and biological interpretation) share the session context and a memory of prior steps (Fig. 1). Two key features of this architecture are central to STAT’s reliability. First, each analysis method is wrapped as a self-describing skill: a module that contains the instructions the language model uses to invoke the method and declares, in a machine-readable form, the data conditions under which the method is applicable (modality, data level, slice count). Unlike tool-schema frameworks such as LangChain [35] or the Model Context Protocol (MCP) [36], in which the language model semantically chooses from all registered tools, STAT excludes methods incompatible with the loaded data and lets the model choose among the compatible subset; new methods are added to the system by writing a single new file. Second, skill selection is staged: planning, filtering, semantic matching over the compatible subset, and prerequisite collection all precede code generation. When code runs, the error reflector classifies failures as mechanically fixable or unsupportable, repairing the former and surfacing the latter as a human-readable explanation rather than looping indefinitely. Reliability therefore lives in the architecture rather than in prompt engineering, as confirmed by an ablation in which STAT maintains performance across different frontier language models (Fig. 3d,e).

STAT supports diverse ST platforms, including Visium [37], Visium HD [34], Xenium [38], MERFISH [39], and Stereo-seq [40], and handles cell-resolution and spot-level data uniformly within the same session. Its skill library covers the analytical categories that spatial studies routinely rely on — spatial domain and niche detection, cell-type annotation and deconvolution, spatially variable gene detection, cell-cell communication, batch integration, spatial registration, and pathway enrichment — and is extensible by adding a single Markdown file per new method (Fig. 1; Supplementary Table 12). The framework integrates with AnnData, Scanpy [41], and Squidpy [42], and every session can be exported as a Markdown analysis history or an executable `.ipynb` notebook for auditing, re-running, or sharing with collaborators who prefer a scripted workflow. The STAT software and documentation are publicly available at https://github.com/YangLabHKUST/STAT-agent, and a live web demo is hosted at https://huggingface.co/spaces/CyhVVVV/stat-agent-demo. A side-by-side comparison with seven prior biomedical and spatial analysis agents [26], [27], [28], [29], [30], [31], [32] is given in Supplementary Tables 1 and 2.

The Results below are organised in three parts. We first benchmark STAT against a vanilla LLM, Biomni, and SpatialAgent across eleven analytical task categories and three datasets spanning three platforms and both data resolutions. We then dissect the multi-agent pipeline that underlies this advantage — planner, skill filter, semantic skill matcher, prerequisite verifier, code executor with error reflector, and biological analyzer

— and show that its reliability is preserved across seven frontier language-model backbones, demonstrating that STAT’s robustness is a property of its pipeline rather than of any particular backbone. Finally, we apply STAT to two disease studies. The first study centers on a mixed-resolution breast cancer dataset [33] that integrates two cell-resolution replicates from Xenium with a Visium spot-level slice. Through natural language dialogue and tissue-canvas operations, we demonstrate that STAT enables a wide spectrum of spatial analyses. The second study is a published Visium HD CRC dataset [34], where we successfully replicate the key findings using STAT.

### STAT outperforms competing AI agents on a benchmark spanning eleven tasks and three spatial platforms

To test whether STAT outperforms existing alternatives on real ST analyses, we benchmarked it against three systems chosen to span the relevant design space: a vanilla LLM as the no-agent baseline, the general-purpose biomedical agent Biomni [26], and the autonomous spatial agent SpatialAgent [31]. All four systems were run on the same Claude Sonnet 4 backbone to control for the language model itself. We assembled a comprehensive query set of 40 queries covering the 11 analytical task categories that are routine in ST analysis (cell-type annotation, deconvolution, spatial-domain identification, niche detection, spatially variable gene (SVG) detection, cell-cell communication, batch integration, spatial registration, Gene Ontology (GO) enrichment, neighborhood enrichment, and trajectory analysis) under three query types (clear single-task, vague single-task, and multi-step-task), plus a 6-query boundary-case category covering infeasible, out-of-scope, and missing-prerequisite tasks. To make the evaluation comprehensive across current ST technology, the queries were issued on three datasets that together span the three major platforms (Visium, MERFISH, Xenium), both spatial resolutions (spot-level and cell-resolution), and two tissue contexts (cortex and breast cancer): a Visium DLPFC cohort [43] (spot-level, four slices), a MERFISH mouse-brain dataset [44] (cell-resolution, two slices), and a Xenium breast cancer sample [33] (cell-resolution, 160K cells); the full query list is in Supplementary Table 3 and a category-by-difficulty map in Fig. 2a,b,c. Each (query, system) pair produced one stored run, giving 160 runs in total, each anonymised and rated for binary success, integer quality (1–5), and per-system rank by Claude Opus 4.6, the strongest reasoning model available at the time of evaluation, against task-specific success criteria fixed in advance (Methods; Supplementary Table 4). STAT ranked first on all four aggregate metrics: 39/40 queries completed (vs. 37 SpatialAgent, 35 Biomni, 9 vanilla LLM); highest average quality (4.5 vs. 4.2 / 3.8 / 3.8); lowest average rank (1.3); and the best system on 31/40 queries (Fig. 2d, Supplementary Fig. 1). Its lead was broad across task categories rather than concentrated in any one (Fig. 2e), and the margin widened sharply on the boundary set (83% success for STAT vs. 67% / 33% / 17%; Fig. 2g).

**Figure 2.**
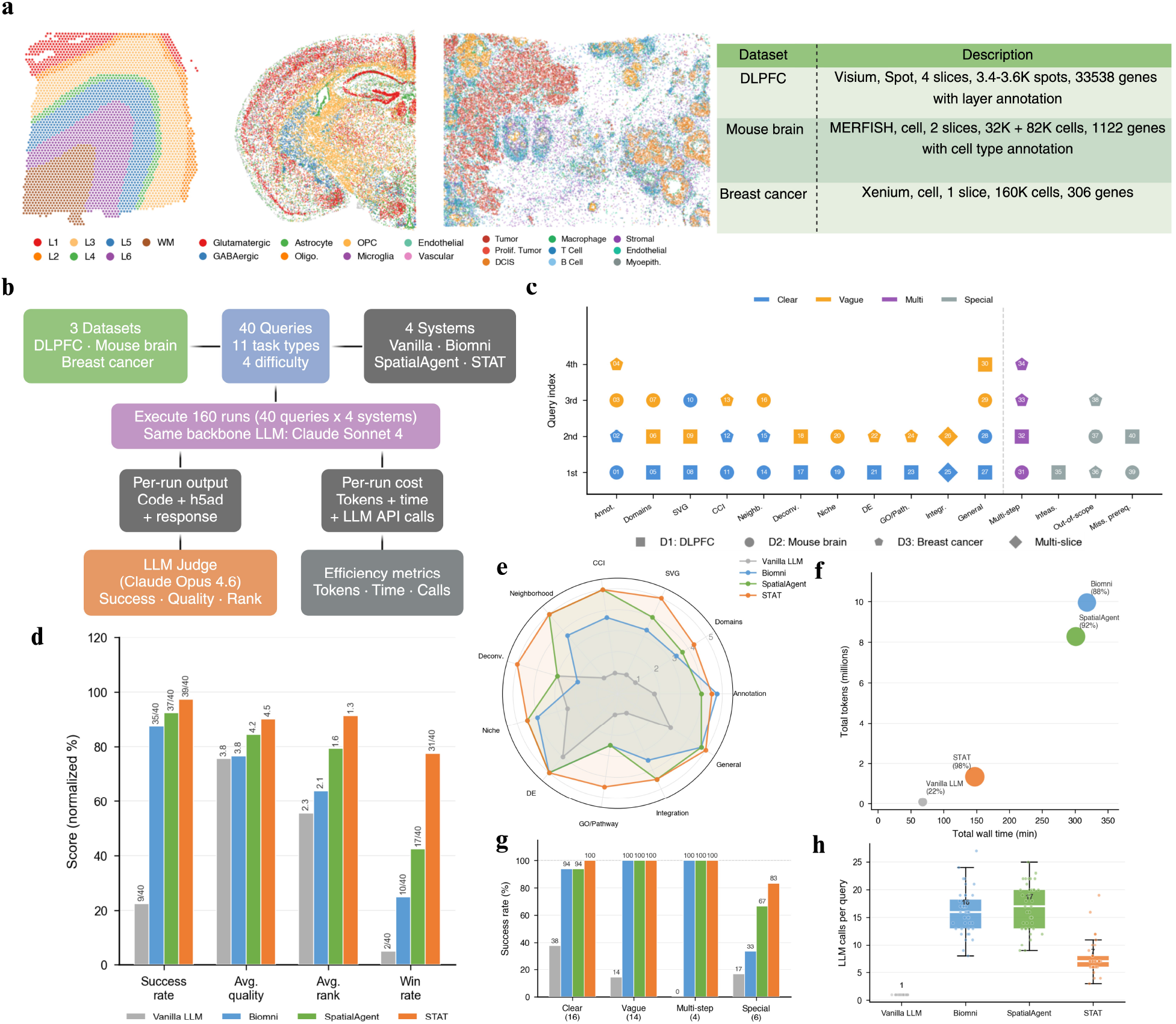
STAT outperforms competing AI agents on a benchmark spanning eleven analytical tasks, three spatial platforms, and both data resolutions. **a**, Three benchmark datasets used in this study: a Visium human dorsolateral prefrontal cortex (DLPFC) cohort with manually curated cortical-layer annotations (4 slices, 3.4–3.6K spots, 33,538 genes), a MERFISH mouse-brain dataset with curated cell-type annotations (2 slices, 32K and 82K cells, 1,122 genes), and a 10x Genomics Xenium human breast cancer slice (1 slice, 160K cells, 306 genes). **b**, Benchmark design. Forty queries — three query types (clear single-task, vague single-task, and multi-step-task) plus a separate boundary-case category (infeasible, out-of-scope, missing-prerequisite) — across eleven task categories were issued to four systems (vanilla LLM, Biomni, SpatialAgent, STAT) using the same backbone (Claude Sonnet 4), yielding 160 runs. Per-run output was scored by Claude Opus 4.6, the strongest reasoning model available at the time of evaluation, as an anonymised expert judge for success, quality (1–5), and rank; per-run cost was used to derive efficiency metrics. **c**, Per-query map of dataset × task × query type (Clear, Vague, Multi, Special). Marker shape encodes dataset; the 30 core single-task queries (T1–T11) lie to the left of the dashed divider, and the remaining 10 queries (4 multi-step plus 6 boundary cases — infeasible, out-of-scope, and missing-prerequisite) lie to the right. **d**, Four aggregate metrics. Success rate (out of 40): 9, 35, 37, 39. Average quality (1–5): 3.8, 3.8, 4.2, 4.5. Average rank: 2.3, 2.1, 1.6, 1.3. Win rate (out of 40): 2, 10, 17, 31. **e**, Per-task-category mean quality scores (radar plot). **f**, Per-system efficiency: total wall time (x) versus total tokens (y); bubble size and percent labels encode success rate. **g**, Success rate by query-type category (query counts in parentheses). **h**, LLM API calls per query (medians: 1, 16, 17, 7). Detailed failure-mode analysis: Supplementary Table 5.

Inspecting the judge’s rationales alongside the generated code and stored outputs (Supplementary Table 5) revealed four recurring failure patterns in Biomni and SpatialAgent, each contrasting with how STAT handled the same query. The first pattern was the fabrication of outputs on infeasible tasks. On Q38 (predict patient survival from tissue), both competitors synthesised survival labels and trained classifiers on the fabricated outcomes, while STAT’s error reflector recognised the absence of real outcome data and surfaced the infeasibility as a readable explanation; on Q37 (RNA velocity on data without spliced and unspliced count layers), Biomni rebranded a spatial-gradient calculation as RNA velocity and SpatialAgent pivoted to a partition-based graph abstraction (PAGA) trajectory analysis, while STAT also failed, the only query STAT missed (we discuss it below). The second pattern was returning a manual annotation in place of a statistical test. On Q23 (GO enrichment on seven marker genes), both competitors returned a literature-derived functional categorisation of the genes rather than running an enrichment test, while STAT correctly invoked its `pathway-go-enrichment` skill (gseapy with Fisher’s exact test and Benjamini-Hochberg false discovery rate (BH-FDR) control). The third pattern was the use of a non-spatial proxy where a spatial-aware method was needed. On the spot-level deconvolution queries (Q17, Q18), Biomni’s non-negative least-squares (NNLS) implementation collapsed proportions onto a single dominant cell type in more than 75% of spots, treating the spot-as-mixture problem as label transfer; on SVG detection (Q09, Q10), Biomni scored candidates with a plain coefficient of variation rather than a spatial-autocorrelation statistic. STAT instead routed to RCTD for deconvolution and to SpatialDE [16] on spot-level Visium and Moran’s I [45] on cell-resolution Xenium for SVG detection, with each skill’s `filter_requirements` declaring its compatible data level. The fourth pattern was the substitution of an alternative input when a prerequisite was missing. On Q40 (test cell-type co-localisation on a slice that carries cortical-layer annotations but no `celltype` column), Biomni substituted Leiden clusters as ad-hoc cell-type labels and SpatialAgent used the brain-layer annotations as a cell-type proxy, while STAT’s verifier flagged the missing prerequisite before code generation.

The single STAT failure (Q37, RNA velocity) is itself worth examining. RNA velocity is not an ST method and has no corresponding skill in STAT’s registry (Supplementary Table 12), so the matcher returned no match and the pipeline fell back to raw LLM behaviour: the code generator called scVelo [46] without verifying that the required spliced and unspliced count layers were present, reproducing the same hallucination pattern as the autonomous competitors. This case illustrates a critical trade-off that STAT was designed to clarify. When a method is encapsulated as a skill, the combination of compatibility pre-filter, prerequisite verifier, and data-resolution-aware matcher effectively prevents the failure modes mentioned earlier. However, in situations where no skill currently addresses the request, STAT preserves the general-purpose flexibility of the language model, which we consider a feature rather than a flaw. This flexibility enables users to explore a broader range of analyses beyond the existing registry, albeit at the cost of potential hallucination risks inherent to any unguarded language model. The logical response, then, is not to deny queries that fall outside the registry but to expand it. Adding new methods is intentionally straightforward: each method can be incorporated as a single Markdown file with a YAML header (see Methods, Supplementary Table 12), making it easy to bring additional analyses under STAT’s framework of reliability.

Two further advantages of STAT emerge from the same benchmark. First, on the two tasks for which canonical ground-truth labels exist, namely spatial-domain identification on the cortical-layer-annotated DLPFC cohort and cell-type annotation on the MERFISH brain, STAT’s outputs reached the highest label agreement of the four systems (Supplementary Fig. 2), an external validation that does not depend on the LLM judge. Second, STAT was substantially cheaper to run. Biomni and SpatialAgent plan and self-critique over long tool chains, consuming 8–10M tokens, 300+ minutes of wall time, and a median of 16–17 LLM calls per query, whereas STAT’s staged pipeline, which commits to a compatible skill before generating code and exits the retry loop once the reflector classifies a failure as unfixable, finished the same benchmark in 150 minutes and 1.5M tokens with a median of seven LLM calls per query (Fig. 2f,h). Inspection of STAT’s wider output gallery confirms that its method choices were statistically appropriate at each resolution: RCTD [7] deconvolution on spot-level DLPFC (Supplementary Fig. 3); SpatialDE for SVG detection on the same cohort versus Moran’s I on cell-resolution Xenium breast cancer (Supplementary Fig. 4, Supplementary Fig. 5); niche detection on MERFISH recovering the classical anatomical regions of the mouse brain (Supplementary Fig. 6); and Harmony [21] integration producing a well-mixed DLPFC embedding and a well-integrated MERFISH embedding in which a lone residual cluster is biologically explained by a cell type present only in the second slice (Supplementary Fig. 7, Supplementary Fig. 8). Taken together, the benchmark shows that STAT’s lead extends not only to completion rate and quality but also to the classes of failure it avoids. Whether these gains depend on the choice of language-model backbone is addressed in the next section.

### Web interface and the multi-agent pipeline behind STAT’s reliability

The interactive workspace exposes STAT’s session and multi-agent pipeline through a three-column web interface (Fig. 3a). The left column is a skills configuration panel in which every registered analysis method is listed with its declared requirements (modality, data level, slice count) and prerequisites; the user can selectively enable or disable methods, and the same declarations drive the pipeline’s compatibility filter, so what the user sees is exactly what the agent is allowed to choose. The centre is an interactive tissue canvas that renders cells or spots over the staining image with real-time controls for opacity, point size, slice, and visualization mode (cell type, gene expression, or deconvolution proportions); a 3D viewer is available for multi-slice cohorts (Supplementary Fig. 9). Adjacent to the canvas, basic spatial visualization controls expose slice, modality, and cell-type subsets, while ROI selection tools (rectangle, polygon, free-hand) promote drawn regions into named references that the agent can act on across turns. The right column hosts a conversation widget in which planner clarifications, skill-selection prompts, verifier questions, and analyzer interpretations are surfaced inline with the executed code, and logs and notebook controls in the header export the full session as a Markdown analysis history or an executable Jupyter notebook. Each component is paired with a counterpart inside the pipeline: ROIs become named references the agent can query, the canvas state is read by the analyzer when answering region-specific questions, and the visualization mode updates automatically whenever the agent modifies the underlying data — so the surface the user works with and the state the agent reasons over are the same object rather than two parallel views that must be kept in sync.

**Figure 3.**
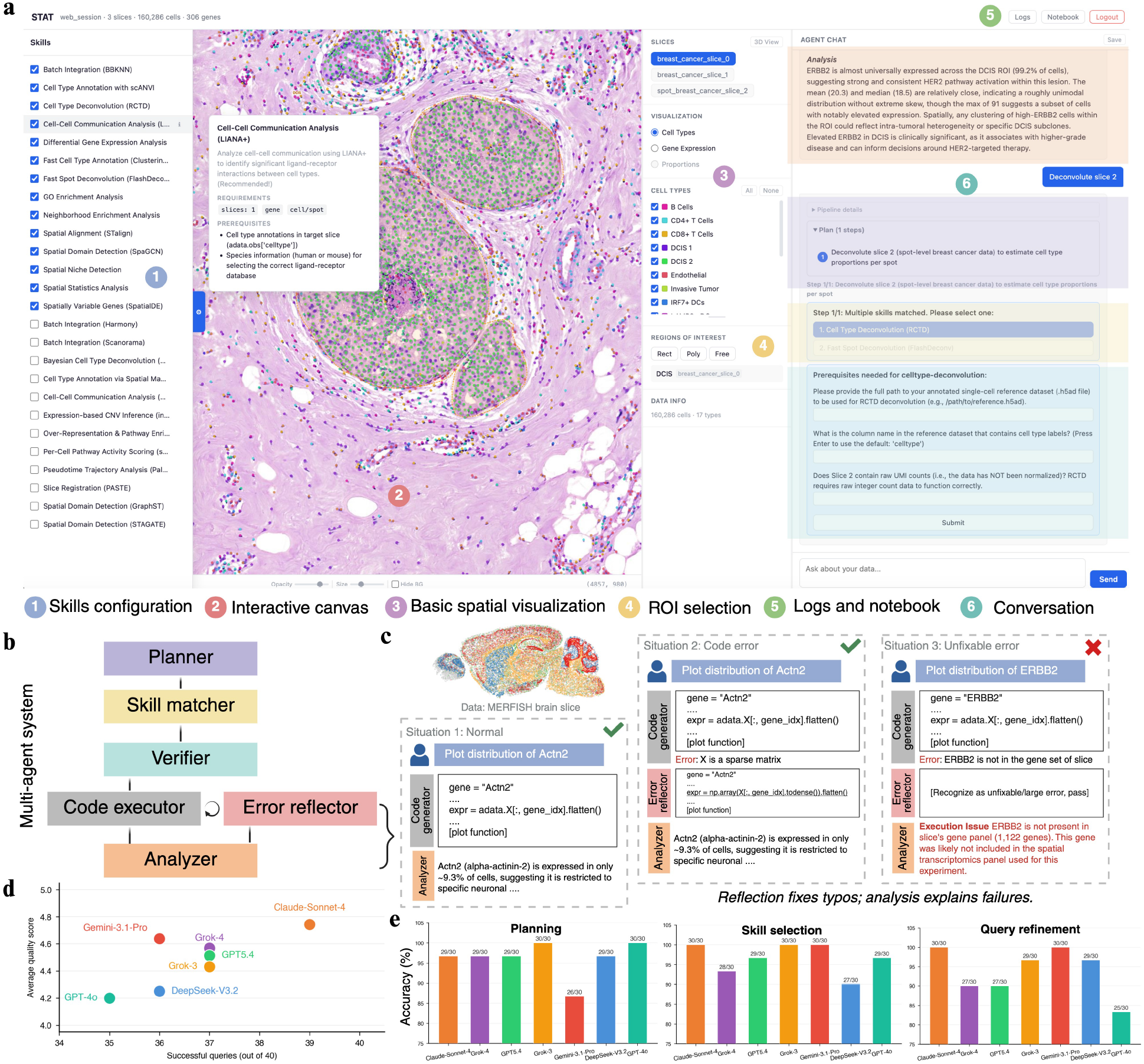
User-facing components and architecture-driven robustness of STAT. **a**, Three-column web interface for an annotated Xenium breast cancer session with six numbered callouts: ① skills configuration panel (left) — methods listed with declared requirements and prerequisites, enabled/disabled by the user; the hovered tooltip shows the LIANA+ skill metadata; ②interactive tissue canvas (centre) rendering cells over H&E with real-time controls; ③ basic spatial visualization controls (slice, modality, cell-type subset, mode); ④ ROI selection tools (rectangle, polygon, free-hand) promoting drawn regions into named references; ⑤ logs and notebook export buttons (top-right); ⑥conversation widget on the right surfacing clarifications, skill selection (RCTD vs FlashDeconv), and prerequisite collection inline with executed code. **b**, Multi-agent pipeline: planner →skill matcher →verifier →code executor with attached error reflector →analyzer. **c**, Three error-handling situations of the error reflector on a MERFISH mouse-brain slice: normal (left), fixable code error (centre, sparse-matrix indexing rewritten with `np.array(X[:, gene_idx].todense()).flatten()`), and unfixable error (right, ERBB2 absent from gene panel surfaced as a readable execution issue). **d**, Cross-LLM ablation on the same 40-query benchmark of Fig. 2 with seven frontier backbones (Claude Sonnet 4, GPT-5.4, GPT-4o, Grok-3, Grok-4, Gemini 3.1 Pro, DeepSeek V3.2). x-axis: successful queries (out of 40); y-axis: average judge-assigned quality (1–5). **e**, Per-stage accuracy (out of 30) of the planner, skill matcher, and refined-query stages on a dedicated 90-query pipeline test set (Supplementary Tables 7–10) for the same seven backbones. All backbones cleared 86.7% / 90% / 83.3% on the three stages respectively; per-failure analysis: Supplementary Table 11.

Reliability inside that loop is enforced by a small number of explicit roles — planner, skill matcher, verifier, code executor with error reflector, and analyzer (Fig. 3b) — each with a narrowly scoped responsibility. The error reflector is the role that determines how STAT responds when code execution does not succeed (Fig. 3c). On a MERFISH brain slice, three behaviors illustrate its scope. In the normal case (“Plot distribution of *Actn2*”), the generator produces correct code and the analyzer summarises the biology directly. In a fixable case, the same query first fails because the expression matrix is sparse and the indexing call returns the wrong type; the reflector classifies this as a mechanical error, rewrites the offending line to densify the slice gene matrix (`np.array(X[:, gene_idx].todense()).flatten()`), and the analyzer returns the same biological interpretation as if the error had never occurred. In an unfixable case (“Plot distribution of *ERBB2*”), the gene is absent from the MERFISH panel; the reflector recognises this as structural infeasibility rather than a typo, and the analyzer reports a readable execution issue (“*ERBB2* is not present in slice’s gene panel (1,122 genes); this gene was likely not included in the spatial transcriptomics panel used for this experiment”) rather than entering an indefinite retry loop. This split between mechanically fixable and structurally infeasible failures is what allows the pipeline to converge on either a correct answer or an honest refusal in a bounded number of turns, and it is the mechanism that prevents the fabrication failure modes observed in the autonomous competitors of Fig. 2.

To test how much of STAT’s reliability follows from this architecture rather than from a particular language model, we re-ran the agent on the same 40-query benchmark with seven frontier backbones — Claude Sonnet 4, GPT-5.4, GPT-4o, Grok-3, Grok-4, Gemini 3.1 Pro, and DeepSeek V3.2 [47] — and additionally evaluated the planner, skill-matcher, and refined-query stages independently on a dedicated 90-query pipeline test set (30 per stage across six synthetic test sessions of varying composition: one slice, two slices, or four slices; cell-level or spot-level; with or without celltype annotations; Supplementary Tables 7–10). End-to-end, every backbone completed at least 87.5% of the benchmark queries, with success rates ranging from 35/40 (GPT-4o) to 39/40 (Claude Sonnet 4) and quality scores tightly clustered above 4.0 (Fig. 3d, Supplementary Fig. 10, Supplementary Table 6). At the per-stage level, all seven backbones cleared 86.7% on the planner, 90% on skill matching, and 83.3% on refined-query generation (Fig. 3e, Supplementary Table 11). The residual failures were not random across models but concentrated in a small number of LLM-dependent patterns: planners that collapsed explicit “first … then …” chains into a single step (PA23–PA29), semantic matchers that returned an empty match for paraphrased but valid queries (PB18), and refined-query stages that occasionally hallucinated session metadata absent from the user query (PC25, PC27). Because the pipeline’s structural guards downstream — the verifier and the error reflector — either ask for clarification, fall back to an alternative skill, or surface a readable explanation, these stage-level slips did not propagate into comparable gaps in completed analyses, supporting the claim that STAT’s reliability lives in its architecture rather than in the prompt-level idiosyncrasies of any single language model.

### Conversational analysis of a mixed-resolution breast cancer cohort

Traditional ST workflows pay a fixed setup cost — searching methods, reading documentation, configuring environments, writing code, and tuning parameters — that must be paid again for every new question asked of the same dataset (Fig. 4a). The consequence is that analyses are usually planned linearly, revisions are expensive, and genuinely exploratory cycles are rare. STAT replaces this pattern with a session in which data are loaded once and subsequent analyses are initiated by natural language queries, each inheriting the state produced by the previous turn (Fig. 4a). To demonstrate the workflow end-to-end, we assembled a single session from three breast cancer slices of heterogeneous provenance: two Xenium cell-resolution replicates and one Visium spot-level slice, each paired with its H&E image (Fig. 4b). In the first turn, the user asked “Annotate cell types”. Rather than issuing a single annotation call, the planner recognised that the session contained two cell-resolution slices and one spot-level slice, and expanded the query into a three-step plan: scANVI [9] label transfer for the two Xenium slices and RCTD deconvolution for the Visium slice (Fig. 4b). The verifier collected the reference path and cell-type column name once and reused them across the three steps, yielding a consistent 17-type annotation across the cohort (Fig. 4b). The data-level distinction between annotation and deconvolution that a user would normally need to enforce by hand is absorbed here by the session, the planner, and the skill matcher acting together.

**Figure 4.**
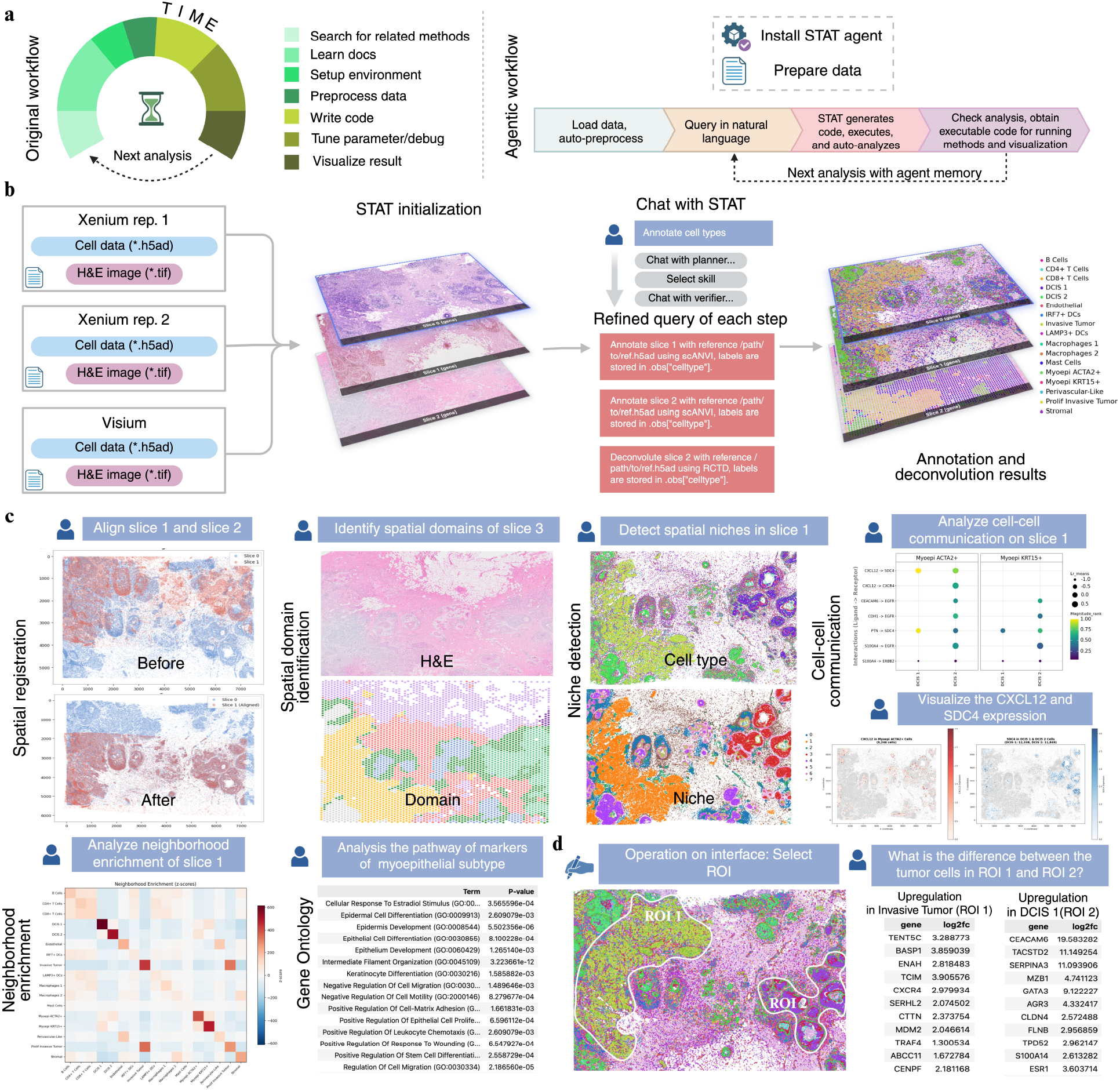
Conversational analysis of a mixed-resolution breast cancer cohort. **a**, Workflow comparison: conventional scripted workflow (left) repeats a fixed setup loop for every new analysis; STAT agentic workflow (right) loads data once and accepts subsequent analyses as natural language queries reusing agent memory. **b**, STAT initialization on a single mixed-resolution session containing two Xenium cell-resolution replicates and one Visium spot-level slice (left). The first user query “Annotate cell types” is decomposed by planner / matcher / verifier into three refined queries: scANVI annotation for the two Xenium slices and RCTD deconvolution for the Visium slice (centre), yielding a consistent 17-type annotation across the cohort (right). **c**, Diverse downstream analyses driven by single natural language queries: spatial registration of slices 1 and 2 (before/after); spatial domain identification on slice 3 (H&E vs SpaGCN domain map); niche detection on slice 1 (cell type vs eight-niche partition); cell-cell communication on slice 1 (LIANA-ranked ligand–receptor pairs between Myoepi *ACTA2*+ / *KRT15*+ and DCIS 1 / DCIS 2); follow-up gene-expression visualization (*CXCL12* in Myoepi *ACTA2*+, *SDC4* in DCIS); neighborhood enrichment z-score matrix; Gene Ontology enrichment on myoepithelial markers (gseapy Fisher’s exact + BH FDR). **d**, ROI-driven differential expression: ROI 1 (invasive tumor) and ROI 2 (DCIS 1) drawn directly on canvas; the natural language query “What is the difference between tumor cells in ROI 1 and ROI 2?” runs Wilcoxon DE restricted to tumor cells in each region, returning genes upregulated in invasive tumor (left, including *CXCR4, ENAH, CTTN, BASP1*) and in DCIS 1 (right, including *CEACAM6* log2fc 19.6, *TACSTD2* 11.1, *GATA3* 9.1, *ESR1* 3.6) — recovering the canonical DCIS-vs-invasive transcriptional separation.

From the annotated session, a sequence of downstream questions was answered in the same conversational register (Fig. 4c). Asked to “align slice 1 and slice 2”, STAT produced a before/after overlay in which the two Xenium tissues come into register. Asked to identify spatial domains on slice 3, it returned a domain map on the Visium spot grid whose boundaries follow the histological architecture visible in the paired H&E image. On slice 1, niche detection partitioned the tissue into eight microenvironments distinguishing tumor, stroma, and ductal carcinoma in situ (DCIS) compartments; a follow-up ligand–receptor analysis identified *CXCL12*→*SDC4, CXCL12*→*CXCR4, CEACAM6*→*EGFR, PTN*→*SDC4*, and *S100A4*→*ERBB2* among the top-ranked communications between the two myoepithelial subtypes and the DCIS populations, and a subsequent request to visualise *CXCL12* and *SDC4* placed the two genes back onto the tissue, confirming that the inferred ligand is expressed in Myoepi *ACTA2*+ cells and the cognate receptor in DCIS 1 and DCIS 2 — a one-turn cross-check that scripted pipelines typically require several additional scripts to assemble. Neighborhood enrichment returned a z-score matrix with strong self-enrichment for DCIS 1, Invasive Tumor, and the myoepithelial subtypes, mirroring their histological grouping; and a GO enrichment on markers of the myoepithelial subtypes returned a coherent epithelial-differentiation and cell-migration programme, with Intermediate Filament Organization at *p* = 3 × 10^™12^, Keratinocyte Differentiation, Epithelial Cell Differentiation, and multiple cell-motility and cell–matrix-adhesion terms among the top hits. In every case the method was chosen automatically by STAT’s skill matcher according to the data level of the slice the user had named, so the breadth of analyses exercised here was reached without the user specifying a method, a parameter file, or a data-format conversion.

Tissue-level analyses are complemented by region-level comparison through STAT’s drawn-ROI primitive (Fig. 4d). Rather than selecting cells by cluster index or expression threshold, a user can delineate regions directly on the tissue canvas and refer to them in subsequent natural language queries. The same canvas can render the underlying state in any of three modes — cell-type maps, single-gene expression (for example, *ERBB2* across the breast cancer slice), or per-spot deconvolution proportions — with the user switching among them interactively and without re-running the underlying analyses (Supplementary Fig. 11, Supplementary Fig. 12, Supplementary Fig. 13). We drew two regions on an annotated Xenium slice — ROI 1 over an invasive tumor pocket and ROI 2 over a DCIS lobule — and asked “What is the difference between the tumor cells in ROI 1 and ROI 2?”. STAT restricted the differential-expression test to tumor cells intersecting each region and returned two biologically interpretable gene lists. Invasive Tumor cells in ROI 1 were enriched for migration- and invasion-associated genes (*CXCR4, ENAH, CTTN, BASP1, TRAF4, MDM2*), while DCIS cells in ROI 2 were enriched for canonical DCIS and luminal-epithelial markers (*CEACAM6* at log2-fold-change 19.6, *TACSTD2* at 11.1, *GATA3* at 9.1, *ESR1* at 3.6, *CLDN4, SERPINA3*).

The two gene lists reproduce the canonical DCIS-versus-invasive transcriptional distinction, obtained from a single natural language question that referred to two hand-drawn ROIs. A more elaborate variant of this region-driven workflow is shown in Supplementary Fig. 14, in which three ROIs drawn on Invasive Tumor, DCIS 1, and DCIS 2 enriched regions supported a sequence of natural language queries that first returned the cell-type composition stacked across the three regions and then compared tumor and myoepithelial gene expression across the same regions as a per-region heatmap — recovering the canonical DCIS 1 →DCIS 2 →Invasive Tumor progression and the accompanying myoepithelial breakdown — all without leaving the conversational interface. This kind of visual-hypothesis-to-quantitative-comparison turn is what the integrated workspace makes possible: drawn regions, cell-type annotations from earlier turns, and the session’s knowledge of which slice the user is viewing are all pieces of the same state, so the agent can act on them together rather than asking the user to reconcile them in code.

### Agentic reproduction of a published Visium HD colorectal cancer study

A critical test of STAT is whether it can reproduce a published end-to-end study from natural language prompts alone, on data it has never been told about. We posed this test on a recent Visium HD CRC study (Oliveira et al., Nat. Genet. 2025) [34], which characterises the spatial organisation of tumor, periphery, and surrounding tissue across three CRC patients (P1, P2, P5) at 8-µm spot resolution and identifies functionally distinct *SPP1*^+^ and *SELENOP*^+^ macrophage subtypes that structure the tumor microenvironment. Reproducing the study’s central characterisation of the *SPP1*^+^ / *SELENOP*^+^ macrophage axis at the tumor periphery — and the divergent transcriptional, pathway, and ligand–receptor programmes that distinguish the two subtypes — requires a chain of dependent analyses: cell-type deconvolution, spatial region labelling, marker-based subtyping of a specific cell population, differential expression and pathway enrichment, niche-restricted comparisons, and ligand–receptor inference, each of which would, in the original workflow, demand its own scripted pipeline (Fig. 5). We loaded the three patient slices into a single STAT session and issued a series of natural language prompts that walked the agent through six analytical steps; the prompts described the desired procedure but did not name tools and did not reference the original paper (Supplementary File 1). One deliberate deviation from the original analysis was made: *SELENOP*^+^ macrophages were defined here as the macrophage spots that express *SELENOP* rather than as all non-*SPP1*^+^ macrophages, so that both subtype labels rest on positive marker evidence rather than on an exclusion rule.

**Figure 5.**
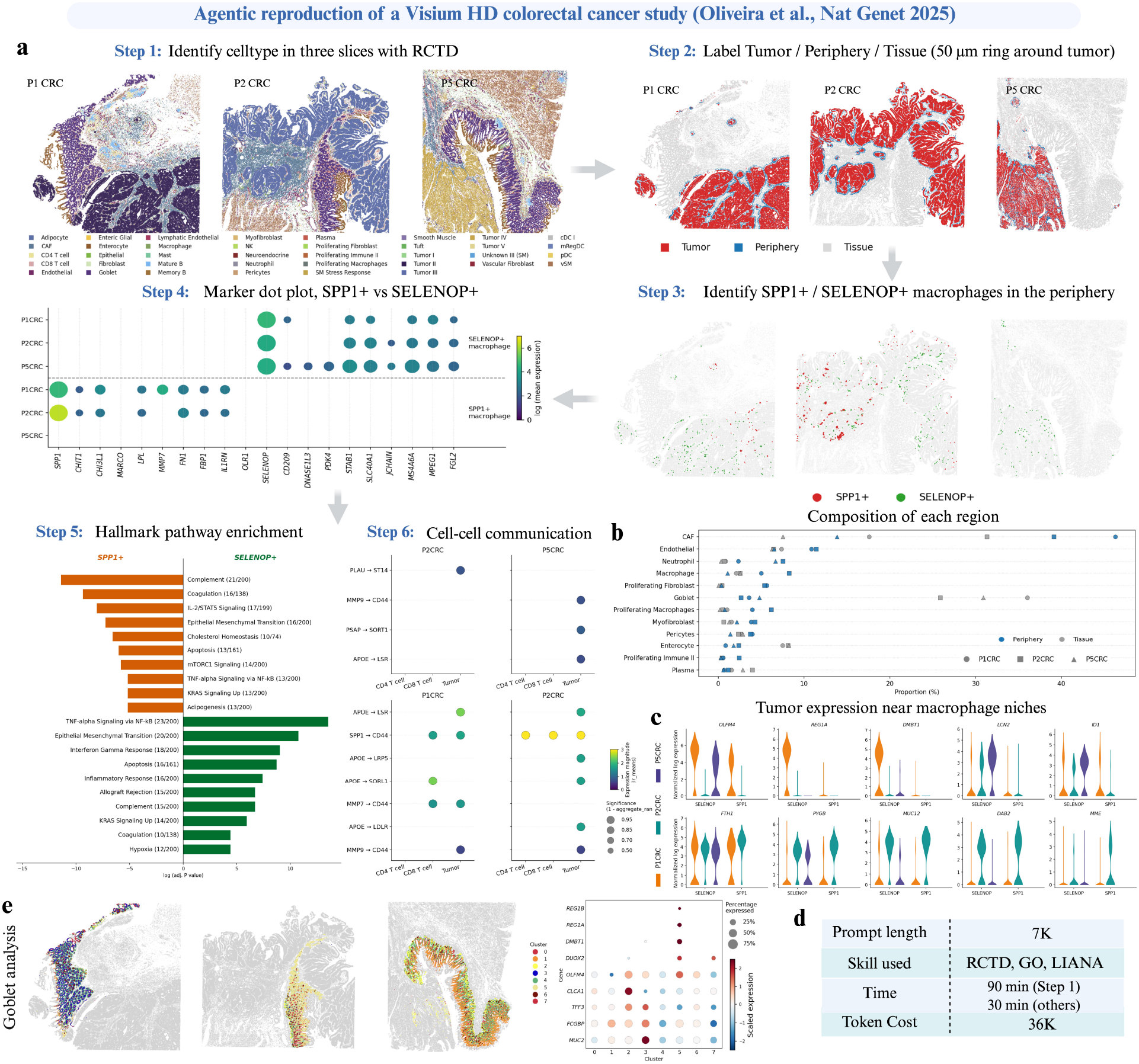
Agentic reproduction of a published Visium HD colorectal cancer study. Reproduction of the central figures of Oliveira et al. (Nat. Genet. 2025) on three patient slices (P1 CRC, P2 CRC, P5 CRC; 8 µm spot-level Visium HD) loaded into a single STAT session. Six analytical steps were issued as natural language prompts; the agent did not see the original paper. **Step 1**, RCTD deconvolution of each slice against the matched single-cell reference. **Step 2**, region labelling — each spot assigned to Tumor, Periphery, or Tissue, with the periphery defined as a 50 µm ring around core tumor seeds (≥25 tumor neighbours within 50 µm). **Step 3**, identification of *SPP1*^+^ and *SELENOP*^+^ macrophages within the periphery. **Step 4**, Wilcoxon marker dot plot between the two macrophage subtypes. **Step 5**, three downstream analyses on the marker sets: hallmark pathway enrichment via gseapy, tumor expression near macrophage niches, and goblet-cell sub-clustering with canonical and inflammatory markers (*REG1A, REG1B, DUOX2, DMBT1*). **Step 6**, LIANA cell-cell communication on the tumor-microenvironment object combining periphery spots and abutting tumor spots. Summary panel: three named skills (RCTD, GO/gseapy, LIANA), 7K input / 36K output tokens, 90 min for Step 1 and 30 min per subsequent step on a single workstation. *SELENOP*^+^ macrophages here are macrophage spots expressing *SELENOP*, rather than all non-*SPP1*^+^ macrophages as in the original analysis.

Step 1 invoked the RCTD skill to deconvolute each slice against the matched single-cell reference, returning per-spot cell-type proportions consistent with the published annotations (Fig. 5, Step 1). Step 2 derived a tumor/periphery/tissue label by identifying core tumor seeds (spots whose cell-type label matched each slice’s dominant tumor subtype and that had ≥25 tumor neighbours within 50 µm), expanding a 50-µm periphery ring around them, and assigning everything else as surrounding tissue; the resulting periphery composition recapitulated the immune enrichment reported by the original study (Fig. 5, Step 2). Step 3 split periphery macrophage spots into *SPP1*^+^ and *SELENOP*^+^ subtypes from raw-count expression of the two anchor genes, and Step 4 ran a Wilcoxon test between the pooled populations and rendered the discriminating markers as a per-patient dot plot that recovers the macrophage marker programmes reported by the original publication (Fig. 5, Steps 3–4). Step 5 then chained three further analyses on those marker sets: a two-group hallmark-pathway enrichment between *SPP1*^+^ and *SELENOP*^+^ markers via the GO enrichment skill, a niche-restricted differential-expression test between tumor spots immediately adjacent to each macrophage population, and a goblet-cell sub-clustering with canonical and inflammatory markers (*REG1A, REG1B, DUOX2, DMBT1*) — all of which returned biological signals consistent with the published analysis (Fig. 5, Step 5). Finally, Step 6 invoked the LIANA skill on a tumor-microenvironment object combining the periphery spots and the tumor spots immediately abutting them, and surfaced the ligand–receptor pairs that differentiate *SPP1*^+^ from *SELENOP*^+^ macrophage signalling at the tumor boundary (Fig. 5, Step 6). The complete reproduction used three named skills (RCTD, gseapy, LIANA) and additional supporting analyses, consumed approximately 7K input tokens of prompts, and took roughly 90 minutes for the deconvolution step and 30 minutes for each subsequent step on a single workstation (Fig. 5, summary panel).

To test how much of this reproduction depends on the recipe-level detail of the full prompt set, we re-ran the same six-step pipeline using a much shorter, conversational prompt set (1.4K input tokens in total) in which each step was specified at the level a senior collaborator would naturally write it — naming regions and macrophage subtypes, but omitting most numerical thresholds and code-level instructions (Supplementary File 2). The agent recovered the same qualitative conclusions: the periphery composition, the *SPP1*^+^/ *SELENOP*^+^ split with the discriminating markers, the divergent pathway programmes, and the boundary cell-cell communication patterns were all reproduced from a prompt set approximately fivefold shorter than the full version (Supplementary Fig. 15). Beyond reproduction, the same dataset also supports analyses that the spot-level publication could not perform at single-cell resolution: because Visium HD is subcellular but not strictly single-cell, we segmented each spot grid into cells with CellART [48] and re-ran cell-level niche detection, marker discovery, and neighbourhood enrichment through STAT, recovering finer-grained spatial structure than the spot-level analysis could expose (Supplementary Fig. 16, Supplementary Fig. 17, Supplementary Fig. 18). Together, these results show that STAT can reproduce a paper-quality ST study end-to-end from natural language prompts and can push such studies into analytical regimes the original publication did not reach, with the prompt budget shrinking roughly fivefold when the user is willing to describe steps at the level they would write on a slide rather than in a script.

## Discussion

The evidence presented here establishes a single coherent claim: that an integrated, interactive, skill-aware architecture changes what an analyst can do with an ST dataset, and that this change is a property of the architecture rather than of any particular language-model backbone. On a benchmark of 40 queries across eleven task categories, three platforms, and both data resolutions, STAT completed 39 against 35–37 for execution-capable competitors and won 31 head-to-head under an anonymised LLM judge; the recurring failures of competing systems — fabricated outputs on infeasible tasks, silent substitution of weaker statistical proxies, under-specification of spatial methods, and ignored prerequisites — each mapped directly to an architectural feature that STAT enforces before code generation. Re-running the same benchmark across seven frontier backbones, together with stage-level tests on a dedicated 90-query pipeline test set, kept end-to-end success above 87% and per-stage accuracy above 86.7%, indicating that this reliability lives in the pipeline rather than in any single model. On a mixed-resolution breast cancer cohort, the same agent absorbed the cell-level versus spot-level distinction without prompting, executed a diverse sequence of downstream analyses through natural language dialogue, and turned a hand-drawn region of interest into a quantitative differential-expression test that recovered the canonical DCIS-versus-invasive transcriptional separation. On a published Visium HD CRC study, STAT reproduced the central characterisation of the *SPP1*^+^ / *SELENOP*^+^ macrophage axis from natural language prompts alone, with a 1.4K-token conversational variant reaching the same biological conclusions as a 7K-token recipe-level version.

Two design philosophies have so far dominated AI agents for biomedical analysis. Autonomous report generators such as Biomni and SpatialAgent execute long tool chains from a single prompt and return a finished report, trading intermediate visibility, token and time efficiency, and mid-flight redirection for minimal user effort. Conversational orchestrators such as ChatSpatial preserve the dialogue but treat the dataset as a text stream rather than a persistent shared object that the user and the agent jointly inspect. STAT represents a third design that takes the visual and iterative character of spatial analysis seriously: the dataset, the tissue viewer, and the dialogue are three views of the same state, and reliability is enforced before code generation rather than retrofitted by self-critique loops afterwards. The practical consequence is a workflow in which the analyst can draw a region on tissue and ask about it in the same sentence, switch between spatial and statistical views without re-running analyses, and audit or re-execute the entire conversation as a notebook.

Several limitations qualify these claims. STAT’s compatibility filter and verifier rely on the metadata declared in each skill file; methods whose applicability declarations are absent or wrong cannot be safely chosen, so the system’s safety is ultimately bounded by the quality of its skill library. The registry is intentionally extensible — a new method requires one Markdown file with a YAML header and an instructional body — but communities will need to maintain it as platforms and tools evolve. Some analyses are not yet covered as skills (RNA velocity, custom deep-learning models on the staining image, bespoke downstream pipelines), and on these the agent collapses to vanilla-LLM behaviour rather than to any of STAT’s reliability guards — a pattern visible in STAT’s only benchmark failure (Q37 RNA velocity, where with no matching skill the code generator called `scVelo` [46] directly without checking for spliced / unspliced layers, reproducing the competitors’ fabrication failure mode). The asymmetry is informative: STAT’s reliability is mediated by registered skills, so growing the registry is the natural way to bring more analyses under the same guards. Because each new method is added as a single Markdown file with a YAML header, the cost of doing so is intentionally low; we view the long-term solution as community curation of the skill library rather than a brittle blanket-refusal policy on out-of-registry queries. The drawn-ROI primitive is also presently 2D and slice-local, so true 3D-tissue or cross-slice region reasoning will require new primitives; and although cross-LLM stability is high, the strongest backbones still set the ceiling on the most ambiguous queries, so further gains will come from both better skill declarations and stronger generalist reasoning. Beyond these immediate limitations, two extensions appear especially natural: pairing STAT’s session with multi-modal spatial assays such as spatial proteomics [49], spatial ATAC [50], or 3D tissue atlases, and porting the same pattern — shared session state, declarative skill applicability, staged selection, classified error reflection — to neighbouring domains such as single-cell genomics, perturbation screens, or multi-omics integration.

As the volume and modality of biological data grow, the rate-limiting step is increasingly not data generation but interpretation. STAT shows that an agent can handle assembly and execution while the analyst retains visual and analytical agency, that an architecture can prevent whole classes of failure rather than retry past them, and that the same object the agent reasons about can be the one the analyst sees and acts on. We hope the integrated, interactive, skill-aware pattern STAT instantiates becomes a template for the next generation of biological analysis tools, in which conversational ease of use does not come at the price of trustworthiness or scientific control.

## Methods

### STAT system architecture

STAT is implemented as a multi-agent system in which a single conversation orchestrator coordinates six specialised roles — a query planner, a programmatic skill filter, a semantic skill matcher, a skill verifier, a code generator with an attached error reflector, and a biological analyzer — that together translate a natural language request into an executed, interpreted analysis (Fig. 1, Fig. 3). All six roles share the same persistent session and the same conversation memory; none of them owns its own copy of the data, and there is no notion of a “current slice” inside the agent — slice identifiers are passed forward explicitly by the planner and read back from session state by every downstream role. Each user turn proceeds along a single deterministic path: the orchestrator first checks whether the message is a response to an outstanding clarification (and routes accordingly), then invokes the planner to determine target slices and decompose multi-step queries, then runs the programmatic compatibility filter against the loaded session, then asks the language model to choose among the surviving compatible skills, then runs the verifier to collect prerequisites, then generates code, executes it in an isolated Python kernel, and finally generates a natural language interpretation. Errors raised during code execution are routed to the error reflector, which classifies them as mechanically fixable or structurally unsupportable and either repairs them or surfaces a human-readable explanation rather than entering an unbounded retry loop. The architecture is intentionally independent of any third-party agent framework (no LangChain [35], LangGraph, or Model Context Protocol [36] dependency); the entire pipeline is implemented in pure Python with a small surface of asynchronous primitives.

### Session and data model

The unit of state in STAT is the `SimpleSession`, an in-memory object that holds an ordered list of `DataSlice` instances together with a session-level `ROIManager`. Each `DataSlice` wraps one AnnData object plus its associated images, and exposes metadata that the rest of the agent can query without re-inspecting the raw data: a unique integer `slice_id`, a `modality` field that takes the values `gene` or `protein`, a `data_level` field that takes the values `cell` or `spot`, the cell or spot count, the list of available annotations, and convenience predicates such as `is_cell_level`, `is_spot_level`, `is_gene`, and `has_celltype()`. AnnData inputs are required to carry per-cell or per-spot coordinates in `adata.obs[‘x’]` and `adata.obs[‘y’]`; `adata.obs[‘celltype’]` is optional and may be added or rewritten by later turns. Spot-level slices additionally carry `adata.uns[‘spot_shape’]` and `adata.uns[‘spot_diameter’]` to drive correct rendering on the canvas, and `adata.obsm[‘deconv_weights’]` once a deconvolution skill has been run. The `ROIManager` records named regions of interest as one of four geometric primitives (axis-aligned bounding box, polygon, free-hand stroke, or circle), each pinned to a specific slice and modality, and supports algebraic combinations (union, intersection, set difference) between named ROIs.

### Skill abstraction and registry

Each analysis method that STAT can invoke is wrapped as a self-describing *skill* stored as a single Markdown file (`SKILL.md`) under `stat_agent/skills/<slug>/`. The file contains a YAML frontmatter that declares the skill’s identifier, a one-paragraph description used by the semantic skill matcher, a `filter_requirements` block that specifies the data conditions under which the skill is applicable (`num_slices`, `modalities`, `data_levels`), an enumerated `prerequisites` list that the verifier reads to collect missing inputs, and a `default_skill` flag. Below the frontmatter, the body of the file is a Markdown document that gives the language model the API surface, the input requirements, the expected outputs, and a worked code template for invoking the underlying library. The registry comprises 27 skills as of the version evaluated in this paper (Supplementary Table 12). Skills are loaded *progressively*: at session startup the registry parses only the YAML frontmatter so that the compatibility filter and matcher can operate on a lightweight catalog; the full Markdown body is loaded on demand only once a skill has been selected for execution. Adding a new method to STAT requires writing one new `SKILL.md` file — no agent code needs to be modified.

### Pipeline stages

### Query planner

The planner receives the raw user message together with a structured summary of the session and the relevant fragment of conversation memory, and returns a JSON object that either decomposes the query into one or more steps with explicit `target_slice_ids` and a `refined_query` per step, or raises a `needs_clarification` flag with a free-text question. The planner resolves explicit slice references, tissue-name references, ROI references, and the multi-target keywords “both”, “all”, “each”, “compare”, and “between”; it falls back to the single available slice when only one is loaded, and asks for clarification when multiple slices match an under-specified request. Full prompt template: Supplementary Note 1, §1.

### Skill filter (programmatic)

The filter is the only deterministic stage in the selection chain. Given the loaded session it computes the tuple `(num_slices_in_request, modalities_present, data_levels_present)` from the planner’s `target_slice_ids` and removes from the candidate pool every skill whose `filter_requirements` are not satisfied. This step is the architectural mechanism that prevents the silent wrong-tool selection failure mode reported for autonomous competitor systems in the benchmark.

### Semantic skill matcher

From the surviving compatible subset, the language model selects up to `top_k = 2` skills whose `description` field specifically matches the user’s request. The matcher is given the compatible candidates as a flat catalog of `slug` →`description` lines and an explicit instruction to be conservative, returning an empty array when no skill is a specific fit. Full prompt template: Supplementary Note 1, §2.

### Skill verifier

The verifier loads the full Markdown body of the selected skill, reads its `prerequisites` list, and decides for each prerequisite whether it is already satisfied by session state, askable from the user via a clarification question, or not obtainable in the current session and requires a prior analytical step. Full prompt template: Supplementary Note 1, §3.

### Code generator

The code generator runs after planning, filtering, matching, and verification have all succeeded. It is invoked with a fixed system prompt containing the session API surface, the standard data conventions (coordinate fields, spot-shape and -diameter for spot data, `obsm[‘deconv_weights’]` for deconvolved spots), ROI access conventions, and a contract on response format (a brief natural language summary, then one or more `python` fenced code blocks, with an optional `INTERPRET :` directive on the last line). Full prompt template: Supplementary Note 1, §4.

### Error reflector

When code execution raises an exception, the reflector receives the offending code, the captured error message, and the available session context, and returns a JSON object that classifies the failure as `small` (mechanically fixable) or `large` (structural / unsupportable). For small fixes the reflector returns the corrected code and the executor retries; for large fixes the reflector returns a human-readable explanation.

The retry budget is bounded at two attempts; the loop terminates immediately if the reflector’s confidence is below 0.7. Full prompt template: Supplementary Note 1, §5.

### Analyzer

After execution the analyzer receives the user’s original query, the executed code, and the captured `stdout` and `stderr` for each code block, together with a compact session snapshot and recent conversation history. It returns a structured JSON object containing a response summary, a description of any data changes, an optional 2–4 sentence biological interpretation (when the code generator requested one via the `INTERPRET :` directive), and an explicit execution-issue verdict that flags errors, validation failures, partial successes, or no-effect runs. Surfacing both the standard output and the error channel to the analyzer is what allows the conversation layer to distinguish a clean run from a silent failure. Full prompt template: Supplementary Note 1, §6.

### Multi-turn clarification

The pipeline distinguishes three classes of clarification: planner clarifications (raised when the planner cannot resolve target slices), verifier clarifications (raised when prerequisites are askable but unavailable), and skill-selection clarifications (raised when the matcher returns more than one specifically-relevant skill). All three are tracked in a `ClarificationContext` object that records the type of clarification, the pending steps, the index of the current step, and the running history of questions and answers.

### Memory and state tracking

STAT keeps two complementary forms of session-level state. The first is a conversation memory that maintains the running dialogue: the most recent eight messages are stored in full and provided verbatim to every LLM-driven role, while older messages are replaced with LLM-generated one-paragraph summaries. Each turn in which code is executed also produces an execution summary that records what the code did at a high level. The second form of state is a data modification history maintained by the agent core: every change to AnnData columns (`obs`, `obsm`, `uns`), every new ROI, and every cell-type or color-mapping update is recorded so the verifier can decide whether prerequisites are met. After each code execution the agent core also runs a `_detect_state_changes` pass that compares before/after snapshots and emits structured events used by the frontend to refresh the canvas.

### LLM backend abstraction

Every LLM-driven role talks to a single asynchronous interface defined by `LLMBackend`, which abstracts over multiple providers (Anthropic, OpenAI, Google, xAI, DeepSeek) behind one method. Token usage and latency are recorded per call. The version of STAT evaluated in this paper uses Claude Sonnet 4 as the backbone for every pipeline role in all main-text experiments; the LLM-as-judge benchmark uses Claude Opus 4.6 as an independent grader. The cross-LLM ablation additionally evaluates Claude Sonnet 4, GPT-5.4, GPT-4o, Grok-3, Grok-4, Gemini 3.1 Pro, and DeepSeek V3.2 as backbones.

### Graphical user interface

The STAT graphical interface is a single-page web application that exposes the session and the multi-agent pipeline through three columns: a skills configuration panel on the left, an interactive tissue canvas in the centre, and an agent chat widget on the right (Fig. 3a). The backend is a Flask server with a small REST surface and one Server-Sent-Events endpoint (`/api/chat/stream`) that streams structured events from the agent to the frontend. The frontend is a vanilla-JavaScript application with a per-canvas caching layer for instant slice and modality switching. The coordinate convention is direct: a cell at `(x, y)` in `adata.obs` is rendered at image pixel `(x, y)`. ROI drawing is performed in canvas space and translated to data coordinates before being persisted into the session. Three rendering modes are exposed on the canvas — cell-type overlays, single-gene expression heatmaps, and deconvolution-proportion overlays (Supplementary Fig. 11, Supplementary Fig. 12, Supplementary Fig. 13). For multi-slice cohorts a 3D viewer renders the slices as stacked layers (Supplementary Fig. 9). Each session can be exported as a Markdown analysis history and as an executable Jupyter notebook.

### Datasets

### Human DLPFC Visium cohort

The four-slice human dorsolateral prefrontal cortex Visium dataset [43] was obtained from the spatialLIBD project (http://spatial.libd.org/spatialLIBD/), with manually curated cortical-layer annotations (L1–L6 plus white matter) provided per spot.

### Single-cell reference for DLPFC deconvolution

The matched human prefrontal cortex single-cell reference profiled on the 10x Genomics Chromium platform was downloaded from the Gene Expression Omnibus accession GSE144136 and used as the cell-type reference for RCTD deconvolution of the Visium spots.

### MERFISH whole-mouse-brain dataset

In-situ spatially resolved transcriptomic profiles of individual cells from the whole mouse brain generated by multiplexed error-robust fluorescence in situ hybridisation (MERFISH) using a 1,122-gene panel [44] (Zhuang lab; Zhang et al. 2023). The full collection spans four animals (two coronal sets, two sagittal sets) covering 245 coronal and sagittal sections; 9.3M segmented cells passed quality control and were integrated with the Allen Institute scRNA-seq atlas, of which 5.8M cells additionally passed cell-cluster label-transfer confidence and are displayed on the Allen Brain Cell (ABC) Atlas. Spatial coordinates are aligned to the Allen Common Coordinate Framework (Allen-CCF-2020) and cell-type labels follow the whole-mouse-brain (WMB) taxonomy. In this work we used the two coronal specimens, Zhuang-ABCA-1 and Zhuang-ABCA-3, as the two MERFISH benchmark slices.

### Xenium human breast cancer dataset

The 10x Genomics Xenium In Situ human breast cancer preview dataset (https://www.10xgenomics.com/products/xenium-in-situ/preview-dataset-human-breast), cell-resolution, 160,000 cells across a 313-gene panel (306 protein-coding plus controls). The single-slice benchmark of Fig. 2 uses replicate 1 only.

### Mixed-resolution Xenium + Visium breast cancer cohort (Fig. 4)

Three breast cancer slices loaded into one STAT session for the interactive workflow demonstration: the two Xenium cell-resolution replicates from the preview dataset above (replicate 1 — the same slice used in Fig. 2 — and replicate 2, both profiled on adjacent sections of the same tumor block) and a paired Visium spot-level slice from an adjacent section, each accompanied by its H&E image.

### Visium HD colorectal cancer cohort (Fig. 5)

Three Visium HD slices from the 10x Genomics Visium HD human colorectal cancer dataset family (https://www.10xgenomics.com/platforms/visium/product-family/dataset-human-crc patients P1, P2, P5; processed at 8 µm bin size), as released and characterised in Oliveira et al. (Nat. Genet. 2025) [34]. The matched single-cell reference used by RCTD was the one published with the original study.

### Benchmark protocol

#### Forty-query benchmark (Fig. 2)

40 queries across 11 task categories (cell-type annotation, spatial-domain detection, SVG, cell-cell communication (CCC), neighborhood enrichment, deconvolution, niche, differential expression (DE), pathway, batch integration, exploratory) and four difficulty regimes (clear, vague, multi-step-task, special boundary cases). Full query list: Supplementary Table 3. Four systems (vanilla LLM, Biomni, SpatialAgent, STAT) run on the same backbone (Claude Sonnet 4); 160 runs total. Per-run output and per-run cost (tokens, wall time, LLM API calls) captured.

#### LLM-as-judge evaluation

Each run scored by Claude Opus 4.6 as anonymised expert judge for binary success, integer quality (1–5), and per-query rank against task-specific success criteria fixed in advance (Supplementary Table 4). Failure-mode analysis: Supplementary Table 5.

#### Cross-LLM ablation (Fig. 3d)

Same 40-query benchmark re-run with seven backbones (Claude Sonnet 4, GPT-5.4, GPT-4o, Grok-3, Grok-4, Gemini 3.1 Pro, DeepSeek V3.2). Per-backbone failures: Supplementary Table 6, Supplementary Fig. 10.

#### Per-stage pipeline ablation (Fig. 3e)

Dedicated 90-query pipeline test set (30 each for planner, skill filter+matcher, refined-query) on six mock sessions (Supplementary Tables 7–10). Sets A and B used deterministic ground-truth comparison; Set C used Claude Sonnet 4 as a programmatic judge (Supplementary Table 10). Per-failure analysis: Supplementary Table 11.

### Per-task methodology

#### Cell-type deconvolution

RCTD [7] on raw UMI counts; alternatives Cell2location [8] and FlashDeconv exposed as separate skills.

#### Cell-type annotation

Default scANVI [9] (scvi-tools); alternatives CellTypist [10] (logistic-regression label transfer against pretrained or user-supplied references), Tangram [11], and clustering+LLM marker annotation.

#### Spatial-domain identification

Default SpaGCN [4]; alternatives STAGATE [5] and GraphST [6].

#### Spatially variable genes

SpatialDE [16] for spot-level; Moran’s I via squidpy [42] for cell-level.

#### Niche detection

Harmonics [19] on the celltype neighbourhood graph.

#### Cell-cell communication

Default LIANA+ [13]; alternative CellPhoneDB [14] for human-only.

#### Differential expression and pathway enrichment

scanpy [41] `rank_genes_groups` Wilcoxon for DE; gseapy [51] for GO/Hallmark/KEGG/Reactome enrichment with Fisher’s exact + BH FDR; single-sample gene set enrichment analysis (ssGSEA) via gseapy for per-cell pathway scoring.

#### Batch integration

Harmony [21], BBKNN [22], Scanorama [23] exposed as separate skills.

#### Spatial alignment and registration

STalign [24] (LDDMM); PASTE [25] (optimal transport).

#### Trajectory inference

Palantir [52] and scanpy diffusion pseudotime (DPT) for pseudotime.

#### Hardware

All Mac-side analyses (sessions, GUI, most benchmark runs) executed on an Apple M1 Max workstation with 32 GB unified memory. GPU-dependent methods (scANVI, Cell2location, Tangram training) executed on a Linux compute node with two NVIDIA Tesla V100 GPUs and an Intel Xeon Gold 6152 CPU at 2.10 GHz.

## Software, code, and data availability

STAT is implemented in pure Python and distributed as the PyPI package `stat-agent` (`pip install stat-agent`). The agent source code, including the skill registry, is available at https://github.com/YangLabHKUST/STAT-agent under an open-source license, and a live web demo is hosted at https://huggingface.co/spaces/CyhVVVV/stat-agent-demo. All datasets used in this work are available from public repositories. The Visium DLPFC cohort is available at spatial.libd.org/spatialLIBD; the matched human prefrontal cortex single-cell reference at GEO accession GSE144136; the MERFISH whole-mouse-brain dataset from the Allen Institute Whole Mouse Brain Cell Atlas (portal.brain-map.org/atlases-and-data/ bkp/abc-atlas); the Xenium human breast cancer preview dataset from 10xgenomics.com/products/xenium-in-situ/preview-dataset-human-breast; and the Visium HD colorectal cancer cohort from the 10x Genomics Visium HD dataset family (10xgenomics.com/platforms/visium/product-family/dataset-human-crc; patients P1, P2, P5; processed at 8 µm bin size), as released and characterised in [34]. The mixed-resolution breast cancer cohort of Fig. 4 combines the Xenium replicates above with a paired Visium spot-level slice from an adjacent section, available from the same Xenium preview release.

## Acknowledgements

This work was supported in part by the Innovation and Technology Commission (ITCPD/17-9); Hong Kong Research Grants Council Grants, AoE/E-601/24-N, C6040-24G, 16308120, 16307221, 16307322, 16302823, 16309424, and 16308925; The Hong Kong University of Science and Technology Startup Grants R9405 and Z0428 from the Big Data Institute. The computation tasks for this work were performed using the X-GPU cluster supported by the Research Grants Council Collaborative Research Fund Grant C6021-19EF and HKUST SuperPOD. J.X. was supported by National Natural Science Foundation of China (Grant No. 12401384); Guangdong Natural Science Foundation General Project (Grant No. 2025A1515011603); Sun Yat-sen University Startup Grant (Grant No. 2026_51000_B26833); and Shenzhen Science and Technology Program (Grant No. RCBS20231211090613024).

## Author Contributions

C. Y. conceived the study and supervised the project. Y. C. designed, implemented, and validated STAT. J. X. and Y. L. contributed to the experimental design. S. H., Z. C., F. Z., H. C., and J. W. helped with analyzing the results. Y. C. and C. Y. wrote the manuscript with input from all the authors.

## References

[1] L. Moses and L. Pachter, “Museum of spatial transcriptomics,” Nature Methods, vol. 19, pp. 534–546, 2022, doi: 10.1038/s41592-022-01409-2.

[2] X. Wan et al., “Integrating spatial and single-cell transcriptomics data using deep generative models with SpatialScope,” Nature Communications, vol. 14, no. 1, p. 7848, 2023.

[3] V. Marx, “Method of the Year: spatially resolved transcriptomics,” Nature Methods, vol. 18, pp. 9–14, 2021, doi: 10.1038/s41592-020-01033-y.

[4] J. Hu et al., “SpaGCN: Integrating gene expression, spatial location and histology to identify spatial domains and spatially variable genes by graph convolutional network,” Nature Methods, vol. 18, pp. 1342–1351, 2021, doi: 10.1038/s41592-021-01255-8.

[5] K. Dong and S. Zhang, “Deciphering spatial domains from spatially resolved transcriptomics with an adaptive graph attention auto-encoder,” Nature Communications, vol. 13, p. 1739, 2022, doi: 10.1038/s41467-022-29439-6.

[6] Y. Long et al., “Spatially informed clustering, integration, and deconvolution of spatial transcriptomics with GraphST,” Nature Communications, vol. 14, p. 1155, 2023, doi: 10.1038/s41467-023-36796-3.

[7] D. M. Cable et al., “Robust decomposition of cell type mixtures in spatial transcriptomics,” Nature Biotechnology, vol. 40, pp. 517–526, 2022, doi: 10.1038/s41587-021-00830-w.

[8] V. Kleshchevnikov et al., “Cell2location maps fine-grained cell types in spatial transcriptomics,” Nature Biotechnology, vol. 40, pp. 661–671, 2022, doi: 10.1038/s41587-021-01139-4.

[9] C. Xu, R. Lopez, E. Mehlman, J. Regier, M. I. Jordan, and N. Yosef, “Probabilistic harmonization and annotation of single-cell transcriptomics data with deep generative models,” Molecular Systems Biology, vol. 17, no. 1, p. e9620, 2021, doi: 10.15252/msb.20209620.

[10] C. Domínguez Conde et al., “Cross-tissue immune cell analysis reveals tissue-specific features in humans,” Science, vol. 376, no. 6594, p. eabl5197, 2022.

[11] T. Biancalani et al., “Deep learning and alignment of spatially resolved single-cell transcriptomes with Tangram,” Nature Methods, vol. 18, pp. 1352–1362, 2021, doi: 10.1038/s41592-021-01264-7.

[12] G. Wang, J. Zhao, Y. Yan, Y. Wang, A. R. Wu, and C. Yang, “Construction of a 3D whole organism spatial atlas by joint modelling of multiple slices with deep neural networks,” Nature Machine Intelligence, vol. 5, no. 11, pp. 1200–1213, 2023.

[13] D. Dimitrov et al., “LIANA+ provides an all-in-one framework for cell-cell communication inference,” Nature Cell Biology, vol. 26, pp. 1571–1582, 2024, doi: 10.1038/s41556-024-01469-w.

[14] M. Efremova, M. Vento-Tormo, S. A. Teichmann, and R. Vento-Tormo, “CellPhoneDB: inferring cell-cell communication from combined expression of multi-subunit ligand-receptor complexes,” Nature Protocols, vol. 15, pp. 1484–1506, 2020, doi: 10.1038/s41596-020-0292-x.

[15] S. Jin et al., “Inference and analysis of cell-cell communication using CellChat,” Nature communications, vol. 12, no. 1, p. 1088, 2021.

[16] V. Svensson, S. A. Teichmann, and O. Stegle, “SpatialDE: identification of spatially variable genes,” Nature Methods, vol. 15, pp. 343–346, 2018, doi: 10.1038/nmeth.4636.

[17] Z. Wang et al., “A unified framework for identification of cell-type-specific spatially variable genes in spatial transcriptomic studies,” Proceedings of the National Academy of Sciences, vol. 122, no. 46, p. e2503952122, 2025.

[18] L. Shang, P. Wu, and X. Zhou, “Statistical identification of cell type-specific spatially variable genes in spatial transcriptomics,” Nature communications, vol. 16, no. 1, p. 1059, 2025.

[19] Y. Liu et al., “Hierarchical distribution matching enables comprehensive characterization of common and condition-specific cell niches in spatial omics data,” Under review, 2025.

[20] X. Yu et al., “NicheScope: Identifying Multicellular Niches and Niche-Regulated Cell States in Spatial Transcriptomics,” bioRxiv, pp. 2025–2028, 2025.

[21] I. Korsunsky et al., “Fast, sensitive and accurate integration of single-cell data with Harmony,” Nature Methods, vol. 16, pp. 1289–1296, 2019, doi: 10.1038/s41592-019-0619-0.

[22] K. Polański, M. D. Young, Z. Miao, K. B. Meyer, S. A. Teichmann, and J.-E. Park, “BBKNN: fast batch alignment of single cell transcriptomes,” Bioinformatics, vol. 36, no. 3, pp. 964–965, 2020, doi: 10.1093/bioinformatics/btz625.

[23] B. Hie, B. Bryson, and B. Berger, “Efficient integration of heterogeneous single-cell transcriptomes using Scanorama,” Nature Biotechnology, vol. 37, pp. 685–691, 2019, doi: 10.1038/s41587-019-0113-3.

[24] K. Clifton et al., “STalign: Alignment of spatial transcriptomics data using diffeomorphic metric mapping,” Nature Communications, vol. 14, p. 8123, 2023, doi: 10.1038/s41467-023-43915-7.

[25] R. Zeira, M. Land, A. Strzalkowski, and B. J. Raphael, “Alignment and integration of spatial transcriptomics data,” Nature Methods, vol. 19, pp. 567–575, 2022, doi: 10.1038/s41592-022-01459-6.

[26] K. Huang et al., “Biomni: A General-Purpose Biomedical AI Agent,” bioRxiv, 2025, doi: 10.1101/2025.05.30.656746.

[27] Y. Xiao et al., “CellAgent: An LLM-driven Multi-Agent Framework for Automated Single-cell Data Analysis,” arXiv preprint, 2024.

[28] S. Alber, B. Chen, E. Sun, A. Isakova, A. J. Wilk, and J. Zou, “CellVoyager: AI CompBio agent generates new insights by autonomously analyzing biological data,” Nature Methods, 2026, doi: 10.1038/s41592-026-03029-6.

[29] Y. Mao, Y. Mi, P. Liu, M. Zhang, H. Liu, and Y. Gao, “scAgent: Universal Single-Cell Annotation via a LLM Agent,” arXiv preprint, 2025.

[30] C. Yang, X. Zhang, and J. Chen, “ChatSpatial: Schema-Enforced Agentic Orchestration for Reproducible and Cross-Platform Spatial Transcriptomics,” bioRxiv, 2026, doi: 10.64898/2026.02.26.708361.

[31] H. Wang et al., “SpatialAgent: An autonomous AI agent for spatial biology,” bioRxiv, 2025, doi: 10.1101/2025.04.03.646459.

[32] Z. Lin et al., “Spatial transcriptomics AI agent charts hPSC-pancreas maturation in vivo,” bioRxiv, 2025, doi: 10.1101/2025.04.01.646731.

[33] A. Janesick et al., “High resolution mapping of the tumor microenvironment using integrated single-cell, spatial and in situ analysis of FFPE tissue,” Nature Communications, vol. 14, p. 8353, 2023, doi: 10.1038/s41467-023-43458-x.

[34] M. F. d. Oliveira et al., “High-definition spatial transcriptomic profiling of immune cell populations in colorectal cancer,” Nature Genetics, vol. 57, no. 6, pp. 1512–1523, 2025.

[35] Lang Chain, “LangChain: Building applications with LLMs through composability.” [Online]. Available: https://github.com/langchain-ai/langchain

[36] Anthropic, “Model Context Protocol: An open standard for connecting AI assistants to data sources and tools.” [Online]. Available: https://modelcontextprotocol.io/

[37] P.L. Ståhl et al., “Visualization and analysis of gene expression in tissue sections by spatial transcriptomics,” Science, vol. 353, no. 6294, pp. 78–82, 2016, doi: 10.1126/science.aaf2403.

[38] A. Janesick et al., “High resolution mapping of the tumor microenvironment using integrated single-cell, spatial and in situ analysis of FFPE tissue,” Nature Communications, vol. 14, p. 8353, 2023, doi: 10.1038/s41467-023-43458-x.

[39] K. H. Chen, A. N. Boettiger, J. R. Moffitt, S. Wang, and X. Zhuang, “Spatially resolved, highly multiplexed RNA profiling in single cells,” Science, vol. 348, no. 6233, p. aaa6090, 2015, doi: 10.1126/science.aaa6090.

[40] A. Chen et al., “Spatiotemporal transcriptomic atlas of mouse organogenesis using DNA nanoball-patterned arrays,” Cell, vol. 185, no. 10, pp. 1777–1792.e21, 2022, doi: 10.1016/j.cell.2022.04.003.

[41] F. A. Wolf, P. Angerer, and F. J. Theis, “SCANPY: large-scale single-cell gene expression data analysis,” Genome Biology, vol. 19, p. 15, 2018, doi: 10.1186/s13059-017-1382-0.

[42] G. Palla et al., “Squidpy: a scalable framework for spatial omics analysis,” Nature Methods, vol. 19, pp. 171–178, 2022, doi: 10.1038/s41592-021-01358-2.

[43] K. R. Maynard et al., “Transcriptome-scale spatial gene expression in the human dorsolateral prefrontal cortex,” Nature Neuroscience, vol. 24, pp. 425–436, 2021, doi: 10.1038/s41593-020-00787-0.

[44] Z. Yao et al., “A high-resolution transcriptomic and spatial atlas of cell types in the whole mouse brain,” Nature, vol. 624, pp. 317–332, 2023, doi: 10.1038/s41586-023-06812-z.

[45] P. A. P. Moran, “Notes on continuous stochastic phenomena,” Biometrika, vol. 37, no. 1/2, pp. 17–23, 1950, doi: 10.2307/2332142.

[46] V. Bergen, M. Lange, S. Peidli, F. A. Wolf, and F. J. Theis, “Generalizing RNA velocity to transient cell states through dynamical modeling,” Nature Biotechnology, vol. 38, no. 12, pp. 1408–1414, 2020.

[47] DeepSeek-AI, “DeepSeek-V3 Technical Report,” arXiv preprint, 2024, doi: 10.48550/arXiv.2412.19437.

[48] Y. Chen et al., “CellART: a unified framework for cell segmentation and annotation in high-resolution spatial transcriptomics,” Manuscript under review, 2026.

[49] Y. Goltsev et al., “Deep profiling of mouse splenic architecture with CODEX multiplexed imaging,” Cell, vol. 174, no. 4, pp. 968–981.e15, 2018, doi: 10.1016/j.cell.2018.07.010.

[50] Y. Deng et al., “Spatial profiling of chromatin accessibility in mouse and human tissues,” Nature, vol. 609, pp. 375–383, 2022, doi: 10.1038/s41586-022-05094-1.

[51] Z. Fang, X. Liu, and G. Peltz, “GSEApy: a comprehensive package for performing gene set enrichment analysis in Python,” Bioinformatics, vol. 39, no. 1, p. btac757, 2023, doi: 10.1093/bioinformatics/btac757.

[52] M. Setty, V. Kiseliovas, J. Levine, A. Gayoso, L. Mazutis, and D. Pe’er, “Characterization of cell fate probabilities in single-cell data with Palantir,” Nature Biotechnology, vol. 37, pp. 451–460, 2019, doi: 10.1038/s41587-019-0068-4.

