## Supplementary for "STAT: A multi-agent framework for integrated and interactive spatial transcriptomics analysis"

#### Supplementary Contents


Supplementary Figures ........ S2
Supplementary Tables ........ S18
Supplementary Note 1 — STAT pipeline prompt templates ........ S45
Supplementary File 1 — CRC analytic prompts, full (recipe-level) version ........ S51
Supplementary File 2 — CRC analytic prompts, conversational (slide-level) version ........ S60

### Supplementary Figures

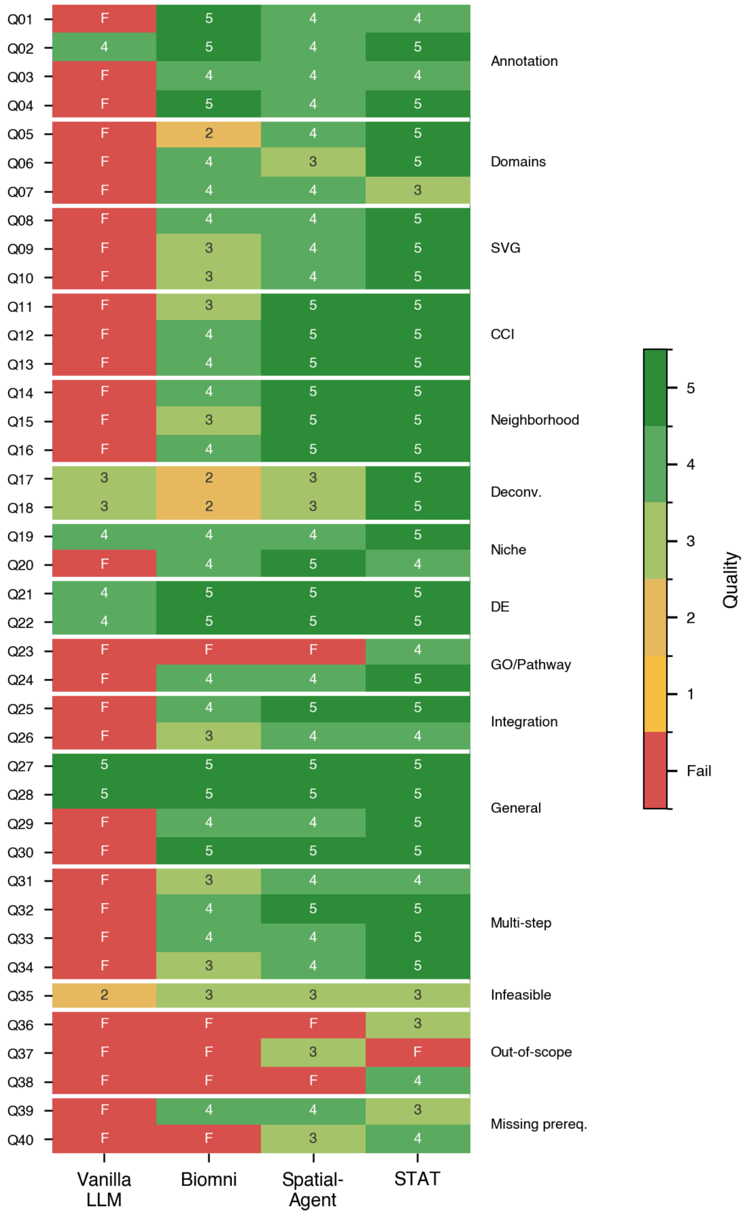


**Supplementary Figure 1. Per-query benchmark results across all four systems.** Per-query heatmap of judge-assigned outcomes for each of the 40 benchmark queries × 4 systems (Vanilla LLM, Biomni, SpatialAgent, STAT). Cells are coloured by quality score (1–5; green high, orange low, red fail "F"); rows are grouped on the right by task category (Annotation, Domains, SVG, CCI, Neighborhood, Deconv., Niche, DE, GO/Pathway, Integration, General, Multi-step, Infeasible, Out-of-scope, Missing prereq.). STAT shows the densest distribution of green cells across all categories, and the only failure for STAT is on the out-of-scope RNA-velocity query (Q37) — the same query that defeated all three competitors. This per-query view is the underlying breakdown of the aggregate metrics summarised in Fig. 2d.

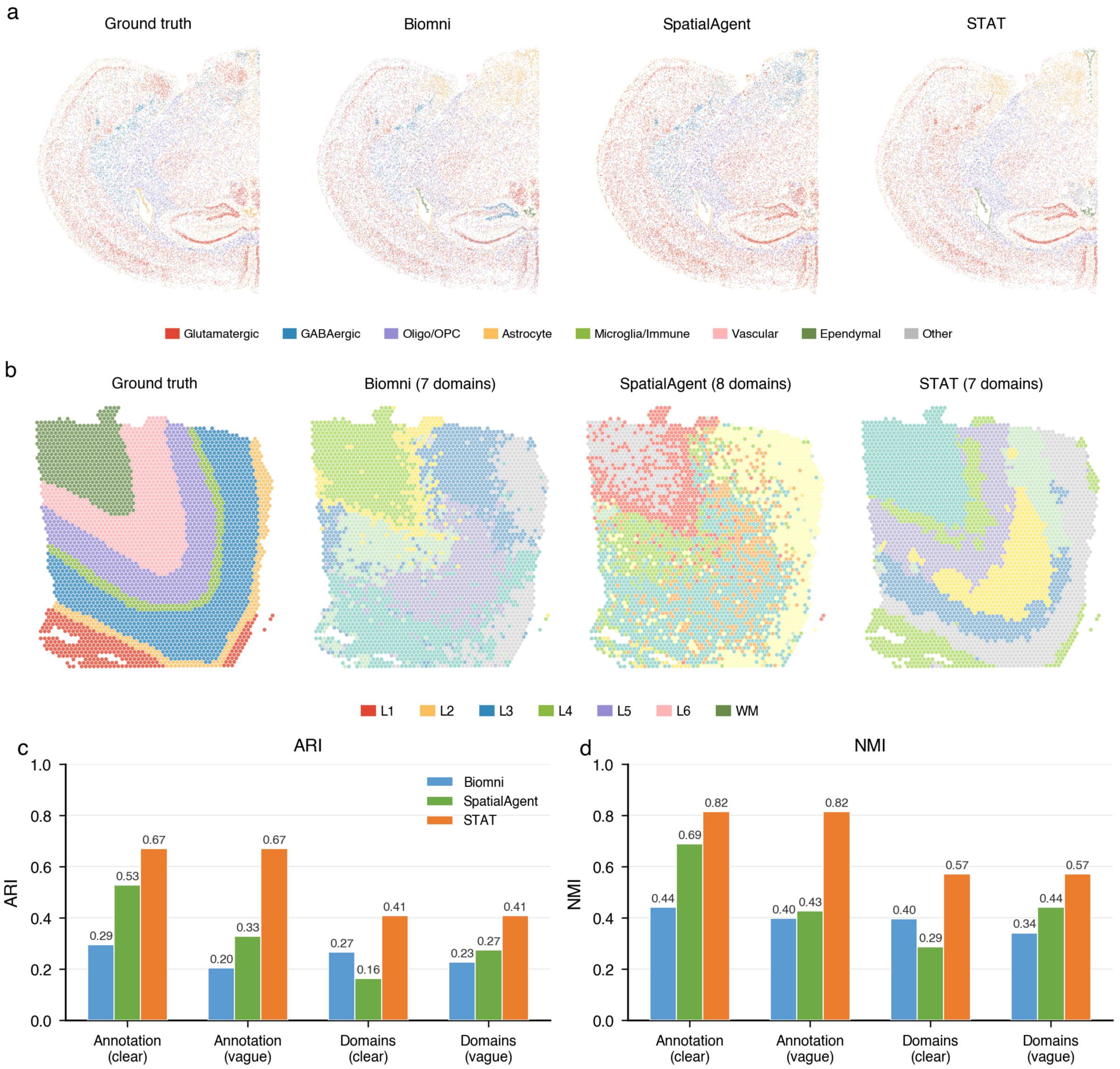


**Supplementary Figure 2. Ground-truth label agreement on cell-type annotation (MERFISH) and spatial-domain identification (DLPFC). a,** Cell-type annotation on a MERFISH mouse-brain slice — the curated ground-truth labels (left) and the per-spot label maps recovered by Biomni, SpatialAgent, and STAT, coloured by the seven canonical cell-type families (Glutamatergic, GABAergic, Oligo/OPC, Astrocyte, Microglia/Immune, Vascular, Ependymal; plus “Other”). **b,** Spatial-domain identification on a DLPFC slice — the manually curated cortical-layer ground truth (left) and the domain maps recovered by each system, coloured by the seven canonical layers L1–L6 plus white matter (WM); the “(N domains)” annotation in each panel header reports the number of clusters returned by that system. **c,** Adjusted Rand Index (ARI) between the recovered labels and the ground truth on four query variants — Annotation (clear) Q01, Annotation (vague) Q03, Domains (clear) Q05, Domains (vague) Q06 — for Biomni, SpatialAgent, and STAT. **d,** Normalised Mutual Information (NMI) on the same four query variants. STAT achieves the highest agreement on every panel of **c** and **d**, consistent with the LLM-judge ranking summarised in Fig. 2d.

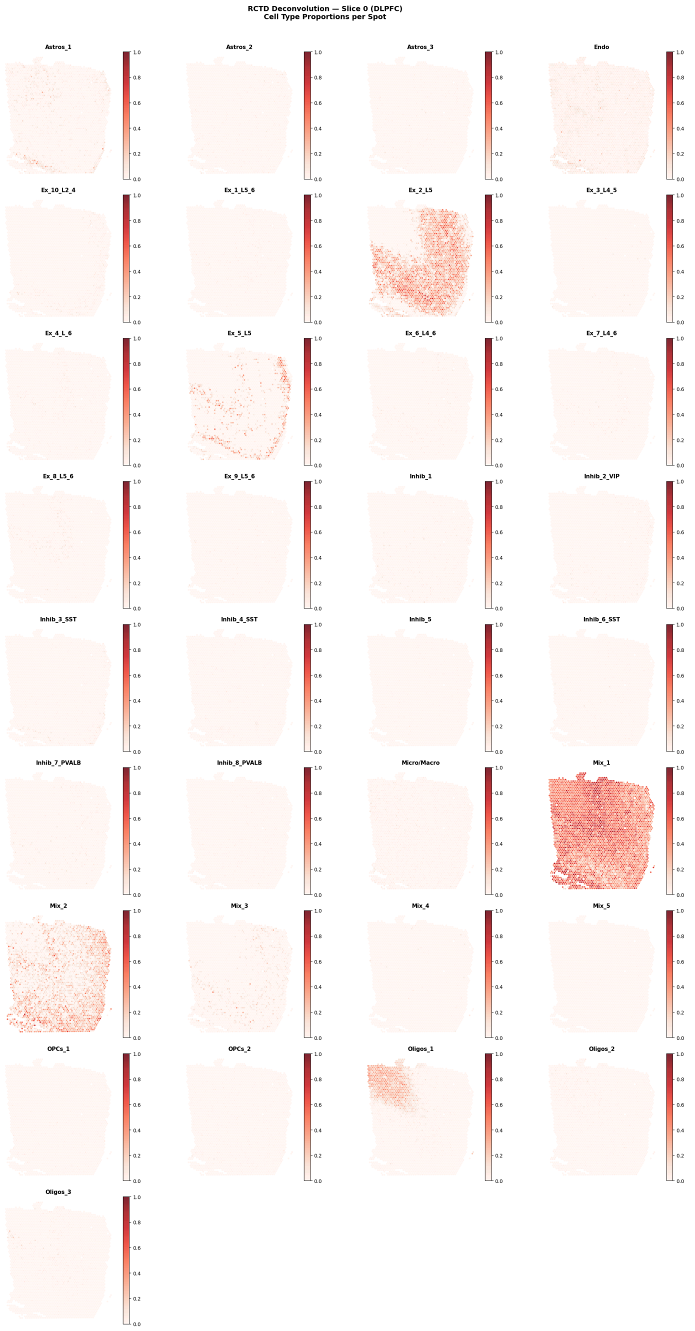


**Supplementary Figure 3. RCTD spot-level cell-type deconvolution on the DLPFC Visium cohort.** Per-spot proportion maps for each cell-type column returned by RCTD on a representative DLPFC slice (slice 0), after STAT's skill matcher selected the `celltype-deconvolution` (RCTD) skill against the matched human prefrontal-cortex single-cell reference (GSE144136). Each subplot is one cell-type subcluster from the reference (Astro, Ex layer-specific subtypes, Inh subtypes, Micro/Macro, Oligo, OPC, and additional reference subclusters), with colour intensity encoding the per-spot proportion of that cell type. The expected layered pattern of upper-cortical excitatory subtypes towards the pia and deep-layer subtypes towards white matter is recovered, and the rare populations (microglia, OPC) show diffuse low-level distributions consistent with their biology rather than collapsing onto single dominant spots.

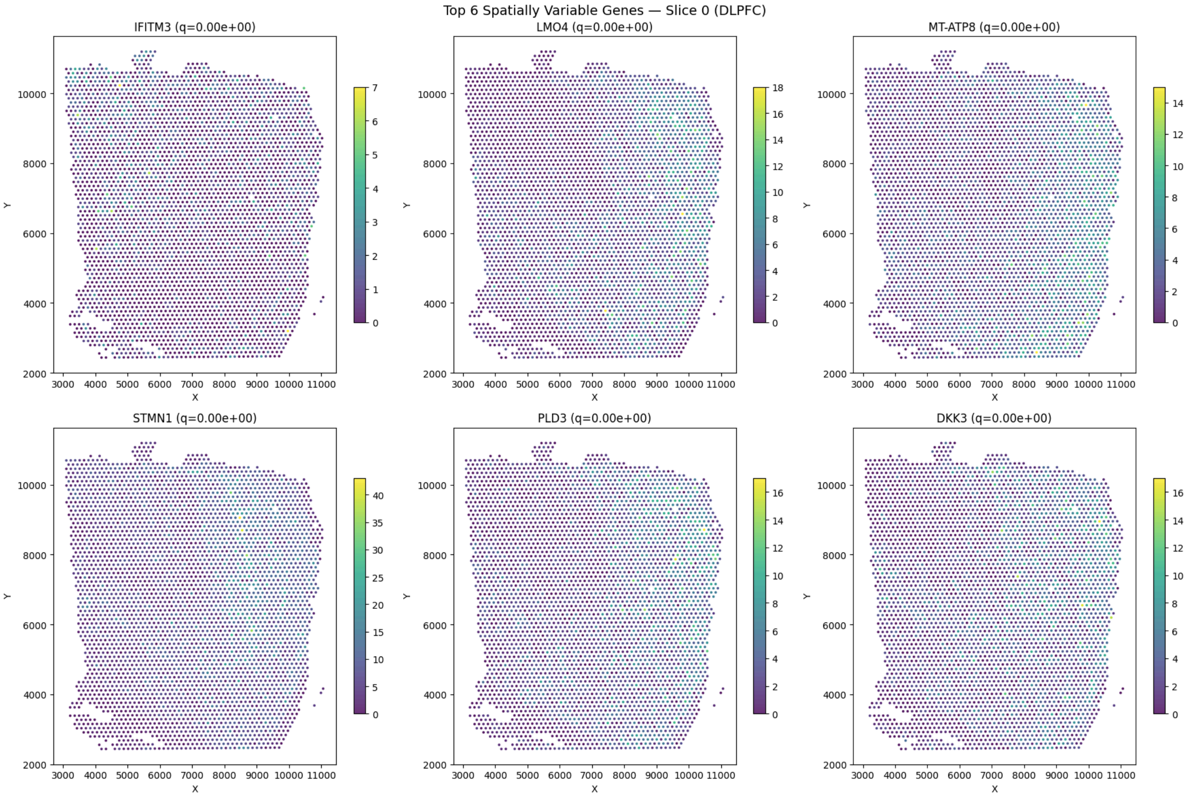


**Supplementary Figure 4. Top spatially variable genes on the DLPFC Visium cohort, identified by SpatialDE.** The six top-ranked spatially variable genes returned by STAT after the `svg-spatialde` skill was selected on the spot-level DLPFC data: *IFITM3*, *LMO4*, *MT-ATP8*, *STMN1*, *PLD3*, and *DKK3* (all $q \approx 0$ under the SpatialDE Gaussian-process likelihood ratio). Each panel shows the per-spot expression on slice 0; the recovered patterns track the cortical-layer organisation of the tissue and exemplify the laminar markers expected for human DLPFC. STAT's skill matcher selects SpatialDE specifically for spot-level Visium data, where the Gaussian-process likelihood is the appropriate statistical model.

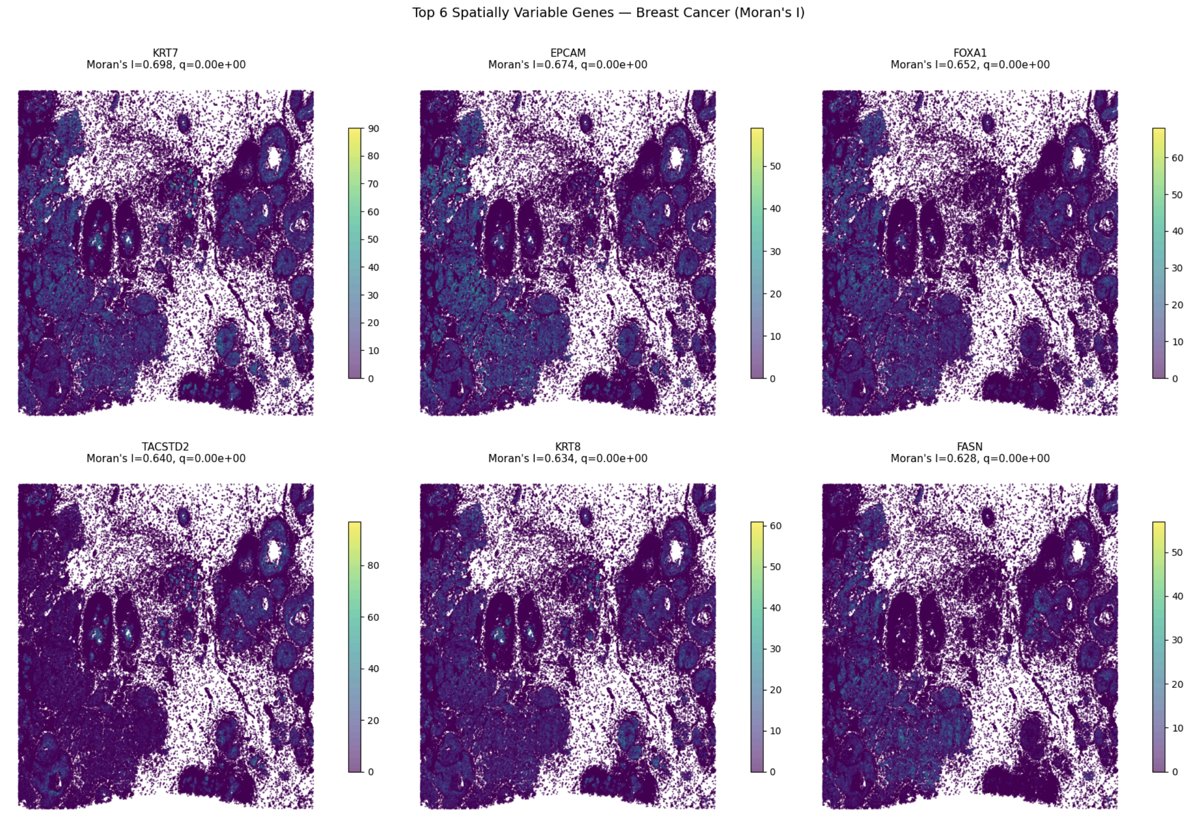


**Supplementary Figure 5. Top spatially variable genes on the Xenium breast cancer slice, identified by Moran's I.** The six top-ranked spatially variable genes returned by STAT after the `spatial-statistics` skill was selected on the cell-resolution Xenium breast cancer data: *KRT7* (Moran's I = 0.698), *EPCAM* (0.674), *FOXA1* (0.652), *TACSTD2* (0.640), *KRT8* (0.634), and *FASN* (0.628), all with permutation q ≈ 0. Each panel maps per-cell expression of one gene over the tissue; the recovered autocorrelation patterns trace the tumour-epithelial compartments distinct from the surrounding stroma. On cell-resolution data the skill matcher selects Moran's I (squidpy permutation framework) rather than the Gaussian-process SpatialDE used for spot-level data — illustrating the data-resolution-aware method choice that is encoded in the skill `filter_requirements`.

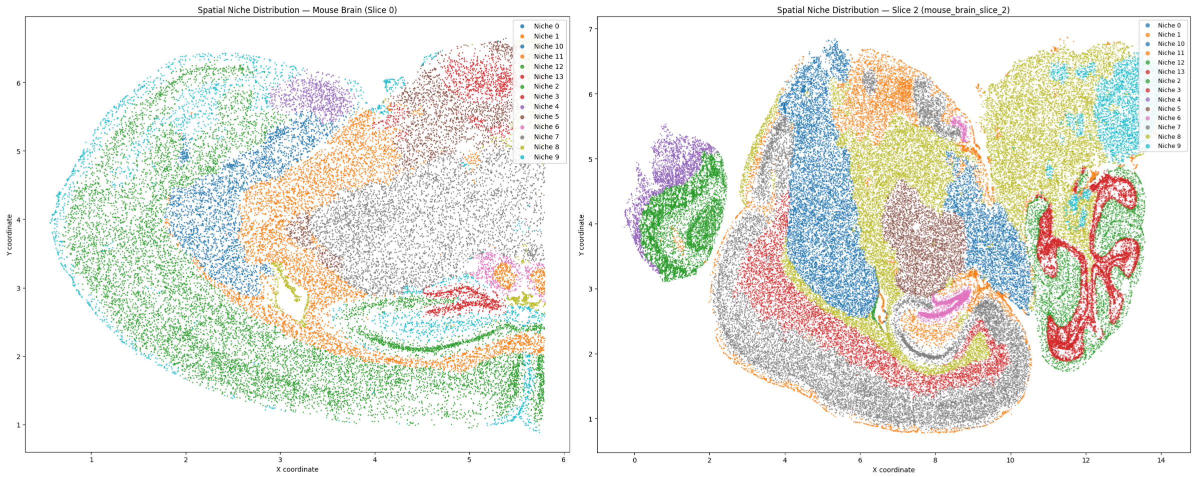


**Supplementary Figure 6. Niche detection on the two-slice MERFISH mouse-brain dataset.** Spatial niche distribution returned by STAT after the `niche-detection` (Harmonics) skill was applied to each MERFISH slice — slice 0 (left) and slice 2 (right) — using the curated cell-type labels as input. Each cell is coloured by its assigned niche (14 niches, legend on right). The recovered niches trace the classical anatomical and functional organisation of the mouse brain (cortical layers, hippocampus, thalamus, hypothalamus, fibre tracts) without the user specifying anatomical priors, illustrating that the same skill recovers consistent partitions across two independent slices of the same tissue.

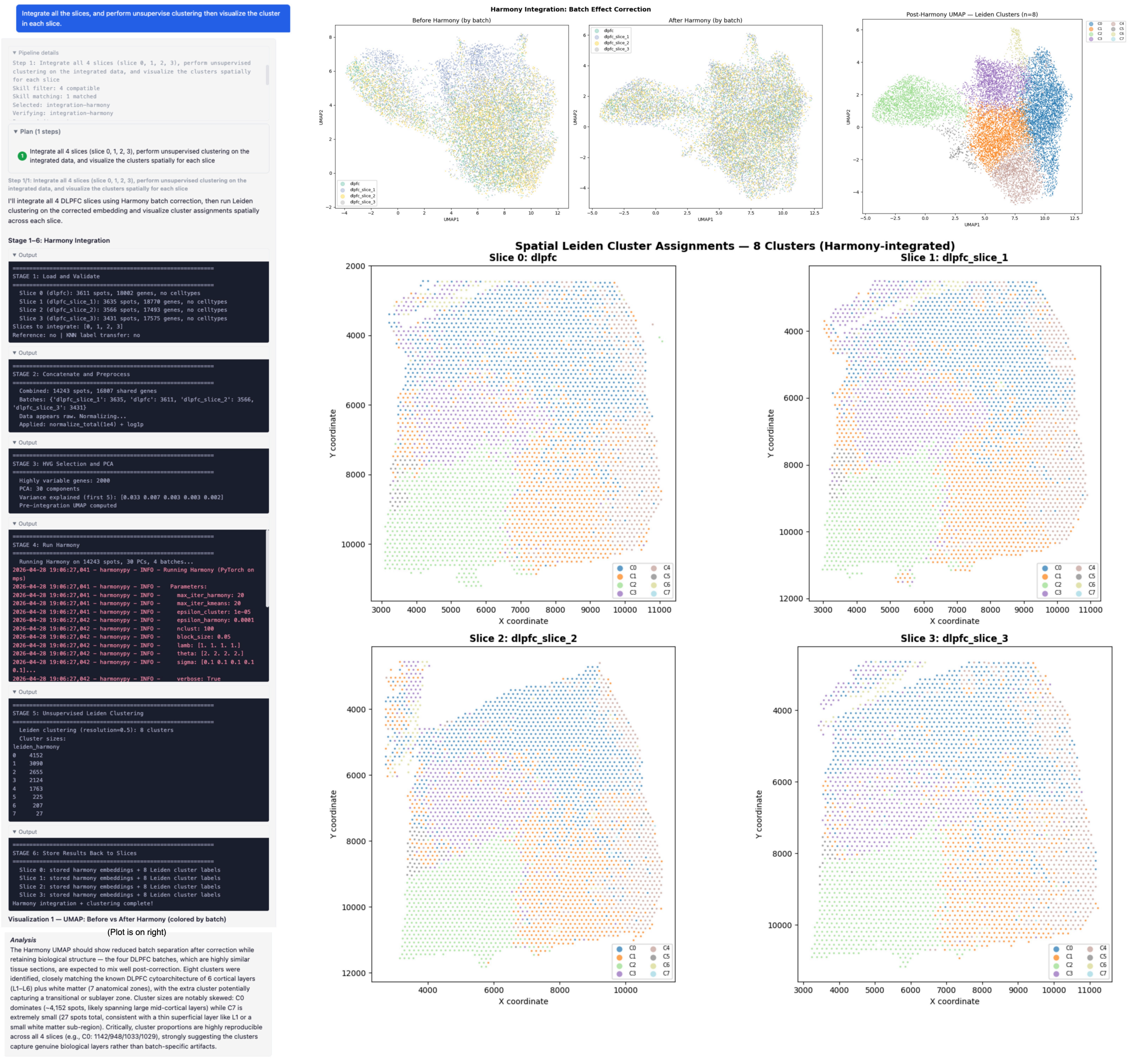


**Supplementary Figure 7. Harmony integration of the four-slice DLPFC Visium cohort, with downstream Leiden clustering and spatial mapping.** The conversational trace (left) shows STAT being asked to integrate the four DLPFC slices and then to cluster from the integrated representation; the pipeline routes the request through Harmony for batch correction followed by Leiden on the corrected embedding. UMAPs (top right) compare the joint embedding before and after Harmony correction (coloured by batch) and the post-correction UMAP coloured by the eight recovered Leiden clusters; the four slices are well mixed in the corrected space. The bottom panel maps the eight Harmony-integrated Leiden clusters back onto each of the four tissue slices, showing that the recovered clusters trace coherent spatial domains across the cohort.

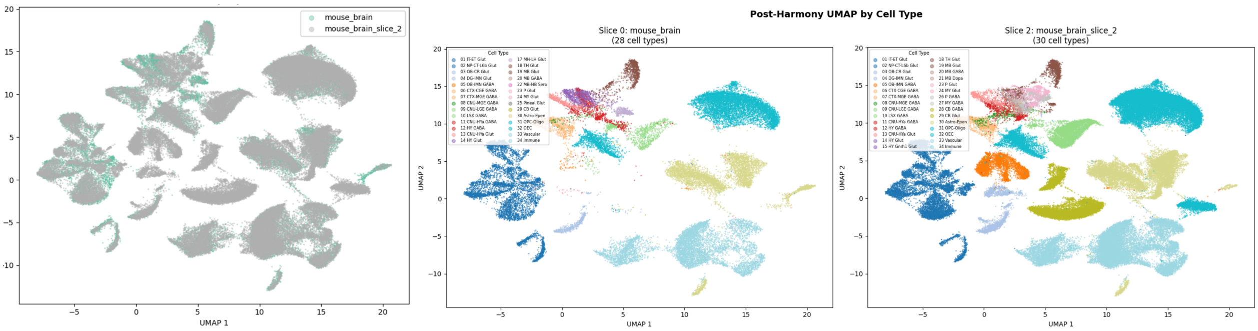


**Supplementary Figure 8. Harmony batch integration of the two-slice MERFISH mouse-brain dataset.** Post-Harmony joint UMAP returned by STAT after the `integration-harmony` skill was applied to the two MERFISH brain slices, coloured three ways: by source slice (left, both slices in grey to highlight overall mixing), by cell type within slice 0 (centre, 28 cell-type categories), and by cell type within slice 2 (right, same legend). After correction the two slices share one embedding manifold and the matched cell types co-localise across slices; one residual cluster stays slice-specific because the corresponding cell type is present only in the second slice — a biologically expected separation rather than a failure of the correction.

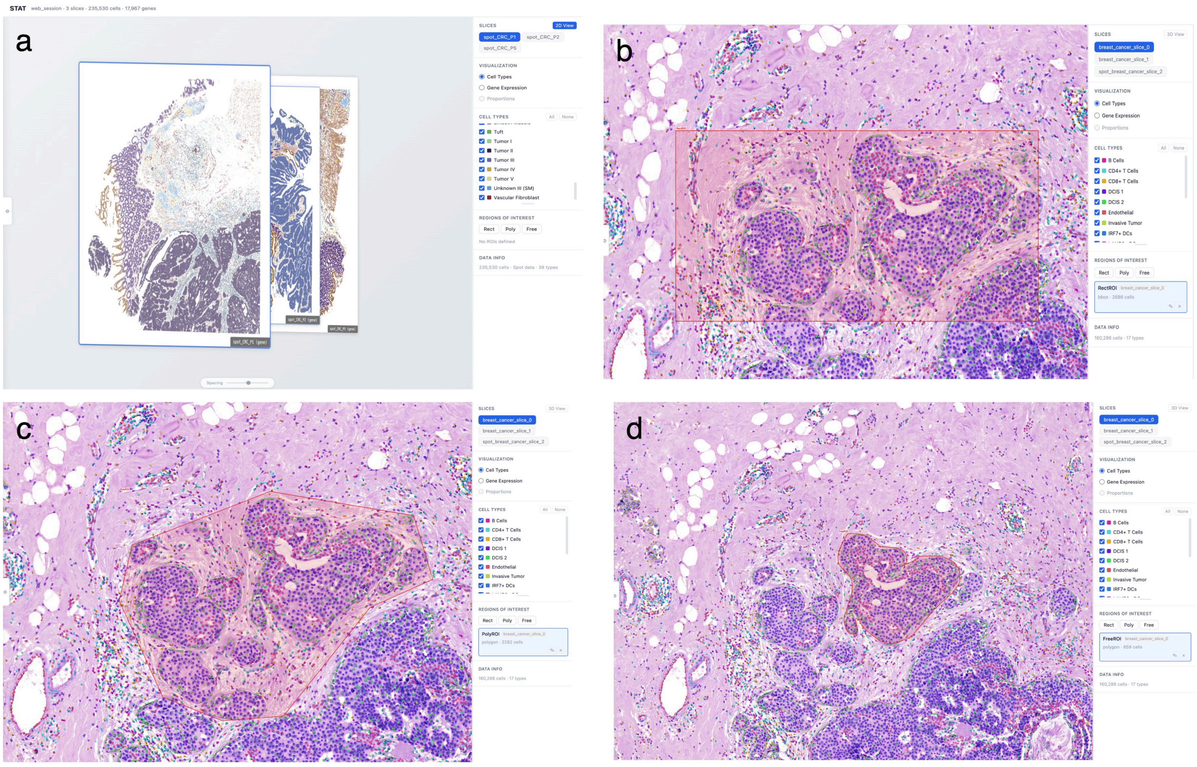


**Supplementary Figure 9. 3D viewer and ROI-driven region inspection on a multi-slice breast cancer cohort. a,** The 3D viewer renders the multi-slice cohort as stacked tissue layers in a single canvas, allowing the user to inspect cross-slice anatomy at a glance. **b,** On a single slice, an axis-aligned rectangle ROI (red dashed box) is drawn directly over a region of interest on the H&E view. **c,** The same ROI displayed with the cell-type overlay enabled, showing the cell-type composition of cells inside the ROI together with the surrounding tissue context. **d,** The same ROI displayed with the deconvolution-proportion overlay enabled, switching the visualization mode without re-running any analysis. The right-hand control panel exposes ROI tools (Rect, Poly, Free) and a list of named ROIs that the agent can subsequently address by name in natural-language queries.

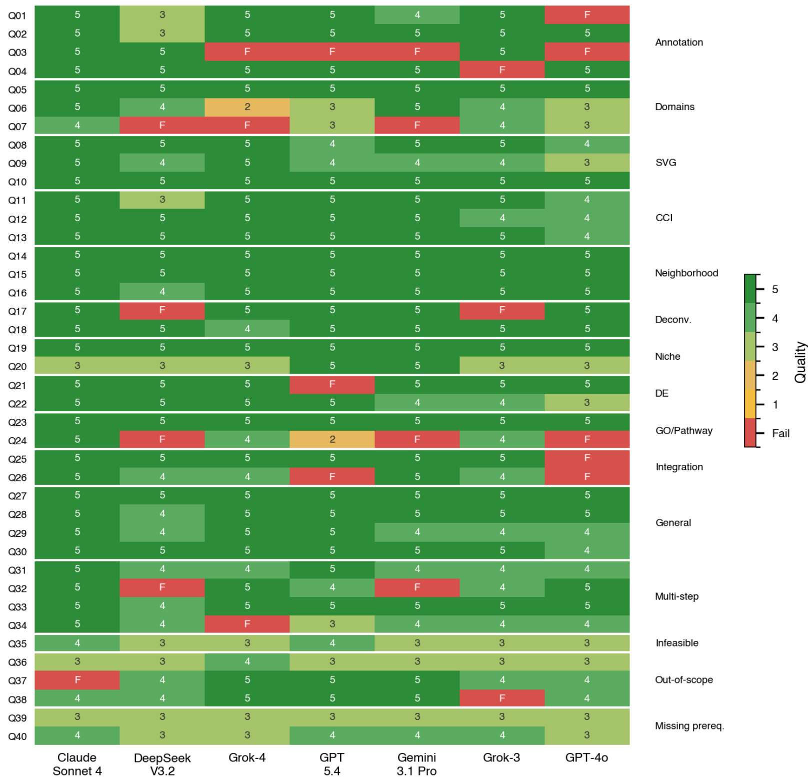

**Supplementary Figure 10. Per-LLM × per-query results on the 40-query benchmark for the seven-backbone ablation.** Per-query heatmap of judge-assigned outcomes when the same 40-query benchmark was re-run with each of seven frontier backbones in turn — Claude Sonnet 4, DeepSeek V3.2, Grok-4, GPT-5.4, Gemini 3.1 Pro, Grok-3, and GPT-4o (columns) — holding the STAT pipeline architecture, skill registry, and LLM-judge protocol identical to the main benchmark. Cells are coloured by quality score (1–5; green high, orange low, red fail "F"), with rows grouped on the right by task category. Failures are sparse and concentrated on out-of-scope and missing-prerequisite queries plus a small number of model-specific multi-step decompositions; the bulk of every column is green, supporting the architectural-stability claim of Fig. 3d and Supplementary Table 6.

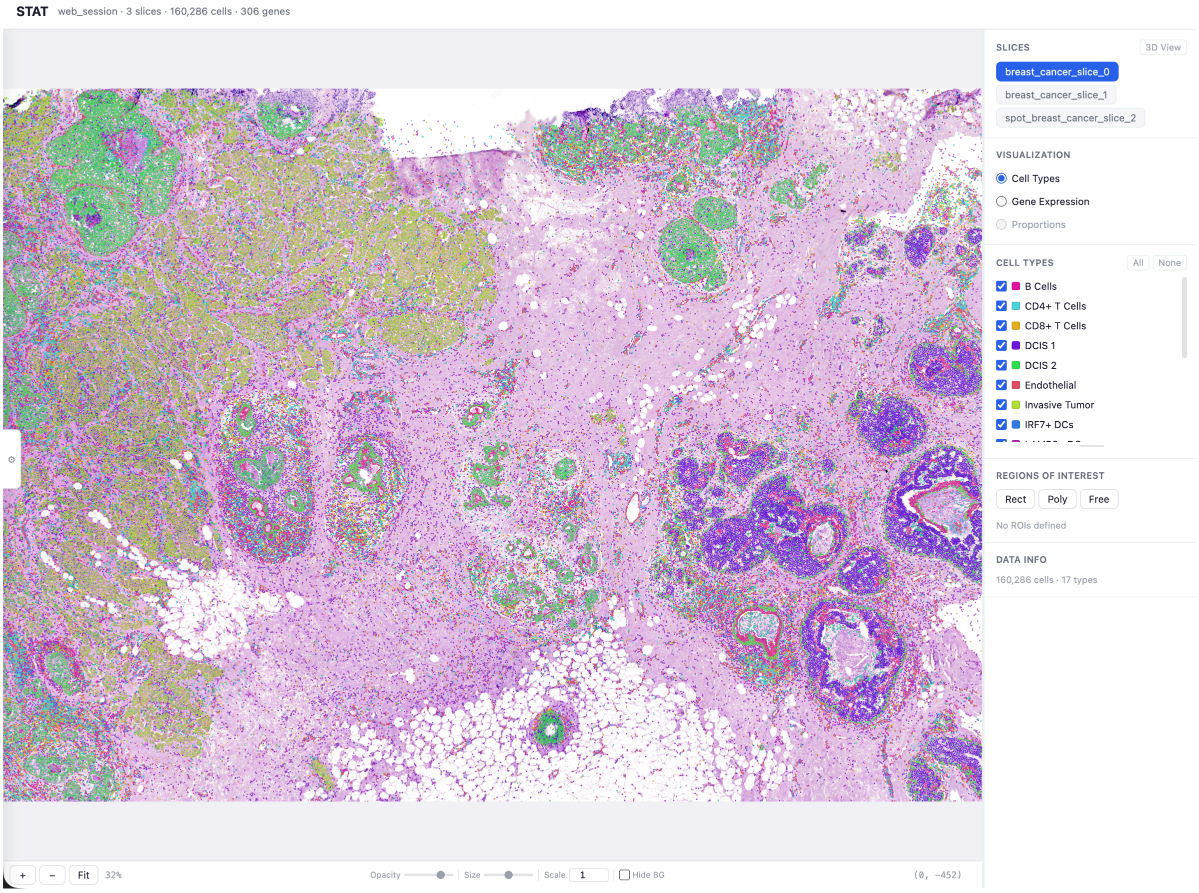


**Supplementary Figure 11. Cell-type rendering mode of the STAT interactive tissue canvas.** Screenshot of the STAT web interface (Xenium breast cancer session, 3 slices, 160,286 cells, 306 genes) with the **Cell Types** visualization mode selected on the right-hand control panel. The canvas renders each cell as a coloured dot over the H&E image, with the colour determined by the cell-type column of the active slice; the cell-type checkboxes (right) selectively show or hide each population, the slice selector (top right) switches the active slice without losing zoom/pan state, and ROI tools (Rect, Poly, Free) below allow a region to be drawn directly on tissue and added to the named-ROI list for subsequent agent queries. The same canvas is shown in single-gene mode in Supplementary Fig. 12 and in deconvolution-proportion mode in Supplementary Fig. 13.

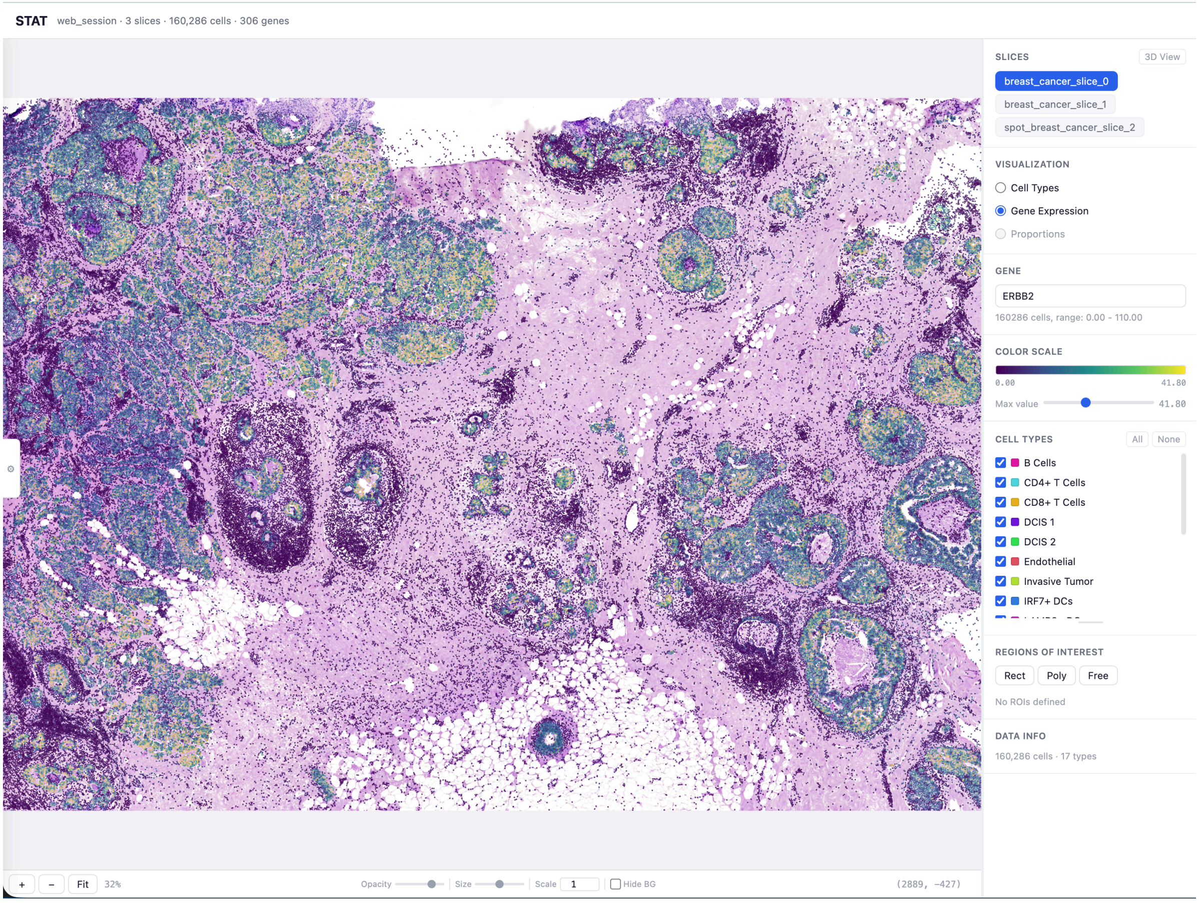


**Supplementary Figure 12. Single-gene rendering mode of the STAT interactive tissue canvas.** The same Xenium breast cancer session shown in Supplementary Fig. 11, with the **Gene Expression** visualization mode selected and *ERBB2* entered in the gene field. The canvas now colours each cell by its *ERBB2* expression on a perceptually uniform viridis-style scale (range 0.00–193.00 across the slice; max-value cap adjustable via the slider on the right), highlighting the *ERBB2*-positive tumour-epithelial regions of the tissue while preserving cell positions and ROI drawings. Switching between modes does not re-run the underlying analyses — the same per-cell state is re-projected.

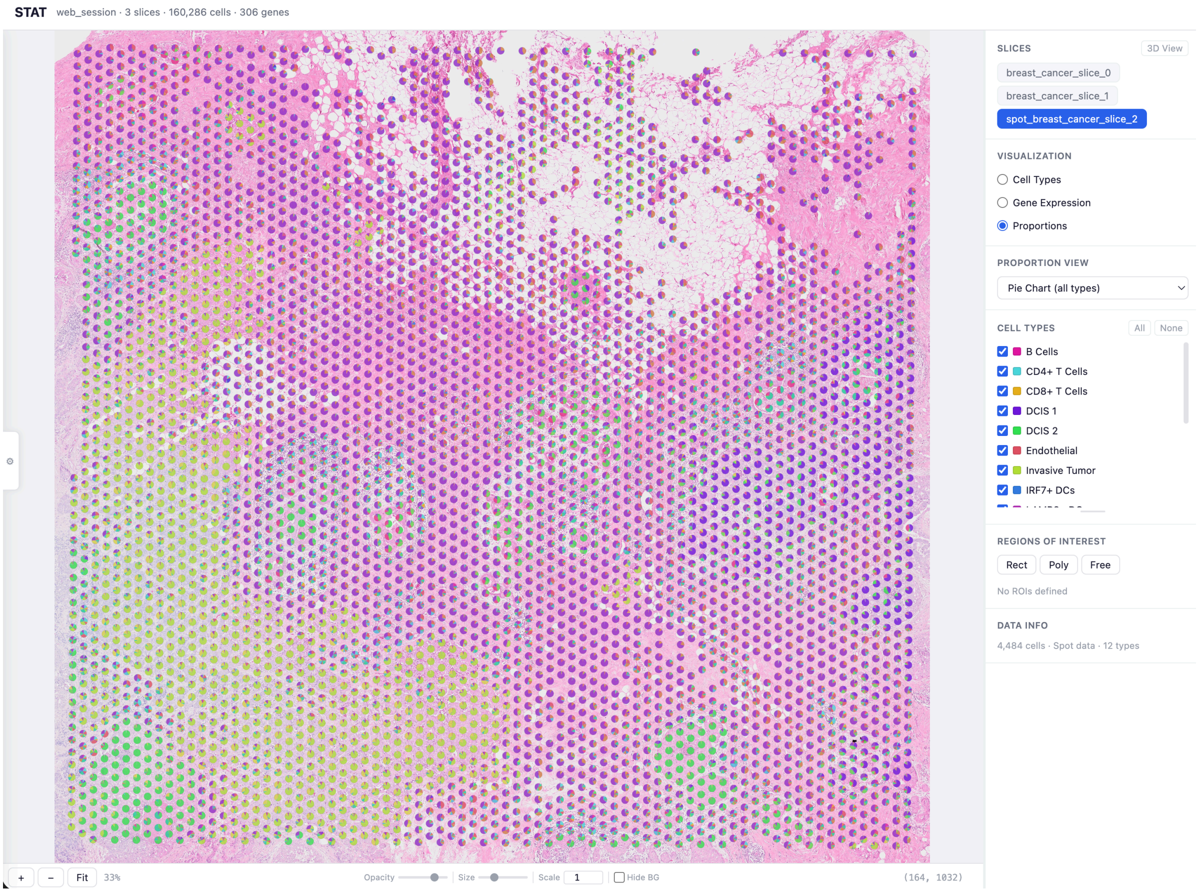


**Supplementary Figure 13. Deconvolution-proportion rendering mode of the STAT interactive tissue canvas.** Same session as Supplementary Fig. 11 and Supplementary Fig. 12 with the spot-level Visium slice (`spot_breast_cancer_slice_2`, 4,994 spots, 12 cell types) selected in the slice list and the **Proportions** visualization mode active. Each spot is rendered as a pie chart whose wedges encode the per-cell-type proportions returned by the RCTD deconvolution skill (legend on right; "Pie Chart (all types)" view is selected). Mode-switching here is again a re-projection of the existing `obsm['deconv_weights']`; the deconvolution itself is not re-run.

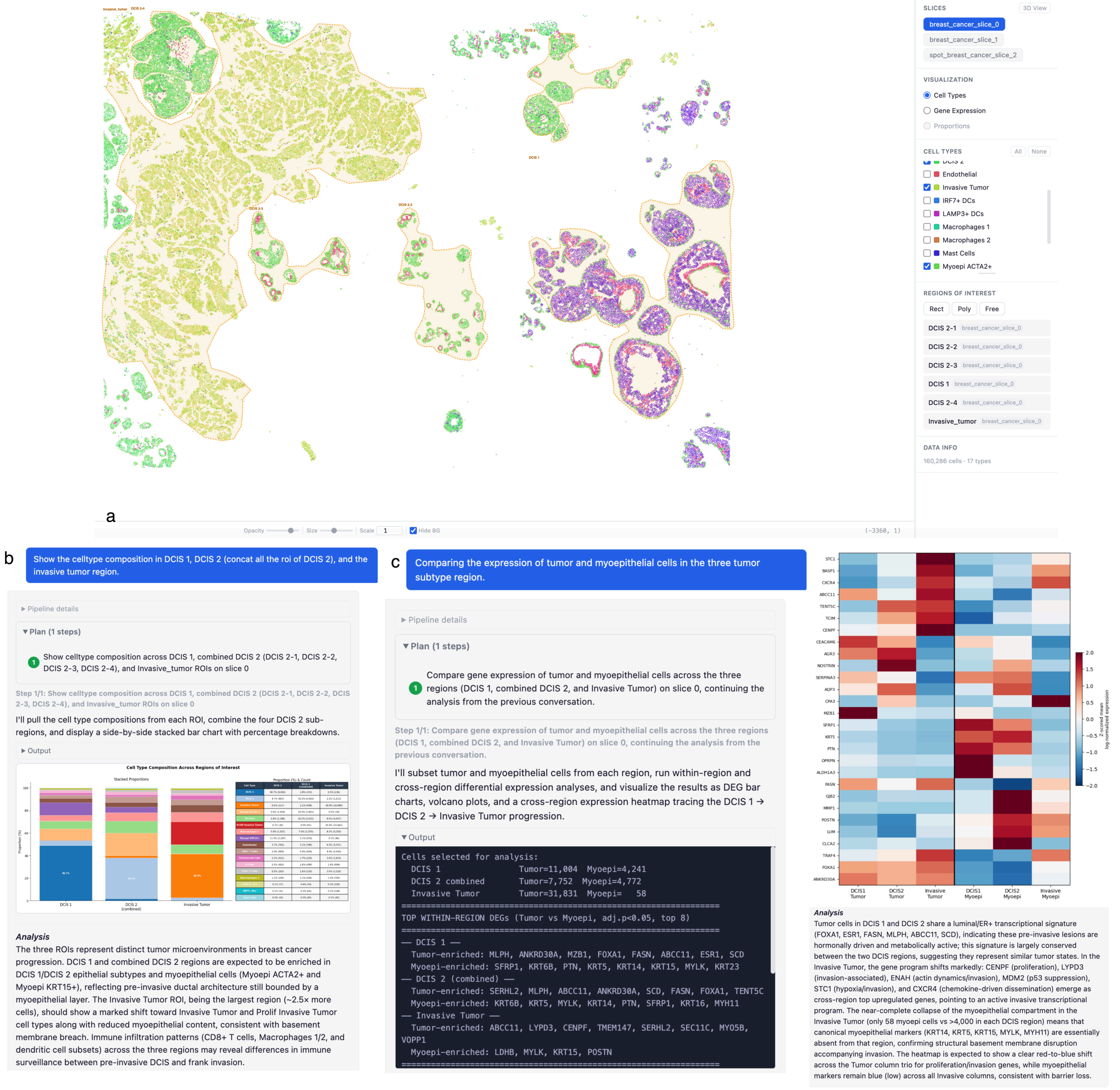


**Supplementary Figure 14. Three-ROI workflow on the breast cancer cohort. a,** Three ROIs drawn directly on the canvas over invasive-tumor, DCIS 1, and DCIS 2 enriched regions of an annotated Xenium slice. **b,** The natural-language query "Show celltype composition in DCIS 1, DCIS 2, combined DCIS 1 + DCIS 2, and the invasive tumor ROIs" returns a side-by-side stacked-bar breakdown of cell-type proportions within each region. **c,** The follow-up query "Compare the expression of tumor and myoepithelial cells in the three tumor subtype regions" runs a within-region differential test and renders the discriminating tumor and myoepithelial markers as a per-region heatmap, recovering the canonical DCIS 1 → DCIS 2 → Invasive Tumor progression and the accompanying myoepithelial breakdown.

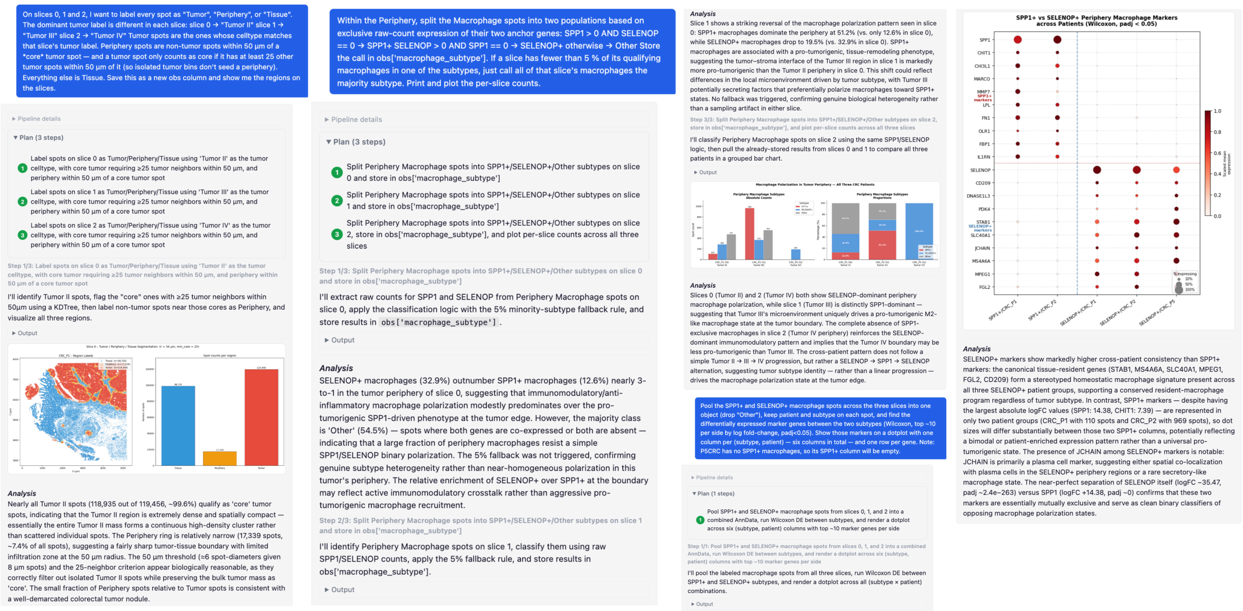


**Supplementary Figure 15. CRC reproduction with the short conversational prompt set ( 1.4K input tokens total) — illustrative first three steps.** Conversational trace produced by re-running the same six-step pipeline of Fig. 5 on the three Visium HD CRC slices using the slide-level prompts of Supplementary File 2 (each step specified at the level a senior collaborator would naturally write — naming regions and macrophage subtypes, but omitting numerical thresholds and code-level instructions). Three representative steps of the conversational trace are shown. **Left,** periphery identification: STAT assigns each spot to Tumor, Periphery, or Tissue and returns the per-region cell-type composition, recapitulating the immune enrichment at the periphery. **Centre,** *SPP1*$^+$ / *SELENOP*$^+$ macrophage subtyping within the periphery: macrophage spots are split into the two subtypes from raw-count expression of the anchor genes *SPP1* and *SELENOP* (spots co-expressing both are pooled into a separate "Both" category and excluded from the downstream discriminating-marker analysis). **Right,** per-patient discriminating markers: a Wilcoxon test between the two pooled macrophage populations rendered as a per-patient dot plot, which recovers the macrophage marker programmes reported by the original publication. The remaining steps of the simple-prompt reproduction (two-group hallmark-pathway enrichment, niche-restricted differential expression, boundary cell–cell communication) are reproduced equivalently to the full-prompt run of Fig. 5 but are omitted from this figure for compactness; the same qualitative conclusions are recovered from a prompt budget roughly two orders of magnitude smaller than the full prompt set.

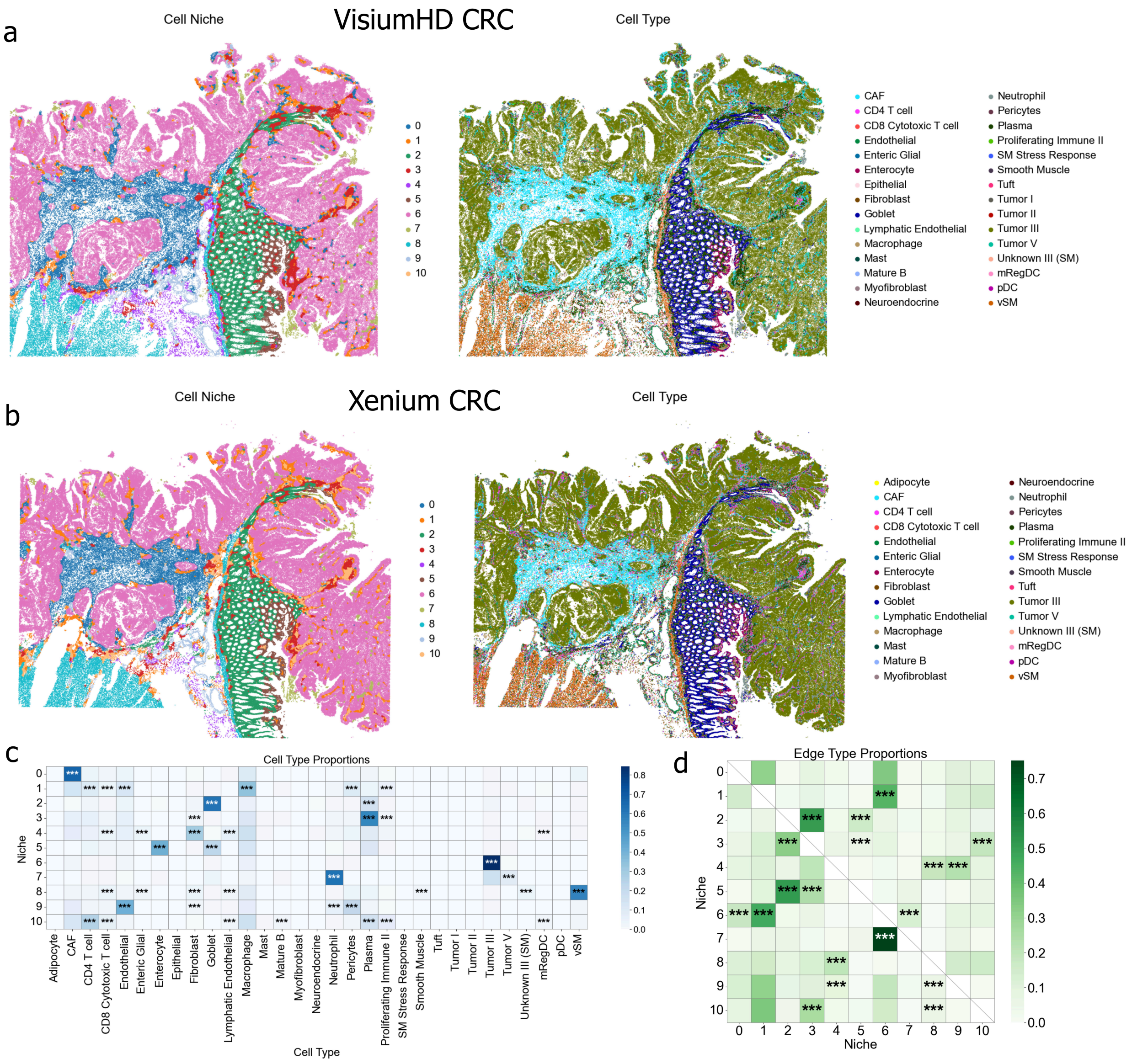


**Supplementary Figure 16. Cell-level niche detection on the CRC cohort across two spatial platforms. a,** Visium HD CRC tissue segmented into cells with CellART, with cells coloured by recovered niche (left, 11 niches 0–10) and by cell type (right, shared cell-type legend). **b,** Matched Xenium CRC tissue analysed through the same STAT skill, again coloured by niche (left) and by cell type (right). **c,** Cell-type proportion matrix per niche (rows: niche 0–10; columns: cell types) computed by STAT, with significance asterisks marking enrichment of each cell type in each niche; the matrix recovers tumor-, periphery-, and tissue-side niches that align with their dominant cell-type compositions. **d,** Inter-niche edge-type proportion matrix (rows × columns: niches 0–10), again with significance asterisks; off-diagonal enrichments mark spatial adjacencies between niches that recapitulate the tumor-periphery–tissue layering. Compared with the spot-level analysis of the original publication, the cell-level run recovers finer-grained spatial structure and shows that the same niche organisation is reproduced across the two cell-resolution platforms.

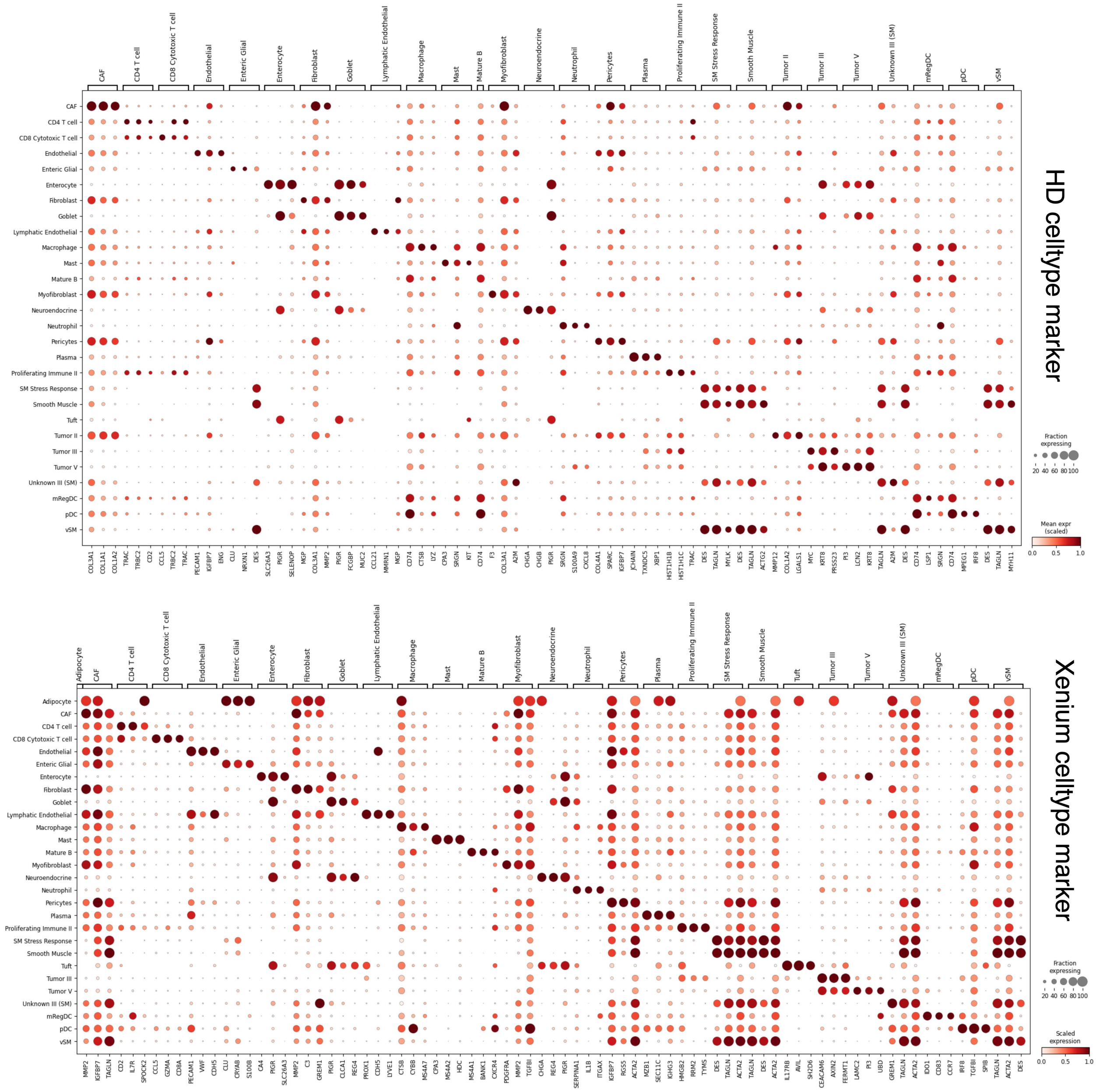


**Supplementary Figure 17. Cell-level marker discovery on the CRC cohort across two spatial platforms.** Per-cell-type marker dot plots returned by STAT after CellART cell segmentation. **Top,** Visium HD CRC slices segmented into cells and annotated through STAT, with marker genes shown across the recovered cell populations (dot size: fraction of cells expressing; colour: scaled mean expression). **Bottom,** matched Xenium CRC slices analysed through the same STAT skill on the cell-resolution data, showing platform-consistent cell-type marker programmes.

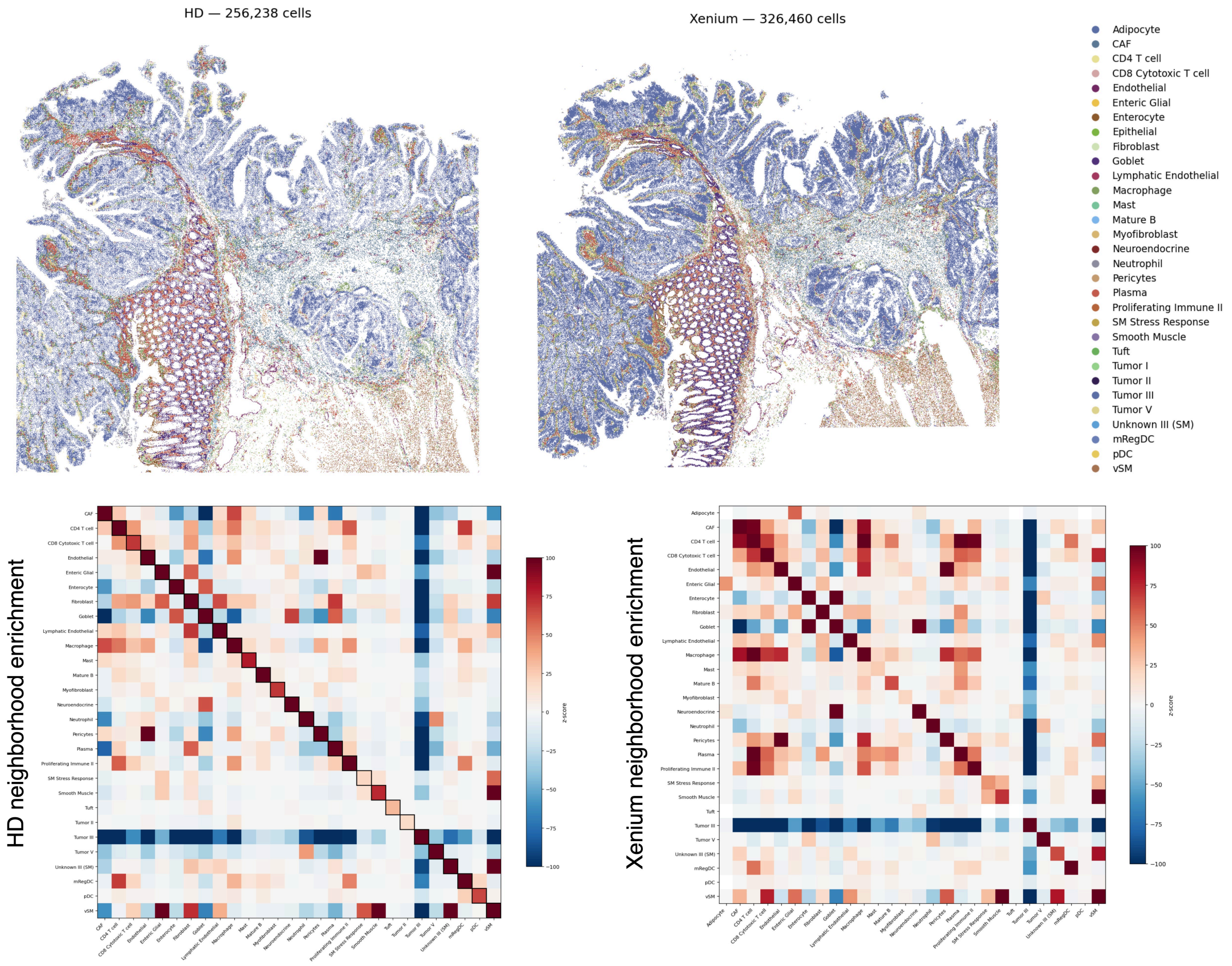


**Supplementary Figure 18. Cell-level neighbourhood enrichment on the CRC cohort across two spatial platforms. Top,** Cell-resolution maps of CRC tissue from Visium HD (256,238 CellART-segmented cells, left) and Xenium (326,460 cells, right) with the shared cell-type legend on the right. **Bottom,** Squidpy permutation-based neighbourhood-enrichment z-score matrices computed by STAT for each platform; the two matrices recover concordant spatial co-localisation patterns between cell-type pairs across the two technologies.

**Supplementary Tables**

#### Supplementary Table 1 — Methods comparison

Methods comparison: STAT and seven AI agents for biomedical / spatial transcriptomics analysis. Bolded cells in the STAT row mark properties not held by any other system in the table.

| System | Domain | Primary intent | Interaction | Spatial GUI | ROI on tissue | Session state | Method selection | Backend |
|---|---|---|---|---|---|---|---|---|
| **STAT** | Spatial transcriptomics | **Integrated, interactive analysis as a conversation** | **Multi-turn with shared GUI** | **Shared, programmatic agent access; interactive overlays + drawn ROI** | **Drawn-on-tissue ROI persisted as named reference** | **Persistent shared object** (platform, data level, annotations queryable across turns) | **Compatibility filter → semantic match → prereq verify** | **Custom multi-agent (no LangChain / LangGraph / MCP)** |
| Biomni | General biomedicine (25 subfields) | Autonomous execution across diverse biomedical tasks | Autonomous one-shot | Web chat UI; no tissue viewer | None | Per-query (data passed as files) | LLM dynamic retrieval over 150 tools / 105 software / 59 databases | Custom (no LangChain / LangGraph) |
| CellAgent | scRNA-seq | Zero-code automated single-cell analysis | Autonomous one-shot | Web chat UI; no tissue viewer | None | Per-query; code-step memory | Planner → Executor → Evaluator over tool list | Custom multi-agent |
| CellVoyager | scRNA-seq | Autonomously surface analyses the user overlooked | Multi-turn within Jupyter | Jupyter notebook; agent interprets figures via VLM (GPT-4o) | None | Jupyter kernel state | LLM blueprint + self-critique + VLM result interpretation | Jupyter + LLM (custom) |
| scAgent | scRNA-seq | Universal cell annotation across tissues | Multi-turn with planning | None (text chat); no tissue viewer | None | Memory module (datasets, embeddings, history; Milvus vector DB) | Planner generates action sequence over plugin space (scGPT + 30+ MoE-LoRA plugins) | Custom multi-agent (Plan + Action + Memory) |
| ChatSpatial | Spatial transcriptomics | Schema-enforced reproducible orchestration | Conversational multi-turn (CLI) | None (paper contrasts itself with “GUI-Based Platforms”) | None | Conversational stateful context | LLM selects from MCP-validated schemas (60+ methods, 15 categories) | MCP-based orchestration |
| SpatialAgent | Spatial transcriptomics | Autonomous spatial-biology pipeline (design → analysis → hypothesis) | Autonomous, optional co-pilot (text follow-up) | None stated in paper | None | Multimodal context memory (semantic + episodic) | Planner with templates, action module over 19 tools | Custom (no LangChain / LangGraph in paper) |
| STAgent | Spatial transcriptomics | Autonomous multimodal agent compressing weeks → minutes | Autonomous; Streamlit frontend allows multi-turn | Streamlit frontend; agent perceives tissue via VLM visual reasoning | None as named reference | Multimodal context memory | Planner + RAG-aided code generation over 19 tools | LangChain + LangGraph |

#### Supplementary Table 2 — Capability matrix

Capability matrix of STAT and seven AI agents ( ✓ supported / partial / ✕ not supported). STAT is the only system that holds every property in the matrix. STAT 16/16; ChatSpatial 6/16; STAgent 5/16; CellVoyager 5/16; SpatialAgent 4/16; scAgent 3/16; Biomni 2/16; CellAgent 1/16.

| Capability | STAT | Biomni | CellAgent | CellVoyager | scAgent | ChatSpatial | SpatialAgent | STAgent |
|---|---|---|---|---|---|---|---|---|
| **Spatial-data handling** | | | | | | | | |
| Spatial coordinates as first-class data | ✓ | ✕ | ✕ | ✕ | ✕ | ✓ | ✓ | ✓ |
| Cell-resolution vs spot-level distinction | ✓ | ✕ | ✕ | ✕ | ✕ | ✓ | ✓ | ✓ |
| Multi-slice / multi-sample in one session | ✓ | ✕ | ✕ | ✕ | ✕ | ✓ | partial | partial |
| **Shared state with agent** | | | | | | | | |
| Persistent session schema | ✓ | ✕ | ✕ | ✕ | ✕ | ✕ | ✕ | ✕ |
| Shared spatial GUI | ✓ | ✕ | ✕ | ✕ | ✕ | ✕ | ✕ | ✓ |
| Programmatic agent access to GUI state | ✓ | ✕ | ✕ | ✕ | ✕ | ✕ | ✕ | ✕ |
| Live overlay auto-update on data modification | ✓ | ✕ | ✕ | ✕ | ✕ | ✕ | ✕ | ✕ |
| Drawn ROI as named agent reference | ✓ | ✕ | ✕ | ✕ | ✕ | ✕ | ✕ | ✕ |
| **Interactivity** | | | | | | | | |
| Multi-turn analytical refinement | ✓ | ✕ | ✕ | ✓ | ✓ | ✓ | partial | partial |
| Clarification on ambiguity | ✓ | ✕ | ✕ | ✕ | ✕ | ✕ | ✕ | ✕ |
| **Architecture & reliability** | | | | | | | | |
| Skill compatibility pre-filter (data-type aware) | ✓ | ✕ | ✕ | ✕ | ✕ | partial | ✕ | ✕ |
| Staged selection pipeline (filter → match → verify) | ✓ | ✕ | ✕ | ✕ | ✕ | ✕ | ✕ | ✕ |
| Classified error reflection (fixable vs unsupportable) | ✓ | partial | partial | partial | partial | ✕ | partial | partial |
| Cross-LLM stability evaluated | ✓ | ✕ | ✕ | ✓ | ✕ | ✓ | ✕ | ✕ |
| Jupyter notebook export of full session | ✓ | ✕ | ✕ | ✓ | ✕ | ✕ | ✕ | partial |
| Independent of LangChain / LangGraph / MCP | ✓ | ✓ | ✓ | ✓ | ✓ | ✕ | ✓ | ✕ |

#### Supplementary Table 3 — Benchmark query set

Forty benchmark queries spanning 11 task categories and four difficulty regimes, distributed across three datasets (D1 = DLPFC Visium, 4 slices, 3.4–3.6K spots, 33,538 genes; D2 = MERFISH mouse brain, 2 slices, 32K and 82K cells, 1,122 genes; D3 = Xenium breast cancer, 1 slice, 160K cells, 306 genes).

| Code | Task category | Description | # Queries |
|---|---|---|---|
| T1 | Cell Type Annotation | Assign biologically meaningful cell-type labels. | 4 |
| T2 | Spatial Domain Detection | Identify spatially coherent tissue regions. | 3 |
| T3 | Spatially Variable Genes | Detect genes with significant spatial expression patterns. | 3 |
| T4 | Cell-Cell Communication | Analyse ligand-receptor interactions between cell types. | 3 |
| T5 | Neighborhood Enrichment | Test spatial co-localisation / avoidance between cell types. | 3 |
| T6 | Deconvolution | Estimate cell-type proportions per spot (Visium). | 2 |
| T7 | Niche Detection | Identify spatial microenvironments from cell composition. | 2 |
| T8 | Differential Expression | Find DE genes between specified groups. | 2 |
| T9 | GO / Pathway Enrichment | Statistical gene-set enrichment analysis. | 2 |
| T10 | Batch Integration | Correct batch effects across multiple tissue slices. | 2 |
| T11 | General / Exploratory | Dataset summary, exploratory analysis. | 4 |
| — | Multi-step | Sequential multi-task pipelines. | 4 |
| — | Infeasible | Task with missing required data. | 1 |
| — | Out-of-scope | Task beyond spatial transcriptomics scope. | 3 |
| — | Missing prerequisite | Task requiring a prior step not yet performed. | 2 |

**Complete query list (Q01–Q40).** Where applicable, scripted responses were provided to handle multi-turn clarification.

| ID | Task | Dataset | Difficulty | Query |
|---|---|---|---|---|
| Q01 | T1 | D2 | clear | "Annotate cell types on this tissue. It is a mouse brain cortex sample." |
| Q02 | T1 | D3 | clear | "Identify cell types in this breast cancer tissue sample. I have a single-cell reference dataset at `./breast_cancer_reference.h5ad` with cell-type annotations that you can optionally use." |
| Q03 | T1 | D2 | vague | "What cell types are present in this tissue?" |
| Q04 | T1 | D3 | vague | "Can you identify the different cell populations here? I have a reference scRNA-seq dataset at `./breast_cancer_reference.h5ad` if that helps." |
| Q05 | T2 | D1 | clear | "Identify spatial domains in this brain tissue. There should be about 7 distinct regions including cortical layers and white matter." |
| Q06 | T2 | D1 | vague | "Can you find the spatial structure in this tissue?" |

| ID | Task | Dataset | Difficulty | Query |
|---|---|---|---|---|
| Q07 | T2 | D2 | vague | "Identify distinct tissue regions based on spatial gene expression." |
| Q08 | T3 | D1 | clear | "Find genes that show significant spatial expression patterns across this tissue." |
| Q09 | T3 | D1 | vague | "Which genes have interesting spatial patterns?" |
| Q10 | T3 | D2 | clear | "Identify spatially variable genes in this mouse brain section." |
| Q11 | T4 | D2 | clear | "Analyze ligand-receptor interactions between cell types in this mouse brain tissue." |
| Q12 | T4 | D3 | clear | "Find cell-cell communication patterns between tumor and immune cells in this human breast cancer sample." |
| Q13 | T4 | D3 | vague | "How are the cells communicating with each other?" |
| Q14 | T5 | D2 | clear | "Test whether certain cell types tend to co-localize or avoid each other spatially in this mouse brain tissue." |
| Q15 | T5 | D3 | clear | "Analyze spatial co-localization patterns between cell types in this breast cancer sample." |
| Q16 | T5 | D2 | vague | "Are certain cell types found near each other in this tissue?" |
| Q17 | T6 | D1 | clear | "Estimate cell-type proportions for each spot in this Visium tissue. I have a single-cell reference at `./dlpfc_reference.h5ad` with cell-type annotations." |
| Q18 | T6 | D1 | vague | "What cell types make up each spot in this tissue?" |
| Q19 | T7 | D2 | clear | "Identify spatial microenvironments (niches) based on the local cell-type composition in this mouse brain tissue." |
| Q20 | T7 | D2 | vague | "Find the spatial neighborhoods in this tissue." |
| Q21 | T8 | D1 | clear | "Find differentially expressed genes between the different spatial domains in this tissue." |
| Q22 | T8 | D3 | vague | "What genes distinguish the tumor regions from the surrounding stroma?" |
| Q23 | T9 | D1 | clear | "Run gene-ontology enrichment on these marker genes: RELN, CUX2, PCP4, MBP, MOBP, NTNG2, TLE4. Species: human." |
| Q24 | T9 | D3 | vague | "What biological pathways are active in the tumor microenvironment?" |
| Q25 | T10 | D1×4 | clear | "Integrate these four brain tissue slices, correcting for batch effects while preserving biological variation." |
| Q26 | T10 | D2×2 | vague | "These two tissue sections are from the same brain region but look quite different. Can you make them comparable?" |
| Q27 | T11 | D1 | clear | "Give me a summary of this dataset — how many spots, how many genes, what spatial extent, and what annotations are already present." |
| Q28 | T11 | D2 | clear | "Show the top 10 most highly expressed genes across all cells in this tissue." |
| Q29 | T11 | D2 | vague | "Give me a complete analysis of this spatial transcriptomics data." |
| Q30 | T11 | D1 | vague | "What can you tell me about this tissue?" |
| Q31 | Multi | D2 | multi-step-task | "First annotate cell types, then analyze which cell types tend to co-localize in this mouse brain tissue." |
| Q32 | Multi | D1 | multi-step-task | "Identify spatial domains in this brain tissue (about 7 regions), then find genes that are differentially expressed between them." |
| Q33 | Multi | D3 | multi-step-task | "Annotate cell types in this breast cancer sample using the reference at `./breast_cancer_reference.h5ad`, then analyze cell-cell communication between tumor and immune cells." |
| Q34 | Multi | D3 | multi-step-task | "Identify cell types, find marker genes for each type, then run pathway enrichment on the tumor-cell markers." |

| ID | Task | Dataset | Difficulty | Query |
|---|---|---|---|---|
| Q35 | Spec. | D1 | infeasible | "Estimate cell-type proportions for each spot without any reference dataset." |
| Q36 | Spec. | D3 | out-of-scope | "Segment individual cells from the H&E image." |
| Q37 | Spec. | D2 | out-of-scope | "Perform RNA velocity analysis on this spatial data." |
| Q38 | Spec. | D3 | out-of-scope | "Train a deep-learning classifier to predict patient survival from this tissue." |
| Q39 | Spec. | D2 | missing-prereq | "Identify spatial niches in this tissue based on cell-type composition." |
| Q40 | Spec. | D1 | missing-prereq | "Test whether cell types co-localize spatially in this tissue." |

**Notes.** For Q39 the input dataset (`mouse_brain_no_celltype.h5ad`) lacks cell-type annotations, which are required for niche detection. For Q40 the DLPFC dataset has spatial-layer annotations but no cell-type column, requiring deconvolution or annotation as a prerequisite.

#### Supplementary Table 4 — LLM-judge evaluation protocol

Claude Opus 4.6 was used as an expert judge to evaluate each system's output across all 40 benchmark queries. For each query the judge received the complete output of all four systems in a single prompt, enabling direct comparison; system identities were anonymised (labelled A–D) to prevent bias.

##### Judge prompt

**System prompt.** You are an expert evaluator for spatial transcriptomics AI agents.

**User prompt.** A user asked the following question about a spatial transcriptomics dataset:

User Query: `{query}`
Dataset: `{dataset_description}`
Expected behavior: `{expected_behavior}`

Below are the responses from 4 different AI systems (A–D). For each system you see its text response, code (if any), execution output, and whether it produced a modified dataset.

`{all_responses_block}`

For EACH system (A–D), judge:

1. **Success** — did the system accomplish the task? Answer YES or NO. YES = the analysis was performed correctly and produced usable results. NO = error, wrong approach, no output, or the system could not handle the task. For out-of-scope queries: YES if the system correctly explained why the task cannot be done. For prerequisite-missing queries: YES if the system correctly identified what is missing.
2. **Quality** (only if Success = YES) — rate 1–5: 5 excellent (correct method, good parameters, biologically sound); 4 good (minor issues); 3 acceptable (notable issues); 2 marginal (questionable); 1 poor (clearly wrong or misleading).
3. **Rank** — rank all SUCCESSFUL systems from best (1) to worst. Failed systems get no rank.

Respond in the following JSON format (no surrounding prose):

```
{
  "A": {"success": true/false, "quality": null|1-5, "rank": null|int, "reason": "..."},
  "B": {"success": true/false, "quality": null|1-5, "rank": null|int, "reason": "..."},
  "C": {"success": true/false, "quality": null|1-5, "rank": null|int, "reason": "..."},
  "D": {"success": true/false, "quality": null|1-5, "rank": null|int, "reason": "..."}
}
```

##### Per-system response block format

For each system, the judge received: the agent text response (truncated to 2000 characters), the generated Python code (truncated to 1500 characters), execution stdout (truncated to 500 characters), execution stderr (truncated to 500 characters), a flag indicating whether an output dataset was saved, and (if saved) a summary of the output dataset including the number of cells/spots, new columns added to `obs`, new keys added to `obsm`/`uns`, and sample values from key result columns.

##### Task-specific success criteria

| Task type | Success criterion |
|---|---|
| Cell-type annotation (T1) | Produced a `celltype` column with biologically meaningful labels (not numeric cluster IDs). |
| Spatial-domain detection (T2) | Assigned spatial-domain labels to cells / spots. |
| Spatially variable genes (T3) | Identified genes with spatial statistics (Moran's I, SpatialDE q-values), not just expression variance. |
| Cell-cell communication (T4) | Identified ligand-receptor interactions using a proper database (CellPhoneDB, CellChat, LIANA). |
| Neighborhood enrichment (T5) | Performed spatial co-localisation or enrichment analysis with statistical testing. |
| Deconvolution (T6) | Estimated cell-type proportions per spot using the provided reference. |
| Niche detection (T7) | Identified spatial niches from local cell-type composition. |
| Differential expression (T8) | Identified DE genes between specified groups with statistical testing. |
| GO / pathway enrichment (T9) | Performed formal statistical enrichment (not manual gene categorisation). |
| Batch integration (T10) | Loaded ALL input files and applied batch correction (Harmony, BBKNN, scVI). |
| General / Exploratory (T11) | Provided a meaningful summary or analysis of the dataset. |
| Multi-step (Q31–Q34) | ALL steps completed — partial completion is judged as failure. |
| Infeasible (Q35) | Attempted a reasonable alternative OR correctly explained the limitation. |

| Task type | Success criterion |
|---|---|
| Out-of-scope (Q36–Q38) | Correctly explained why the task cannot be done with the available data / tools. |
| Missing prerequisite (Q39–Q40) | Identified the missing prerequisite OR performed it as a preliminary step. |

#### Aggregated metrics

From the 40 per-query judgments we computed: **Success rate** (fraction of queries where the system was judged successful); **Mean quality** (average quality score across successful queries, 1–5); **Mean rank** (average rank across successful queries, lower is better); **Win rate** (fraction of queries where the system achieved rank 1).

##### Supplementary Table 5 — Failure and low-quality response analysis

Aggregate counts of failures (judge-assigned success = no) and low-quality completions (success = yes but quality ≤ 3) across the three execution-capable systems on the 40-query benchmark. The vanilla LLM is excluded from the per-query analysis (it failed 31/40 queries due to fundamental architectural limitations and is summarised separately in Supplementary Fig. 1).

| System | Failures (N) | Low quality (Q ≤ 3) | Total issues |
|---|---|---|---|
| Biomni | 5 / 40 | 8 / 35 | 13 |
| SpatialAgent | 3 / 40 | 4 / 37 | 7 |
| STAT | 1 / 40 | 4 / 39 | 5 |

**Biomni — failures (5 queries).**

> **Q23 — GO Enrichment (T9, clear)**
> **Query**: "Run gene-ontology enrichment on these marker genes: RELN, CUX2, PCP4, MBP, MOBP, NTNG2, TLE4. Species: human."
> **What happened**: Performed manual GO-style annotation due to dependency issues — gene-function descriptions and spatial expression patterns, but no statistical enrichment test.
> **Root cause**: No actual GO enrichment was run; only manual literature-based gene annotation, which does not constitute a statistical enrichment test.

> **Q36 — H&E cell segmentation (out-of-scope)**
> **Query**: "Segment individual cells from the H&E image."
> **What happened**: Attempted segmentation despite the dataset already being at single-cell resolution; added morphological-estimate columns. Did not flag the task as out-of-scope.
> **Root cause**: Fabricated morphological features (area, eccentricity, solidity) without actual image data and did not acknowledge the out-of-scope nature.

> **Q37 — RNA velocity (out-of-scope)**
> **Query**: "Perform RNA velocity analysis on this spatial data."
> **What happened**: Attempted scVelo with a "spatial-gradient" alternative; acknowledged missing spliced / unspliced layers but produced velocity-named columns anyway.
> **Root cause**: RNA velocity requires spliced / unspliced count matrices; the substituted spatial-gradient approach is a fabricated alternative branded as the original task.

> **Q38 — Survival prediction (out-of-scope)**
> **Query**: "Train a deep-learning classifier to predict patient survival from this tissue."
> **What happened**: Trained a survival classifier after creating synthetic survival labels; added `patient_id`, `predicted_survival`, `risk_score` columns.
> **Root cause**: Survival prediction requires real clinical outcome data; training on synthetic labels produces meaningless results.

> **Q40 — Co-localisation without cell types (missing-prereq)**
> **Query**: "Test whether cell types co-localize spatially in this tissue."
> **What happened**: Attempted Leiden clustering as a substitute and then ran co-localisation; the output `h5ad` only contains a 3-cluster `leiden` column with no actual cell-type labels.
> **Root cause**: The pipeline used Leiden clusters or layer labels as cell-type substitutes rather than identifying or addressing the missing prerequisite.

**Biomni — low-quality results (Q ≤ 3, 8 queries).**

> **Q05 — Q2 — Spatial domains (T2, clear)**
> **Root cause**: Identified only 4 spatial domains (L3, L5, WM, L6) instead of the expected 7 cortical layers + WM; significantly under-segmented despite multi-resolution clustering.

> **Q09 — Q3 — SVG detection (T3, vague)**
> **Root cause**: Used a simplistic "spatial CV" metric rather than proper spatial statistics (Moran's I, SpatialDE); no formal statistical testing stored in the `h5ad`.

> **Q10 — Q3 — SVG detection (T3, clear)**
> **Root cause**: Used simplistic correlation with spatial coordinates rather than spatial autocorrelation; flagged 65 genes with score > 0.2.

> **Q11 — Q3 — Cell-cell communication (T4, clear)**
> **Root cause**: Used only 3 manually curated ligand-receptor pairs rather than a comprehensive database; results biologically reasonable but limited in scope.

**Q15 — Q3 — Neighborhood enrichment (T5, clear)**
**Root cause**: Distance-based methodology rather than permutation-based; claimed 8–10× enrichment but storage suggests limited statistical rigour.

**Q17 — Q2 — Deconvolution (T6, clear)**
**Root cause**: NNLS deconvolution produced extremely poor diversity (Mix_1 in 92.5% of spots) — failed to resolve cell-type mixtures within spots.

**Q18 — Q2 — Deconvolution (T6, vague)**
**Root cause**: 75.5% of spots assigned to a single type (Ex_5_L5) — closer to label transfer than deconvolution; misses the spot-as-mixture premise.

**Q26 — Q3 — Batch integration (T10, vague)**
**Root cause**: Batch-mixing score of 0.294 indicates incomplete ComBat correction; 82 clusters is heavily over-segmented.

**SpatialAgent — failures (3 queries).**

**Q23 — GO Enrichment (T9, clear)**
**Query**: "Run gene-ontology enrichment on these marker genes: RELN, CUX2, PCP4, MBP, MOBP, NTNG2, TLE4. Species: human."
**What happened**: Categorised genes into myelination / neural-development / synaptic-function / transcriptional-regulation classes with spatial expression analysis.
**Root cause**: Despite claiming GO enrichment, the analysis was manual functional categorisation, not a proper statistical enrichment test against a database (Enrichr, DAVID, gseapy).

**Q36 — H&E cell segmentation (out-of-scope)**
**Query**: "Segment individual cells from the H&E image."
**What happened**: Recognised data is already single-cell and substituted spatial clustering; did not clearly flag the task as out-of-scope.
**Root cause**: Substituted clustering for actual H&E segmentation (which would require Cellpose or StarDist), without explaining the out-of-scope nature.

**Q38 — Survival prediction (out-of-scope)**
**Query**: "Train a deep-learning classifier to predict patient survival from this tissue."
**What happened**: Created synthetic survival outcomes from tumour features and trained a classifier; added survival columns to the output.
**Root cause**: Trained a model on fabricated outcomes; results are meaningless without actual clinical outcome data.

**SpatialAgent — low-quality results (Q ≤ 3, 4 queries).**

**Q06 — Q3 — Spatial domains (T2, vague)**
**Root cause**: Found 10 GraphST domains (over-segmented vs  7 expected); numeric domain labels without biological annotation.

**Q35 — Q3 — Infeasible deconvolution**
**Root cause**: Marker-based 8-celltype proportion estimate is reasonable but limited; acknowledged the lack of reference as a constraint.

**Q37 — Q3 — RNA velocity (out-of-scope)**
**Root cause**: Correctly recognised the lack of spliced/unspliced layers and pivoted to PAGA trajectory; pivot could be more clearly communicated.

**Q40 — Q3 — Co-localisation (missing-prereq)**
**Root cause**: Used brain-layer annotations as a meaningful proxy for cell types and ran proper co-localisation; could have noted more explicitly that layers are not cell types.

**STAT — failure (1 query).**

**Q37 — RNA velocity (out-of-scope)**
**Query**: "Perform RNA velocity analysis on this spatial data."
**What happened**: Attempted scVelo without checking for spliced / unspliced layers; the output `h5ad` retains the original `celltype` but no velocity results are evident.
**Root cause**: RNA velocity is not in STAT's skill registry, so the matcher correctly returned no match; however, the current pipeline still invoked the code generator on the raw query rather than declining, and the generator called `scVelo` without checking for spliced / unspliced layers. An explicit no-skill refusal route would close this gap.

**STAT — low-quality results (Q ≤ 3, 4 queries).**

**Q07 — Q3 — Spatial domains (T2, vague)**
**Root cause**: Used basic K-means rather than spatial-aware methods on MERFISH; no biological annotation of regions in the output.

**Q35 — Q3 — Infeasible deconvolution**
**Root cause**: NMF-based unsupervised deconvolution is the principled reference-free approach; explicitly acknowledged the constraint and 7 components aligned with the DLPFC layer structure. Quality is bounded by inherent task difficulty.

**Q36 — Q3 — H&E segmentation (out-of-scope)**
**Root cause**: Correctly attempted to access H&E images and discovered they were unavailable; the response could have communicated the out-of-scope nature more explicitly.

**Q39 — Q3 — Niche detection (missing-prereq)**
**Root cause**: Correctly identified the missing celltype prerequisite and asked the user for a reference; could have proposed an unsupervised annotation as a preliminary step instead of stopping.

#### Common failure patterns

**Pattern 1 — Fabrication instead of refusal** Biomni Q38 and SpatialAgent Q38 both created synthetic survival data and trained classifiers on fabricated labels rather than declaring the task infeasible. Generating plausible but meaningless results is worse than refusing the task.

**Pattern 2 — Superficial alternatives for missing data** When required data is absent (e.g., spliced/unspliced for RNA velocity), some systems fabricated alternative analyses branded under the original task name (Biomni Q37).

**Pattern 3 — GO enrichment without statistical testing** Both Biomni and SpatialAgent performed manual gene categorisation or literature-based annotation on Q23 instead of formal statistical enrichment; only STAT used `gseapy` with Fisher's exact test.

**Pattern 4 — Deconvolution quality** Biomni's deconvolution consistently produced low-diversity results (>75% of spots assigned to a single dominant type) on Q17/Q18, suggesting the method failed to resolve mixtures.

**Pattern 5 — Prerequisite identification** STAT correctly identified missing prerequisites (Q39) but chose to ask the user rather than proceed with an unsupervised alternative — a more conservative but potentially less helpful default.

##### Supplementary Table 6 — LLM-backbone ablation, failures on the 40-query benchmark

Per-LLM success counts when the same 40-query benchmark of Fig. 2 was re-run with seven different frontier backbones, holding the STAT pipeline architecture, skill registry, and LLM-judge protocol identical to the main benchmark.

| Backbone model | Success | Fail | Success rate |
|---|---|---|---|
| Claude Sonnet 4 | 39 / 40 | 1 | 97.5% |
| Grok-3 | 38 / 40 | 2 | 95.0% |
| Grok-4 | 38 / 40 | 2 | 95.0% |
| GPT-5.4 | 37 / 40 | 3 | 92.5% |
| Gemini 3.1 Pro | 37 / 40 | 3 | 92.5% |
| DeepSeek V3.2 | 37 / 40 | 3 | 92.5% |
| GPT-4o | 36 / 40 | 4 | 90.0% |

**Claude Sonnet 4 — 1 failure.**

> **Q37 — RNA velocity (out-of-scope)**
> **What happened**: Attempted scVelo, hit `KeyError: 'spliced'`, and then proceeded with a fake velocity analysis using `np.random.normal` vectors as a demonstration.
> **Root cause**: Fabricated velocity vectors after the initial failure rather than declaring the analysis infeasible without spliced / unspliced counts.

**DeepSeek V3.2 — 3 failures.**

> **Q07 — Spatial domains (T2, vague)**
> **What happened**: No spatial-domain assignments in the output.
> **Root cause**: Analysis code did not produce or persist spatial-domain labels.

> **Q17 — Deconvolution (T6, clear)**
> **What happened**: RCTD code present in the response but results not persisted to the output `h5ad`.
> **Root cause**: Code execution completed but deconvolution weights were not saved to the output file.

> **Q24 — Pathway enrichment (T9, vague)**
> **What happened**: No pathway / GO enrichment performed.
> **Root cause**: Model did not follow through to the enrichment step after identifying relevant cell populations.

> **Q32 — Multi-step: domains + DE**
> **What happened**: Spatial domains identified but no DE analysis performed.
> **Root cause**: Only the first step (domain detection) completed; the second step was not executed.

**Grok-4 — 2 failures.**

> **Q07 — Spatial domains (T2, vague)**
> **What happened**: Attempted Leiden clustering for spatial regions, encountered errors, did not save domain assignments.
> **Root cause**: Clustering errored during execution; no fallback or retry was attempted.

> **Q34 — Multi-step: annotate + markers + GO**
> **What happened**: No celltype, marker genes, or GO results in output despite code attempting all three steps.
> **Root cause**: Code execution failed silently; none of the three pipeline steps produced persisted results.

**GPT-5.4 — 3 failures.**

> **Q03 — Cell-type annotation (T1, vague)**
> **What happened**: Did not annotate cell types; only checked for an existing `celltype` column and reported none available.
> **Root cause**: Stopped at the existence check rather than running annotation on unannotated data.

> **Q21 — Differential expression (T8, clear)**
> **What happened**: Refused to run DE, claiming spatial domains were not yet defined; failed to recognise the existing `layer` column.
> **Root cause**: Did not inspect available `obs` columns; the `layer` column (L1–L6 + WM) was the intended grouping.

> **Q26 — Batch integration (T10, vague)**
> **What happened**: Loaded both slices and aligned to shared gene space, compared composition and expression, but applied no batch-correction method (no BBKNN / ComBat / Harmony).
> **Root cause**: Treated feature alignment as sufficient; no batch-correction algorithm was invoked.

**Gemini 3.1 Pro — 3 failures.**

**Q03 — Cell-type annotation (T1, vague)**
**What happened**: Same pattern as GPT-5.4 — checked for an existing `celltype` column and reported none available.
**Root cause**: Stopped at the existence check rather than performing annotation on unannotated data.

**Q07 — Spatial domains (T2, vague)**
**What happened**: Attempted Leiden clustering but did not save domain assignments to the output.
**Root cause**: Clustering performed but results were not persisted to the output file.

**Q24 — Pathway enrichment (T9, vague)**
**What happened**: Performed Wilcoxon DE between tumour and immune cells and identified upregulated genes, but did not run formal GO / pathway enrichment.
**Root cause**: Only the first step (DE) was completed; the enrichment step was omitted.

**Q32 — Multi-step: domains + DE**
**What happened**: Identified 7 SpaGCN domains but no DE results stored; `spatial_domain` column present but no `rank_genes_groups`.
**Root cause**: Domain detection completed successfully but downstream DE was not persisted.

**Grok-3 — 2 failures.**

**Q04 — Cell-type annotation (T1, vague, with reference)**
**What happened**: scANVI ran and produced latent representations in `obsm`, but celltype labels were not transferred to `obs`.
**Root cause**: Model trained and stored embeddings; the final label-transfer / prediction step was not completed or not persisted.

**Q17 — Deconvolution (T6, clear)**
**What happened**: RCTD code attempted but failed with an error; no deconvolution weights produced.
**Root cause**: Code execution error during the RCTD run; the pipeline did not recover or produce partial results.

**Q38 — Survival prediction (out-of-scope)**
**What happened**: Did not recognise the task as out-of-scope; simulated survival outcomes and trained a model on fabricated data.
**Root cause**: Created synthetic `survival_time`/`survival_status` columns and trained a classifier — meaningless without real clinical outcome data.

**GPT-4o — 4 failures.**

**Q01 — Cell-type annotation (T1, clear)**
**What happened**: Got stuck requesting a reference dataset and did not proceed with marker-based annotation despite being told no reference was available.
**Root cause**: Entered a clarification loop asking for a reference even after the scripted response said none was available; did not fall back to unsupervised marker-based annotation.

**Q03 — Cell-type annotation (T1, vague)**
**What happened**: Same pattern as GPT-5.4 / Gemini — checked for the `celltype` column and reported none available.
**Root cause**: Existence check rather than annotation.

**Q25 — Batch integration (T10, clear)**
**What happened**: Refused to integrate, incorrectly claiming BBKNN requires cell-type annotations. All 4 slices loaded (14,243 spots) but no batch correction applied.
**Root cause**: Incorrect understanding of BBKNN prerequisites — BBKNN needs only a batch key and PCA embeddings, not cell types.

**Q24 — Pathway enrichment (T9, vague)**
**What happened**: Described a pathway-analysis plan but did not execute enrichment; no GO results in output.
**Root cause**: Planning was done but code execution did not follow through to produce enrichment results.

**Q26 — Batch integration (T10, vague)**
**What happened**: Loaded both slices, normalised, and aligned features, but applied no batch correction.
**Root cause**: Feature alignment treated as integration; no batch-correction algorithm was applied.

#### Common failure patterns

| Pattern | Affected models | Queries |
|---|---|---|
| **Check-but-don't-act** — checked for existing annotations but did not run annotation on unannotated data | GPT-5.4, Gemini, GPT-4o | Q03 |
| **Incomplete multi-step** — first step completed, subsequent steps not executed or not persisted | DeepSeek, Gemini, Grok-4 | Q32, Q34 |

| Pattern | Affected models | Queries |
|---|---|---|
| **Results not persisted** — analysis ran but output not saved to `h5ad` | DeepSeek, Grok-3, Gemini | Q04, Q07, Q17 |
| **Incorrect prerequisite assumptions** — refused valid analysis citing wrong prerequisites | GPT-4o, GPT-5.4 | Q21, Q25 |
| **Fabrication on infeasible tasks** — created synthetic data rather than declaring infeasibility | Grok-3, Claude | Q37, Q38 |
| **Incomplete enrichment** — DE completed but GO / pathway step omitted | DeepSeek, Gemini, GPT-4o | Q24 |

#### Supplementary Table 7 — Pipeline test queries (Set A, planner)

Set A tests the `QueryPlanner` stage in isolation: it determines target slices, decomposes multi-step queries, and identifies ambiguous queries requiring clarification. 30 queries across 4 categories were tested on six mock sessions. **Binary success criterion**: all three fields (`target_slice_ids`, `num_steps`, `needs_clarification`) must match the expected ground truth. For multi-step queries with `alt_expected` listed, any valid decomposition is accepted.

**Mock sessions used in Sets A, B, and C.**

| ID | Description | Slices |
|---|---|---|
| `1-spot` | 1 DLPFC Visium slice | S0: spot-level, 4,226 spots, 33,538 genes |
| `1-cell` | 1 MERFISH mouse brain | S0: cell-level, 80,143 cells, 550 genes, 12 cell types |
| `1-cell-noCT` | 1 MERFISH, no celltype | S0: cell-level, 80,143 cells, 550 genes, no celltype column |
| `2-cell` | 2 MERFISH slices | S0+S1: cell-level, each 80,143 cells, 12 cell types |
| `4-spot` | 4 DLPFC Visium slices | S0–S3: spot-level, 4,200 spots each |
| `mixed` | 1 Visium + 1 MERFISH | S0: DLPFC spot; S1: breast cancer cell, 15 cell types |

**Set A queries (PA01–PA30).**

| ID | Session | Query | Expected slices | Steps | Clarify | Edge case |
|---|---|---|---|---|---|---|
| PA01 | `1-spot` | "Find spatially variable genes." | [[0]] | 1 | No | Single implicit target |
| PA02 | `1-cell` | "Run neighborhood enrichment analysis." | [[0]] | 1 | No | Single implicit target |
| PA03 | `2-cell` | "Annotate cell types on slice 1." | [[1]] | 1 | No | Explicit slice ID |
| PA04 | `2-cell` | "Detect spatial niches on the first section." | [[0]] | 1 | No | Ordinal "first" |
| PA05 | `mixed` | "Run deconvolution on slice 0." | [[0]] | 1 | No | Explicit ID, mixed session |
| PA06 | `mixed` | "Analyze communication in the breast cancer tissue." | [[1]] | 1 | No | Tissue-name reference |
| PA07 | `4-spot` | "Run SpatialDE on the third slice." | [[2]] | 1 | No | Ordinal "third" |
| PA08 | `mixed` | "Detect niches on slice 0." | [[0]] | 1 | No | Explicit ID, mixed session |
| PA09 | `1-spot` | "Analyze the cortical layers in this DLPFC tissue." | [[0]] | 1 | No | Indirect phrasing |
| PA10 | `4-spot` | "Run domain detection on slices 1 and 3." | [[1],[3]] | 2 | No | Non-adjacent IDs |
| PA11 | `2-cell` | "Find SVGs on both slices." | [[0],[1]] | 2 | No | "both" keyword |
| PA12 | `4-spot` | "Detect spatial domains on each slice with 7 clusters." | [[0],[1],[2],[3]] | 4 | No | "each" = N steps |
| PA13 | `4-spot` | "Integrate all four slices." | [[0,1,2,3]] | 1 | No | "all" = 1 cross-slice |
| PA14 | `2-cell` | "Compare gene expression between the two sections." | [[0,1]] | 1 | No | "compare" = cross-slice |
| PA15 | `2-cell` | "Which slice has more oligodendrocytes?" | [[0,1]] | 1 | No | Comparative question |

| ID | Session | Query | Expected slices | Steps | Clarify | Edge case |
|---|---|---|---|---|---|---|
| PA16 | `4-spot` | "Annotate cell types on every section except slice 2." | [[0],[1],[3]] | 3 | No | Exclusion "except" |
| PA17 | `2-cell` | "Annotate cell types." | — | — | Yes | Ambiguous target |
| PA18 | `mixed` | "Run spatial statistics." | — | — | Yes | Ambiguous modality |
| PA19 | `4-spot` | "Detect domains on one of the slices." | — | — | Yes | "one of" ambiguous |
| PA20 | `2-cell` | "I want to analyze this data." | — | — | Yes | Completely vague |
| PA21 | `1-cell-noCT` | "First annotate cell types, then analyze niches." | [[0],[0]] | 2 | No | Sequential same slice |
| PA22 | `2-cell` | "Annotate slice 0 then run niche detection on slice 1." | [[0],[1]] | 2 | No | Sequential, different |
| PA23 | `1-cell-noCT` | "Annotate cells, find DE genes between cell types, and run GO enrichment on the DE results." | [[0],[0],[0]] or [[0],[0]] | 2–3 | No | 3-step chain (flexible) |
| PA24 | `1-spot` | "Detect spatial domains, then find marker genes for each domain." | [[0],[0]] | 2 | No | Domain → DE dependency |
| PA25 | `1-cell-noCT` | "Identify cell types, analyze cell-cell communication, and detect spatial niches." | [[0],[0],[0]] or [[0],[0]] | 2–3 | No | 3-step from no-CT |
| PA26 | `mixed` | "Annotate the breast cancer tissue, then compare its cell types with the DLPFC tissue." | [[1],[0,1]] | 2 | No | Annotate one, compare both |
| PA27 | `2-cell` | "Integrate slices 0 and 1, then detect spatial domains on the merged result." | [[0,1],[0,1]] or [[0,1]] | 1–2 | No | Integration + downstream |
| PA28 | `1-cell-noCT` | "Run the full analysis pipeline: annotate cell types, detect niches, compute neighborhood enrichment, and find spatially variable genes." | Multiple valid | 1–4 | No | 4-step comprehensive |
| PA29 | `1-spot` | "Find spatially variable genes and then run pathway enrichment on the top SVGs." | [[0],[0]] | 2 | No | SVG → pathway chain |
| PA30 | `mixed` | "For each tissue, annotate cell types and then analyze cell communication." | [[0],[0],[1],[1]] or alt. | 2–4 | No | Per-tissue multi-step |

##### Supplementary Table 8 — Pipeline test queries (Set B, skill matching)

Set B tests the `SkillFilter` (programmatic, deterministic) and `SemanticMatcher` (LLM-based) stages together. Target slices are pre-set so the planner is bypassed. 30 queries across 3 categories. **Binary success criterion**: for in-scope queries (PB01–PB23) the top-1 matched skill must appear in the expected skill set; for out-of-scope queries (PB24–PB30) the matcher must return an empty match. The deterministic filter is identical across all backbones; only the LLM-based matcher varies. Skills with `default_skill = false` are excluded from the candidate pool, matching the production pipeline.

| ID | Session | Target | Query | Expected skill | Task |
|---|---|---|---|---|---|
| PB01 | `1-spot` | [0] | "Identify 7 spatial domains using graph-based clustering." | `spatial-domain-detection` | T2 |
| PB02 | `1-cell` | [0] | "Find spatially variable genes using Gaussian-process regression." | `svg-spatialde` | T3 |
| PB03 | `1-cell` | [0] | "Detect spatial microenvironments based on cell-type neighborhoods." | `niche-detection` or `spatial-stats-neighborhood-enrichment` | T7 |
| PB04 | `1-cell` | [0] | "Analyze ligand-receptor interactions in this mouse tissue. Species: mouse." | `cell-communication-liana` | T4 |
| PB05 | `1-cell` | [0] | "Identify cell types using clustering and marker genes. Tissue: mouse brain cortex." | `celltype-annotation-fast` | T1 |
| PB06 | `1-spot` | [0] | "Deconvolve cell-type proportions in each spot using my reference at `./ref.h5ad`." | `celltype-deconvolution` or `deconvolution-flashdeconv` | T6 |
| PB07 | `1-cell` | [0] | "Compute neighborhood enrichment z-scores between cell types." | `spatial-stats-neighborhood-enrichment` | T5 |
| PB08 | `1-cell` | [0] | "Calculate Moran's I spatial autocorrelation for genes PLP1, MBP, GFAP." | `spatial-statistics` | T5 |
| PB09 | `1-spot` | [0] | "Run gene-ontology enrichment on genes MOBP, PLP1, MBP. Species: human." | `pathway-go-enrichment` | T9 |
| PB10 | `2-cell` | [0,1] | "Correct batch effects between these two slices." | `integration-bbknn` | T10 |
| PB11 | `1-cell` | [0] | "Find differentially expressed genes between excitatory and inhibitory neurons." | `differential-expression` | T8 |
| PB12 | `1-cell` | [0] | "Transfer cell-type labels from single-cell reference via scANVI variational inference." | `celltype-annotation-scanvi` | T1 |
| PB13 | `1-cell` | [0] | "Compute GO enrichment for marker genes of each cell type. Species: mouse." | `pathway-go-enrichment` or `differential-expression` | T9 |
| PB14 | `1-spot` | [0] | "Perform cell-cell communication analysis on this human tissue. Species: human." | `cell-communication-liana` | T4 |
| PB15 | `1-spot` | [0] | "Detect spatial domains in this tissue using unsupervised clustering." | `spatial-domain-detection` | T2 |
| PB16 | `1-cell` | [0] | "What genes show interesting spatial patterns?" | `svg-spatialde` | T3 |
| PB17 | `1-cell` | [0] | "Are certain cell types found near each other more often than expected?" | `spatial-stats-neighborhood-enrichment` | T5 |
| PB18 | `1-spot` | [0] | "Can you find the spatial structure in this tissue?" | `spatial-domain-detection` | T2 |
| PB19 | `1-cell` | [0] | "How are cells communicating with each other?" | `cell-communication-liana` | T4 |
| PB20 | `1-cell` | [0] | "What makes tumor cells different from stromal cells at the gene level?" | `differential-expression` | T8 |
| PB21 | `1-spot` | [0] | "What is the cellular composition of each spot?" | `celltype-deconvolution` or `deconvolution-flashdeconv` | T6 |
| PB22 | `1-spot` | [0] | "Cluster spots into distinct spatial domains based on gene expression profiles." | `spatial-domain-detection` | T2 |
| PB23 | `1-cell-noCT` | [0] | "What types of cells are in this tissue? It is a mouse cortex." | `celltype-annotation-fast` or `celltype-annotation-scanvi` | T1 |

| ID | Session | Target | Query | Expected skill | Task |
|---|---|---|---|---|---|
| PB24 | `1-cell` | [0] | "Segment nuclei from the DAPI staining image." | *No skill* (image processing) | out-of-scope |
| PB25 | `1-cell` | [0] | "Build a CNN to classify tissue regions from H&E images." | *No skill* (deep learning) | out-of-scope |
| PB26 | `1-cell` | [0] | "Impute dropout zeros in the expression matrix using MAGIC." | *No skill* (imputation) | out-of-scope |
| PB27 | `1-spot` | [0] | "Convert this AnnData object to a Seurat RDS file for analysis in R." | *No skill* (format conversion) | out-of-scope |
| PB28 | `1-cell` | [0] | "Perform RNA velocity analysis using spliced and unspliced counts." | *No skill* (RNA velocity) | out-of-scope |
| PB29 | `1-cell` | [0] | "Train a variational autoencoder on the gene expression data for dimensionality reduction." | *No skill* (ML training) | out-of-scope |
| PB30 | `1-cell` | [0] | "Predict patient survival outcomes from spatial gene expression features." | *No skill* (clinical prediction) | out-of-scope |

#### Supplementary Table 9 — Pipeline test queries (Set C, refined-query)

Set C tests the full pipeline (planner → filter → matcher → verifier) with `execute_code = false`. An LLM judge checks (1) result-type appropriateness, (2) parameter preservation in the refined query, (3) absence of hallucinated information, and (4) overall routing correctness. 30 queries across 4 categories.

**Category 1 — parameter preservation (PC01–PC10).**

| ID | Session | Query | Key parameters that must be preserved |
|---|---|---|---|
| PC01 | `1-spot` | "Find spatial domains in this tissue. There should be about 7 regions." | Domain count = 7 |
| PC02 | `1-cell` | "Annotate cell types in this tissue. Tissue type: mouse brain cortex." | Tissue = mouse brain cortex; no hallucinated reference path |
| PC03 | `1-spot` | "Deconvolve cell-type proportions with my reference at `./dlpfc_ref.h5ad`. The celltype column name is `cell_label`." | Reference path = `dlpfc_ref.h5ad`; column = `cell_label` (not default `celltype`) |
| PC04 | `1-cell` | "Run LIANA cell-cell communication on this mouse brain data. Species: mouse." | Species = mouse; method = LIANA (not CellPhoneDB) |
| PC05 | `1-spot` | "Run GO enrichment on these genes: MOBP, PLP1, MBP, MAG, CNP. Species: human." | All 5 gene names; species = human |
| PC06 | `1-cell` | "Annotate cell types using scANVI with reference at `./sc_ref.h5ad`. The label column is `leiden_labels`." | Method = scANVI; path = `sc_ref.h5ad`; column = `leiden_labels` |
| PC07 | `2-cell` | "Integrate these two brain sections using Harmony." | Method = Harmony (not BBKNN / Scanorama); both slices |
| PC08 | `1-cell` | "Find differentially expressed genes between excitatory and inhibitory neurons." | Groups = excitatory vs inhibitory |
| PC09 | `1-cell` | "Calculate Moran's I spatial autocorrelation for PLP1 and GFAP." | Statistic = Moran's I; genes = PLP1, GFAP |
| PC10 | `1-cell` | "Score cells for KEGG_APOPTOSIS pathway activity. Species: mouse." | Pathway = KEGG_APOPTOSIS; species = mouse |

**Category 2 — correct routing (PC11–PC18).**

| ID | Session | Query | Expected routing |
|---|---|---|---|
| PC11 | `1-cell-noCT` | "Detect spatial niches in this tissue." | type = `advice` (missing celltype prerequisite) |
| PC12 | `1-cell-noCT` | "Run cell-cell communication analysis on this tissue." | type = `skill_selection` or `advice` (missing celltype) |
| PC13 | `1-spot` | "What cell types make up each spot? I have a reference at `./ref.h5ad`." | type = `success`, skill = deconvolution (not annotation) |
| PC14 | `mixed` | "Run SpatialDE on the breast cancer slice." | type = `success`, target = slice 1 (not slice 0) |
| PC15 | `1-cell` | "What biological processes are enriched in my SVG results?" | type = `clarification` (which genes?) or `advice` |
| PC16 | `1-spot` | "Identify spatial domains in this tissue." | type = `success`; must NOT hallucinate domain count |
| PC17 | `1-cell` | "Train a deep-learning model to predict cell types from microscopy images." | type = `no_skill` (out of scope) |
| PC18 | `1-cell-noCT` | "Find differentially expressed genes between T cells and B cells." | type = `advice` (no celltype column to group by) |

**Category 3 — vague-to-specific refinement (PC19–PC24).**

| ID | Session | Query | Expected interpretation |
|---|---|---|---|
| PC19 | `1-cell` | "What genes show interesting spatial patterns?" | SVG analysis (SpatialDE or Moran's I) |
| PC20 | `1-cell` | "Are certain cell types found near each other more often than expected?" | Neighborhood-enrichment analysis |
| PC21 | `1-spot` | "Can you find the spatial structure in this tissue?" | Spatial-domain detection |
| PC22 | `1-cell` | "Group cells into tissue regions based on their location and expression." | Spatial-domain detection / clustering |
| PC23 | `1-spot` | "What is the cellular composition of each spot in this Visium data?" | Spot-level deconvolution |
| PC24 | `1-cell` | "How are cells communicating with each other in this tissue?" | Cell-cell communication analysis |

**Category 4 — no-hallucination checks (PC25–PC30).**

| ID | Session | Query | Must NOT appear in refined query |
|---|---|---|---|
| PC25 | `1-cell` | "Annotate cell types in this tissue." | Must not fabricate reference path or species |
| PC26 | `1-spot` | "Run pathway enrichment analysis." | Must not fabricate gene list |
| PC27 | `1-cell` | "Detect spatial domains." | Must not fabricate number of clusters |
| PC28 | `1-spot` | "Deconvolve this tissue." | Must not fabricate reference path |
| PC29 | `1-cell` | "Find differentially expressed genes." | Must not fabricate group names or comparison |
| PC30 | `2-cell` | "Analyze cell communication on one of the slices." | Must not pick slice arbitrarily (should clarify) |

#### Supplementary Table 10 — Pipeline-test evaluation protocol

The pipeline-stage accuracy benchmark tests three STAT pipeline stages independently across 7 LLM backbones. Sets A and B use deterministic evaluation; Set C uses an LLM judge.

##### Set A — Query planner

**Method.** Deterministic comparison of planner output against ground truth. **Success criterion** (all three must match): (1) `target_slice_ids` matches the expected slice assignments; (2) `num_steps` matches expected; (3) `needs_clarification` matches expected. **Flexible matching.** For multi-step queries where multiple valid decompositions exist, an `alt_expected` field lists acceptable alternatives. For example, "Annotate cells, find DE genes, and run GO enrichment" can be decomposed as 2 or 3 steps depending on whether GO enrichment is treated as a sub-step of DE.

##### Set B — Skill matching

**Method.** Deterministic comparison of matched skill slug against the expected set. **Success criterion.** For in-scope queries (PB01–PB23) the top-1 matched skill slug must appear in `expected_skill_slugs`; multiple acceptable skills are listed for tasks with valid alternatives (e.g., both `celltype-deconvolution` and `deconvolution-flashdeconv` for spot deconvolution). For out-of-scope queries (PB24–PB30) the skill matcher must return an empty match. **Note.** The `SkillFilter` stage is deterministic and identical across all models (filters by modality, data level, slice count); only the LLM-based `SemanticMatcher` varies between models. Skills with `default_skill = false` are excluded from the candidate pool to match the production pipeline.

##### Set C — LLM judge

**Method.** LLM-as-judge (Claude Sonnet 4) evaluates each pipeline output against the expected behaviour.

**Judge prompt.**

You are evaluating the output of a spatial transcriptomics analysis pipeline.

Original user query: `{query_text}`
Expected behavior: `{expected_behavior}`

Pipeline output:
- Result type: `{result_type}` (`success` = ready to execute with selected skill, `advice` = prerequisite missing, `skill_selection` = multiple relevant skills found asking user to choose, `verifier_clarification` = skill selected but needs more info from user, `planner_clarification` = ambiguous query needs user input, `no_skill` = out of scope)
- Selected skill: `{selected_skill}`
- Skill options (if `skill_selection`): `{skill_options}`
- Final refined query: `{final_query}`
- Verifier questions: `{verifier_questions}`
- Clarification question: `{clarification_question}`
- Advice message: `{advice_message}`

Evaluate whether the pipeline output matches the expected behavior. Consider:
1. Is the result type appropriate? `skill_selection` counts as success if the correct task type was identified. `planner_clarification` counts as success if the expected behavior says the query should ask for clarification.
2. If the refined query exists, does it preserve all user-specified parameters?
3. Does the output avoid hallucinating information not in the original query?
4. Is the overall routing correct?

Respond with ONLY `SUCCESS` or `FAIL` followed by a brief reason.

**Judge output format.** Binary `SUCCESS` / `FAIL` with a one-line justification.

##### Result types

| Type | Meaning | When it occurs |
|---|---|---|
| `success` | Ready to execute with selected skill | Skill matched, prerequisites met, refined query generated |
| `advice` | Prerequisite missing | Required data (e.g., celltype annotations) not available |
| `skill_selection` | Multiple skills matched | User needs to choose between viable alternatives |
| `verifier_clarification` | Skill selected, needs more info | Verifier identified missing parameters |
| `planner_clarification` | Query is ambiguous | Planner cannot determine target slices or intent |

| Type | Meaning | When it occurs |
|---|---|---|
| `no_skill` | Out of scope | No matching skill found in the registry |

**Accuracy calculation.** For each model, accuracy = (number of successful queries) / 30, reported as a percentage; results are reported separately for each set.

##### Supplementary Table 11 — Pipeline-test accuracy and failures by LLM backbone

Per-stage accuracy (out of 30) and failure breakdown for each of the seven backbones on the dedicated 90-query pipeline test set described in Supplementary Tables 7–10.

| Backbone model | Set A — Planner | Set B — Skill match | Set C — Refined query |
|---|---|---|---|
| Claude Sonnet 4 | 29 / 30 (96.7%) | 30 / 30 (100%) | 30 / 30 (100%) |
| Grok-3 | 30 / 30 (100%) | 30 / 30 (100%) | 29 / 30 (96.7%) |
| Grok-4 | 29 / 30 (96.7%) | 28 / 30 (93.3%) | 27 / 30 (90.0%) |
| GPT-5.4 | 29 / 30 (96.7%) | 29 / 30 (96.7%) | 27 / 30 (90.0%) |
| Gemini 3.1 Pro | 26 / 30 (86.7%) | 30 / 30 (100%) | 30 / 30 (100%) |
| DeepSeek V3.2 | 29 / 30 (96.7%) | 27 / 30 (90.0%) | 29 / 30 (96.7%) |
| GPT-4o | 30 / 30 (100%) | 29 / 30 (96.7%) | 25 / 30 (83.3%) |

**Claude Sonnet 4 — 1 failure.**

> **Set A · PA29 — SVG → pathway chain**
> **Query**: "Find spatially variable genes and then run pathway enrichment on the top SVGs."
> **What happened**: Planner returned 1 step, slices `[[0]]`; expected 2 steps, slices `[[0],[0]]`.
> **Root cause**: Planner merged the two sequential tasks (SVG detection → pathway enrichment) into a single step instead of recognising the dependency chain.

**DeepSeek V3.2 — 4 failures.**

> **Set A · PA25 — 3-step pipeline**
> **Query**: "Identify cell types, analyze cell-cell communication, and detect spatial niches."
> **What happened**: Returned 1 step instead of 2–3.
> **Root cause**: Merged all three tasks into a single step instead of decomposing the multi-step pipeline.

> **Set B · PB01 — Spatial-domain detection**
> **Query**: "Identify 7 spatial domains using graph-based clustering."
> **What happened**: Empty match (no skill selected); expected `spatial-domain-detection`.
> **Root cause**: Semantic matcher failed to bind a clear domain-detection query to the skill.

> **Set B · PB18 — Spatial structure (vague)**
> **Query**: "Can you find the spatial structure in this tissue?"
> **What happened**: Empty match.
> **Root cause**: Vague phrasing not resolved to domain detection.

> **Set B · PB23 — Cell-type annotation (vague)**
> **Query**: "What types of cells are in this tissue? It is a mouse cortex."
> **What happened**: Empty match.
> **Root cause**: Cell-type annotation query not matched to any annotation skill.

> **Set C · PC27 — Refined-query hallucination**
> **Query**: "Detect spatial domains."
> **Root cause**: Pipeline injected session-metadata details (mouse brain cortex slice 0) that the user did not provide — counts as hallucination.

**Grok-4 — 5 failures.**

> **Set A · PA23 — 3-step pipeline**
> **Query**: "Annotate cells, find DE genes between cell types, and run GO enrichment on the DE results."
> **What happened**: Returned 1 step instead of 2–3.
> **Root cause**: Merged three sequential tasks into a single step.

> **Set B · PB17 — Neighborhood enrichment (vague)**
> **Query**: "Are certain cell types found near each other more often than expected?"
> **What happened**: Empty match.
> **Root cause**: Vague co-localisation query not resolved to the neighborhood enrichment skill.

> **Set B · PB18 — Spatial structure (vague)**
> **Query**: "Can you find the spatial structure in this tissue?"
> **What happened**: Empty match.
> **Root cause**: Vague spatial-structure query not matched to domain detection.

> **Set C · PC21 — Spatial structure (vague)**
> **Query**: "Can you find the spatial structure in this tissue?"
> **Root cause**: Result type `no_skill` — incorrectly classified a vague but valid query as out of scope.

> **Set C · PC26 — Pathway enrichment**
> **Query**: "Run pathway enrichment analysis."
> **Root cause**: Result type `no_skill` — pathway enrichment is a supported skill that the matcher failed to recognise.

> **Set C · PC28 — Deconvolution**
> **Query**: "Deconvolve this tissue."
> **Root cause**: Returned `no_skill` instead of routing to deconvolution and asking the verifier for the missing reference.

**GPT-5.4 — 4 failures.**

> **Set A · PA25 — 3-step pipeline**
> **Root cause**: Same multi-step merging pattern as DeepSeek — collapsed into 1 step.

> **Set B · PB01 — Spatial-domain detection**
> **Root cause**: Empty match for a clear domain-detection query.

> **Set C · PC25 — Annotation hallucination**
> **Query**: "Annotate cell types in this tissue."
> **Root cause**: Hallucinated species (mouse) and tissue type (brain cortex) the user did not provide.

> **Set C · PC26 — Pathway enrichment**
> **Root cause**: Returned `no_skill` for a supported capability.

> **Set C · PC28 — Deconvolution**
> **Root cause**: Returned `no_skill` instead of routing to deconvolution.

**Gemini 3.1 Pro — 4 failures.**

> **Set A · PA23 — 3-step pipeline**
> **Root cause**: Merged three sequential tasks into one step.

> **Set A · PA24 — Domain → DE chain**
> **Query**: "Detect spatial domains, then find marker genes for each domain."
> **Root cause**: Merged domain detection + DE into a single step despite the explicit "then".

> **Set A · PA25 — 3-step pipeline**
> **Root cause**: Same multi-step merging pattern.

> **Set A · PA29 — SVG → pathway chain**
> **Root cause**: Merged SVG + pathway enrichment despite the explicit "then".

**Grok-3 — 1 failure.**

> **Set C · PC15 — Pathway-on-SVG ambiguity**
> **Query**: "What biological processes are enriched in my SVG results?"
> **Root cause**: Hallucinated "slice 0" and assumed SpatialDE method when the user provided no gene list or specific parameters; should have asked for clarification.

**GPT-4o — 6 failures.**

> **Set B · PB18 — Spatial structure (vague)**
> **Root cause**: Empty match for the vague spatial-structure query.

> **Set C · PC06 — Redundant verification**
> **Query**: "Annotate cell types using scANVI with reference at `./sc_ref.h5ad`. The label column is `leiden_labels`."
> **Root cause**: Verifier redundantly re-asked for the reference path and column name despite the user having already stated them.

> **Set C · PC15 — Pathway-on-SVG ambiguity**
> **Root cause**: Did not recognise the ambiguity about which gene list to enrich; should have asked for clarification.

> **Set C · PC21 — Spatial structure (vague)**
> **Root cause**: Classified as `no_skill` when spatial-domain detection is a core capability.

> **Set C · PC22 — Spatial domain (paraphrased)**
> **Root cause**: Classified as `no_skill` for a clear spatial-clustering task; matcher did not recognise the paraphrasing.

> **Set C · PC23 — Deconvolution intent**
> **Root cause**: Returned `no_skill` when spot-level deconvolution is a valid task.

##### Failure-pattern summary

| Pattern | Models affected | Queries |
|---|---|---|
| **Multi-step merging** — planner collapses sequential tasks into 1 step | All except Grok-3, GPT-4o | PA23–PA25, PA29 |
| **Vague-query rejection** — semantic matcher returns empty for valid but vague queries | DeepSeek, Grok-4, GPT-4o | PB01, PB17, PB18 |
| `no_skill` **over-rejection** — full pipeline rejects supported tasks as out-of-scope | Grok-4, GPT-5.4, GPT-4o | PC21, PC22, PC23, PC26, PC28 |
| **Hallucination in refinement** — pipeline adds session metadata to refined query | DeepSeek, GPT-5.4, Grok-3 | PC15, PC25, PC27 |
| **Redundant verification** — verifier re-asks for already-stated parameters | GPT-4o | PC06 |

#### Supplementary Table 12 — STAT skill registry

STAT exposes each analysis method as a self-describing *skill*: a directory stat_agent/skills/<slug>/ containing a single `SKILL.md` file with a YAML frontmatter that declares the skill's applicability conditions and prerequisites, and a Markdown body that contains the instructions the language model uses to invoke the underlying library. At session startup the registry loads frontmatter only; the Markdown body is loaded on demand once a skill is selected. Adding a new method requires writing one new `SKILL.md` file — no agent code changes. The current registry comprises 27 skills covering 11 analytical task families.

| Slug | Title | Underlying method | Modality | Data level | Slices | Default | Key prerequisites |
|---|---|---|---|---|---|---|---|
| **Cell-type annotation and deconvolution** | | | | | | | |
| `celltype-annotation-scanvi` | Cell-type annotation with scANVI | scANVI (scvi-tools) | gene | cell | 1 | ✓ | reference `.h5ad` + celltype column |
| `celltype-annotation-fast` | Fast cell-type annotation (Clustering + LLM) | Leiden + LLM marker annotation | gene | cell | 1 | ✓ | tissue type |
| `annotation-tangram` | Cell-type annotation via spatial mapping | Tangram | gene | cell / spot | 1 | — | reference `.h5ad` + celltype column |
| `celltype-deconvolution` | Cell-type deconvolution (RCTD) | RCTD | gene | spot | 1 | ✓ | reference + raw UMI counts |
| `deconvolution-cell2location` | Bayesian deconvolution (Cell2location) | Cell2location | gene | spot | 1 | — | reference + GPU recommended |
| `deconvolution-flashdeconv` | Fast spot deconvolution (FlashDeconv) | FlashDeconv | gene | spot | 1 | ✓ | reference + celltype column |
| **Spatial-domain identification and niche detection** | | | | | | | |
| `spatial-domain-detection` | Spatial-domain detection (SpaGCN) | SpaGCN | gene | spot | 1 | ✓ | expected #domains; H&E optional |
| `spatial-domain-stagate` | Spatial-domain detection (STAGATE) | STAGATE | gene | cell / spot | 1 | — | expected #domains (optional) |
| `spatial-domain-graphst` | Spatial-domain detection (GraphST) | GraphST | gene | spot | 1 | — | expected #domains (optional) |
| `niche-detection` | Spatial niche detection | Harmonics | gene | cell | 1 | ✓ | celltype annotations |
| **Spatially variable genes and spatial statistics** | | | | | | | |
| `svg-spatialde` | Spatially variable genes (SpatialDE) | SpatialDE / NaiveDE | gene | cell / spot | 1 | ✓ | none |
| `spatial-statistics` | Spatial statistics analysis | Moran's I, Ripley's K, co-occurrence (squidpy) | gene / protein | cell / spot | 1 | ✓ | gene names or celltypes |
| `spatial-stats-neighborhood-enrichment` | Neighborhood enrichment analysis | squidpy permutation test | gene / protein | cell / spot | 1 | ✓ | celltype annotations |
| **Cell-cell communication** | | | | | | | |
| `cell-communication-liana` | Cell-cell communication (LIANA+) | LIANA+ | gene | cell / spot | 1 | ✓ | celltype annotations + species |
| `cell-communication-cellphonedb` | Cell-cell communication (CellPhoneDB) | CellPhoneDB | gene | cell / spot | 1 | — | celltype annotations + human-only |

| Slug | Title | Underlying method | Modality | Data level | Slices | Default | Key prerequisites |
|---|---|---|---|---|---|---|---|
| **Differential expression and pathway analysis** | | | | | | | |
| `differential-expression` | Differential gene expression | scanpy `rank_genes_groups` (Wilcoxon) | gene | any | any | ✓ | grouping definition |
| `pathway-go-enrichment` | GO enrichment | gseapy (Fisher + BH) | gene | any | any | ✓ | gene list + species |
| `enrichment-ora` | Over-representation & pathway enrichment | gseapy (KEGG / Reactome / Hallmark / WikiPathways / GO) | gene | any | any | — | gene list + species |
| `pathway-enrichment-compare` | Two-group pathway enrichment comparison | gseapy.enrich (Fisher + BH) | gene | any | any | — | two gene lists + species |
| `pathway-ssgsea` | Per-cell pathway activity scoring (ssGSEA) | gseapy ssGSEA | gene | cell / spot | 1 | — | pathway / gene set + species |
| **Batch integration** | | | | | | | |
| `integration-harmony` | Batch integration (Harmony) | Harmony | gene | cell / spot | ≥2 | — | slice IDs; optional reference |
| `integration-bbknn` | Batch integration (BBKNN) | BBKNN | gene | any | ≥2 | ✓ | slice IDs |
| `integration-scanorama` | Batch integration (Scanorama) | Scanorama | gene | any | ≥2 | — | slice IDs |
| **Spatial alignment and registration** | | | | | | | |
| `alignment-stalign` | Spatial alignment (STalign) | STalign (LDDMM) | gene | cell | 2 | ✓ | landmark pairs |
| `registration-paste` | Slice registration (PASTE) | PASTE (optimal transport) | gene | any | ≥2 | — | similar tissues |
| **Trajectory and copy-number inference** | | | | | | | |
| `trajectory-pseudotime` | Pseudotime trajectory (Palantir / DPT) | Palantir / scanpy DPT | gene | cell | 1 | — | celltypes + root hint |
| `cnv-inference` | Expression-based CNV inference | infercnvpy | gene | cell / spot | 1 | — | celltypes + reference normal types |

**How the filter and verifier use this metadata.** The deterministic **skill filter** removes any skill whose `filter_requirements` are not satisfied by the loaded session — for a single-slice MERFISH session at cell resolution, for example, the spot-only deconvolution skills are removed before the language model sees the candidate list. The LLM-based **semantic matcher** then selects up to two skills whose `description` field matches the user's request, with an explicit instruction to be conservative and return an empty match for out-of-scope requests rather than pick the closest available skill. The **verifier** finally reads the `prerequisites` list and either marks them satisfied from the session state, asks the user via a clarification question, or surfaces an advice message when the prerequisite requires a prior analysis step (e.g., niche detection requires a celltype column).

### Supplementary Note 1 — STAT pipeline prompt templates

Verbatim prompt templates for each LLM-driven role in the STAT pipeline (planner, skill matcher, verifier, code generator, error reflector, analyzer). Reproduced from the production templates as of the version evaluated in this paper.

### Supplementary Note 1: STAT Pipeline Prompt Templates

This note reproduces the system prompts used by each LLM-driven role in the STAT pipeline. All placeholders enclosed in `{...}` are filled at runtime from the typed session, the registered skill metadata, the conversation memory, and the current user query. The prompts shown here are the production templates as of the version evaluated in this paper.

---

#### 1. Query planner

The planner determines which slice(s) the query targets and decomposes multi-step requests. Output is a JSON object describing either a clarification question or an explicit list of steps with `target_slice_ids` and a `refined_query`.

```text
You are a query planner for spatial transcriptomics analysis.

**Session Information:**
- Total slices: {n_slices}
{slice_descriptions}     # one bullet per slice with id / modality / data level / cell or spot count / annotations
{roi_descriptions}       # named ROIs already defined on the session
{history_context}        # summary of prior conversation turns

**User Query:** "{user_query}"

{clarification_context}  # appended only on multi-turn clarification rounds

**Your Task:**
Determine which slice(s) the user wants to analyze and how to execute the query.

**Consider:**
- Explicit slice references (e.g., "slice 0", "slice 1")
- Tissue name references (e.g., "breast cancer tissue")
- ROI references (e.g., "in tumor_region")
- Keywords like "both", "all", "each" (may need separate steps)
- Keywords like "compare", "between" (single cross-slice step)
- If only 1 slice exists, assume that slice
- If ambiguous with multiple slices, ASK FOR CLARIFICATION

**Decide:**
1. Do you have enough information to determine target slice(s)?
2. If YES: Which slice(s) and should it be one step or multiple steps?
3. If NO: What clarification question should you ask the user?

**Output Format (JSON):**
{
  "needs_clarification": true|false,
  "clarification_question": "...",
  "steps": [
    {"step_number": 1, "description": "...", "target_slice_ids": [0],
     "refined_query": "Refined query. Do NOT add information the user did not mention."}
  ]
}
```

The full prompt also contains six worked examples covering single-slice queries, ambiguous targets, "both" / "all" / "each" decomposition, cross-slice comparison, tissue-name resolution, and the single-slice fallback.

---

#### 2. Skill matcher (semantic, LLM-based)

The skill matcher receives the compatibility-filtered subset of skills and selects up to `top_k` (default = 2) whose descriptions specifically match the user's request. The matcher is intentionally conservative: it returns an empty array when no skill is a specific fit, which signals an out-of-scope request to the downstream pipeline.

```text
```

```
You are a strict skill matching system. Match skills ONLY when the user's request
SPECIFICALLY asks for what the skill provides.

User Request: "{request}"

Available Skills:
{skills_catalog}        # one line per compatible skill: `- **<slug>**: <description>`

MATCHING CRITERIA:
- ✅ Match: User's request DIRECTLY asks for the skill's specific task/output
- ❌ Don't match: Request is only loosely related or shares general themes
- ❌ Don't match: User can accomplish their goal WITHOUT this skill
- ✅ Be conservative: When in doubt, return empty array

EXAMPLES:
- "Perform niche detection" → ["niche-detection"] ✅
- "Find spatial niches in tumor" → ["niche-detection"] ✅
- "Is gene X correlated with cell type Y?" → [] ❌
- "Show spatial distribution of ERBB2" → [] ❌
- "Compare malignant cells between ROIs" → [] ❌
- "Cluster cells by spatial patterns" → [] ❌

Your task:
1. Identify the SPECIFIC task the user is asking for
2. Match ONLY if a skill provides EXACTLY that task
3. Return at most {top_k} skill slugs, or fewer if not specifically relevant
4. Respond with ONLY a JSON array: ["skill-slug"] or []

CRITICAL: Return [] (empty array) unless the skill is SPECIFICALLY needed for the user's request.

Response (JSON array only):
```

---

#### 3. Skill verifier

The verifier reads the selected skill's prerequisite list and decides whether the prerequisites are met from session state, askable from the user, or require a prior analytical step.

```text
You are a skill prerequisite verifier for spatial transcriptomics analysis.

**Task:** Verify if all prerequisites are met for executing a skill.

**Plan Step:**
- Description: {plan_step.description}
- Query: "{plan_step.refined_query}"
- Target slices: {target_slices}

**Selected Skill:** {selected_skill.name} ({selected_skill.slug})

**Skill Prerequisites:**
{prerequisites_list}

**Current Session State:**
- Total slices: {n_slices}
{slice_descriptions}
{roi_descriptions}
{user_responses_context}    # responses already collected in this clarification round
{history_context}

**Your Task:**
Analyze whether all prerequisites are met based on:
1. Current session state (what data/columns exist)
2. Information already collected from user
3. What's missing and how to obtain it

**Decision Criteria:**
- If prerequisite is already in session (e.g., celltype column exists), or can be obtained from the session (e.g.
tissue name) → MET
```

```
- If prerequisite needs simple user input (e.g., file path) → ASK USER
- If prerequisite needs prior analysis (e.g., missing celltype column, needs annotation first) → ADVISE USER

**Output Format (JSON):**
{
  "prerequisites_met": true|false,
  "missing_prerequisites": ["..."],
  "can_obtain_by_chat": true|false,
  "clarification_questions": ["...", "..."],
  "complete_query": "Complete query with all info",
  "advice": "Advice for user"
}
```

The full prompt also contains five worked examples covering all-met, ask-user, advise-user, partial-info-collected, and all-collected paths.

---

#### 4. Code generator (analysis system prompt)

The code generator runs after planning, filtering, matching, and verification have all succeeded. It receives the system prompt below, the compiled session API, the current memory snapshot, the loaded SKILL body, and the refined query as user input.

```text
You are an expert spatial transcriptomics analysis assistant. You help researchers
analyze spatial transcriptomics data using Python.

**Your capabilities:**
1. **Data Analysis**: Load, process, and analyze spatial transcriptomics data (AnnData format)
2. **ROI Analysis**: Define and analyze regions of interest
3. **Cell Type Analysis**: Study cell type distributions and spatial patterns
4. **Visualization**: Create spatial plots and visualizations
5. **Statistical Analysis**: Perform spatial statistics and neighborhood analysis

**Response Format:**
- Briefly explain what you will do (1-2 sentences). Do NOT reference code, say "this code", or describe
implementation details — the user cannot see the code. Instead, describe the analysis action.
- Write Python code in ```python``` blocks
- If code output needs biological interpretation, add on the LAST line: __INTERPRET__: <what to focus on>
- If output is self-explanatory (e.g., simple counts, plots), do NOT add __INTERPRET__

## Data Access API
Each slice is an independent data unit with unique `slice_id`. Use the explicit
slice access API:
    slice_0 = session.get_slice(0)
    adata = slice_0.adata
    image = slice_0.primary_image

Slice properties: .is_spot_level, .is_cell_level, .is_gene, .is_protein
Iteration: session.iter_slices(modality='gene'), session.iter_slices(data_level='cell')
Session info: session.slice_ids, session.modalities, session.data_levels

## Data Structure (per DataSlice)
- slice_id, modality ('gene'|'protein'), data_level ('cell'|'spot')
- adata.obs['x'], adata.obs['y']: coordinates (REQUIRED)
- adata.obs['celltype']: optional; check before use
- adata.X: expression matrix (may be sparse)
- adata.uns['spot_shape'], adata.uns['spot_diameter'] for spot data
- adata.obsm['deconv_weights'] for deconvolved spot data

## ROI Management
Per-slice ROI access:
    session.create_roi(name, slice_id, roi_definition)
    roi = session.get_roi(name)         # returns ROI object
    sub_adata = session.roi_subsets[name]   # filtered AnnData

## CRITICAL RULES
1. ALWAYS specify slice_id explicitly in analysis code
2. NEVER use a "current slice" concept — user may ask about any slice
3. Check data_level before analysis — spot vs cell requires different methods
```

````text
4. ROIs are per-slice
```

A trimmed selection of the API surface and conventions is shown above; the production prompt additionally includes
(i) a contract on imports and library availability, (ii) examples of correct API usage, (iii) common-pitfall
warnings (sparse-matrix indexing, dense conversion, scanpy public-API discipline), and (iv) the full skill body
that the matcher selected.

---

#### 5. Error reflector

The error reflector is invoked only when generated code raises an exception. It classifies the failure as `small`
(mechanically fixable) or `large` (structural / unsupportable) and returns a JSON object containing either a fixed
code block or an explanation that downstream stages surface to the user.

```text
You are debugging Python code that failed to execute.

**User's Original Question:** "{user_query}"

**Code That Failed:**
```python
{original_code}
```

**Error Message:**
{error_msg}

**Available Context:**
- ROIs: {available_rois}      # name → {slice_id, modality, n_cells}
- Columns: {available_columns}
- Has celltype column: {has_celltype}

**Your Task:**
1. Analyze why the code failed
2. Identify the root cause (wrong ROI name? missing column? logic error? wrong API?)
3. Determine if fix is SMALL or LARGE
4. Generate fixed code ONLY if fix is small

**Change Magnitude Guidelines:**
- SMALL: Typo fix, wrong variable name, incorrect API call, missing import, parameter adjustment
- LARGE: Logic rewrite, algorithm change, adding complex workarounds, restructuring code flow

**Common Issues to Check:**
- KeyError: Wrong ROI name or missing key in dictionary (SMALL fix)
- AttributeError: Wrong method or attribute name (SMALL fix)
- NameError: Variable not defined in scope (SMALL fix)
- ValueError: Invalid value for operation (may be SMALL or LARGE)
- TypeError: Wrong type for operation (may be SMALL or LARGE)
- ImportError/ModuleNotFoundError: Missing import (SMALL fix)

**Common API Fixes (SMALL changes):**
- For sparse matrix checks: Use scipy.sparse.issparse(X) or scipy.sparse.isspmatrix_csr(X)
- For scanpy: Use public API methods only (sc.pp.*, sc.tl.*, sc.pl.*)
- For AnnData: Access .X, .obs, .var directly; use .copy() for subsetting
- For session: Use session.roi_subsets[roi_name] to get ROI data
- For numpy: Always import as np, use np.asarray() for conversions

**Return JSON:**
{
  "error_type": "KeyError|AttributeError|...",
  "root_cause": "Brief explanation of what went wrong",
  "fix_strategy": "What needs to be changed",
  "change_magnitude": "small|large",
  "fixed_code": "Complete corrected Python code (only if change_magnitude is small)",
  "confidence": 0.0-1.0,
  "should_retry": true|false
}

If change_magnitude is "large" or error is unfixable, set should_retry to false.
```
````
````

The retry budget is bounded at two attempts. If the reflector classifies the failure as `large`, or if its `confidence` is below 0.7, the loop terminates and the analyzer surfaces the explanation to the user instead of running another retry.

---

#### 6. Analyzer (biological interpretation and execution audit)

After code execution the analyzer receives the user's original query, the full sequential trace of explanation / code / output / error blocks, a compact session snapshot, and the recent conversation history. It returns a structured JSON object that contains a response summary, a description of any data changes, an optional biological interpretation (only when the code generator requested one via `__INTERPRET__`), key findings, and an explicit execution-issue verdict. Crucially, the analyzer is given both the captured `stdout` and the captured `error` for each code block, so it can detect failures or partial successes that would otherwise be invisible to the conversational layer.

```text
Analyze spatial transcriptomics code execution results.

**{session_info}**            # n slices, total cells, ROI names, per-slice gene counts and celltypes

**Conversation History:**
{conversation_history}
**[End of Conversation History]**

User asked: "{user_query}"

{assembled_context}            # interleaved per code block, see below

Provide JSON response:
{
  "response_summary": "Brief summary of entire response in under 4 sentences (what was done and key results)",
  "data_changes": "Detailed description of what was added/changed in session data (columns, obsm keys, uns keys, ROIs). Be specific about names, types, and values. 2-3 sentences.",
  "interpretation": "{interpret_instruction}" | null,
  "key_findings": ["List of 1-3 specific biological findings with numbers"] | [],
  "execution_issues": {
    "has_issues": true|false,
    "issue_type": "error" | "validation_failed" | "partial_success" | "no_effect" | null,
    "explanation": "Clear explanation of what went wrong and why"
  }
}

Rules:
- response_summary: Always provide, covers entire response
- data_changes: Always provide, be detailed and specific
- interpretation: Only if requested via __INTERPRET__ (otherwise null)
- key_findings: Only if interpretation requested (otherwise empty array)
- execution_issues: Detect if code failed or did not achieve its goal
  - has_issues=true ONLY if: Python error occurred, validation failed, code had no effect, or the user's goal was NOT achieved
  - Truncated print output is NOT an issue (output is truncated at ~3000 chars for display only;
    the in-memory data is complete)
```

The `assembled_context` block is built segment-by-segment from the code generator's response, so the analyzer sees the same sequential flow the user does:

```text
**Your explanation:**
{text segment from the code generator}

**Code:**
```python
{code segment}
```

**Output:**
{stdout, cleaned and truncated to 3000 characters}

**Error:**
```

```
{stderr / exception traceback, if the block raised}
```

This dual `Output` / `Error` channel is what lets the analyzer populate `execution_issues` accurately: a block with no `stdout` but a non-empty `Error` is classified as an `error`; a block that ran without exception but failed to write the expected `obs` column is classified as `no_effect` or `partial_success` based on the `data_changes` reasoning. The structured verdict is then surfaced to the conversation layer alongside the natural-language interpretation.

---

#### Notes on prompt assembly and logging

Every LLM call is recorded by the in-process `PromptLogger` together with the call type (`planning`, `skill_matching`, `verification`, `code_generation`, `error_reflection`, `interpretation`), the input and output token counts, the latency, and the model identifier. These logs are exported as part of the session notebook so that any reported analysis can be re-traced exactly to the prompts that produced it.

### Supplementary File 1 — CRC analytic prompts, full (recipe-level) version

Complete recipe-style prompt set used by STAT to reproduce the central figures of Oliveira et al. (Nat. Genet. 2025) on the CRC Visium HD dataset ( 100K input tokens total).

````
# CRC Analytic Prompts

Prompts designed for the STAT agent to reproduce Figure 4 of
Oliveira et al. (Nat Genet 2025) on the CRC Visium HD dataset
(slices 0=P1CRC, 1=P2CRC, 2=P5CRC, spot-level, 8 µm bins).

Each prompt is self-contained: it does not rely on the agent
knowing the paper. Paste one prompt per turn into the chat.

---

## Prompt 1 — Tumor / Periphery / Tissue labeling (Fig 4a)

```
On slices 0, 1, and 2, create a new obs column `region_label`
with three categories: "Tumor", "Periphery", "Tissue". Use this
exact recipe — do not simplify any step.

Inputs
- Coordinates: adata.obs['x'], adata.obs['y'] (units: micrometers).
- Per-spot dominant cell-type label: use
  adata.obs['DeconvolutionLabel1'] if it exists, else
  adata.obs['celltype'].
- Per-slice tumor label (hard-coded, these are the dominant
  "Tumor *" category in each slice):
      slice 0 -> "Tumor II"
      slice 1 -> "Tumor III"
      slice 2 -> "Tumor IV"

Procedure (run independently for each slice)

1. tumor_mask = (label column == tumor label for this slice).
   All other spots are non-tumor for now.

2. Build a scipy.spatial.cKDTree over ALL spot coordinates
   (x, y) in this slice.

3. Seed filter. A tumor spot qualifies as a "seed" only if the
   number of OTHER tumor spots within 50 micrometers of it is
   >= 25. Spots with fewer than 25 tumor neighbours in that
   radius are dropped as seeds (they do not generate a ring).
   Use tree.query_ball_point(tumor_coords, r=50) and count how
   many returned indices are themselves tumor. Keep only seeds
   meeting the >=25 threshold.

4. Periphery set = the union of every spot returned by
   query_ball_point(seed_coords, r=50), MINUS every tumor spot.
   Tumor spots never become Periphery — they stay Tumor.

5. Assign region_label:
      - tumor_mask                      -> "Tumor"
      - in periphery set AND not tumor  -> "Periphery"
      - everything else                 -> "Tissue"

6. Store as pandas.Categorical with fixed category order
   ["Tumor", "Periphery", "Tissue"] and save to
   adata.obs['region_label']. Also write the tumor label used
   to adata.uns['region_tumor_label'].

7. Print per-slice counts of the three categories.

Then make one figure: a 1x3 panel, one subplot per slice,
scatter of (x, y) coloured by region_label
(Tumor=red, Periphery=blue, Tissue=lightgrey), s=0.5, square
aspect, y-axis inverted. Title each subplot with the slice id
and tumor label. Use matplotlib only; do not save to disk.
```
````

---

#### Prompt 2 – Cell-type composition of Periphery vs Tissue (Fig 4b)

Run after Prompt 1 has completed successfully. It depends on `adata.obs['region_label']` existing on all three slices.

```
Using the `region_label` column you added in the previous step
(categories: Tumor, Periphery, Tissue), compute the cell-type
composition of each region for each of slices 0, 1, 2. Follow
the exact recipe below.

Label source
- adata.obs['celltype']. Call this column `label`.

Per-slice computation
For each slice in [0, 1, 2]:

1. Build a contingency table: rows = region_label
   (Tumor/Periphery/Tissue), columns = label values.
2. Row-normalize so that EACH ROW sums to 100. This gives, for
   every region, the percentage of that region occupied by each
   cell type. Do NOT normalize by column – we want "what is
   the Periphery made of", not "where does each cell type live".
3. Drop the Tumor row. Keep only Periphery and Tissue rows.
4. Reshape to long format with columns:
   [slice_id, patient, region ("Periphery"|"Tissue"), celltype,
    pct]. Map slice_id 0/1/2 to patient labels "P1CRC",
   "P2CRC", "P5CRC".

Combine the three per-slice long tables into one DataFrame
`comp_df`. Print `comp_df.head(20)`.

Cell-type selection for the plot
- From comp_df, keep only cell types whose maximum Periphery
  percentage across the three patients is >= 1%. Also drop any
  label that starts with "Tumor".
- Order celltypes on the y-axis by their mean Periphery
  percentage across patients, descending (most-enriched at top).

Plot (single matplotlib figure, reproduces Fig 4b layout)
- One axes. y-axis = celltype (categorical, ordered as above).
  x-axis = pct (0 to max, linear).
- Draw one dot per (patient, region, celltype) row.
- Color by region: Periphery = "#1f77b4" (blue),
  Tissue = "#9e9e9e" (grey).
- Marker shape by patient: P1CRC="o", P2CRC="s", P5CRC="^".
- Dot size ~40, alpha 0.9. Add a light horizontal gridline at
  each celltype row. Legend with two groups: one for region
  (color), one for patient (shape). Figure size ~ (6, 8).
  Title: "Cell-type composition: Periphery vs rest of tissue".
  plt.show().

Also store comp_df on each slice as
adata.uns['region_composition'] (filter to that slice's rows)
so later steps can reuse it.
```

---

#### Prompt 3 – SPP1+ vs SELENOP+ macrophage subpopulations (Fig 4c/d)

Run after Prompt 1. Depends on `adata.obs['region_label']` existing on all three slices. Goal: split macrophage-labeled spots inside the tumor periphery into two transcriptomic subclusters per patient and label them by marker gene.

```
For each of slices 0, 1, 2, identify two macrophage
subpopulations ("SPP1+" and "SELENOP+") among the spots that
lie in the tumor periphery, and write the result to a new obs
```

```
column `macrophage_subtype`. Follow this recipe exactly.

Subset definition (per slice, independently)
- Consider spots where ALL of the following hold:
    adata.obs['region_label']       == "Periphery"
    adata.obs['celltype']           == "Macrophage"
  Call this index set `mac_idx`. If it has fewer than 30 spots
  or the gene SPP1 is absent from adata.var_names, skip this
  slice (every spot becomes "Other") and print a warning.

Labeling rule (expression-based, per anchor gene)
Each label REQUIRES a positive raw count in its own anchor
gene. "SELENOP+" is NOT simply "SPP1 == 0".

1. Read raw SPP1 and SELENOP counts for every spot in mac_idx
   (densify if sparse). Call them spp1_v and seln_v.

2. Define three masks:
      spp1_only_mask = (spp1_v > 0) & (seln_v == 0)
      seln_only_mask = (seln_v > 0) & (spp1_v == 0)
      both_mask      = (spp1_v > 0) & (seln_v > 0)
   Spots where both counts are zero, AND spots in both_mask,
   are ambiguous and will be left as "Other".

3. Compute the per-slice SPP1-only fraction:
      frac = spp1_only_mask.sum() / len(mac_idx)

4. Per-slice minimum-fraction gate:
   If frac < 0.05, this slice has no genuine SPP1+
   subpopulation. Do NOT assign any "SPP1+":
      - every spot in seln_only_mask OR both_mask -> "SELENOP+"
      - everything else -> "Other"

5. Otherwise (frac >= 0.05), apply the exclusive labels:
      spp1_only_mask  -> "SPP1+"
      seln_only_mask  -> "SELENOP+"
      both_mask       -> "Other"   (ambiguous)
      neither         -> "Other"   (no anchor evidence)

Write-back to the full slice
- Create adata.obs['macrophage_subtype'] as a pandas
  Categorical with categories ["SPP1+", "SELENOP+", "Other"].
  Default every spot to "Other".
- Assign "SPP1+" / "SELENOP+" to spots in mac_idx according to
  the rule above (or all "SELENOP+" if the fraction gate
  triggered).
- Store adata.uns['macrophage_fallback_used'] = True/False so
  Prompt 4 can read it.

Reporting
- Print, per slice: number of periphery-macrophage spots,
  the raw SPP1+ fraction, the final SPP1+ / SELENOP+ counts,
  and whether the fraction-gate fallback triggered.

Plot
- 1x3 matplotlib figure, one subplot per slice. On each
  subplot, scatter ALL spots in light grey (s=0.3, alpha=0.3)
  as background, then overlay macrophage_subtype spots:
  "SPP1+" in "#d62728" (red), "SELENOP+" in "#2ca02c"
  (green). s=6, alpha=0.9. Equal aspect, y-axis inverted.
  Title each subplot with the slice id, the tumor label from
  adata.uns['region_tumor_label'], and the two subtype counts.
  plt.show(). Do not save to disk.
```

---

#### Prompt 4 – Marker dot plot for SPP1+ and SELENOP+ macrophages (Fig 4c)

Run after Prompt 3. Depends on `adata.obs['macrophage_subtype']`
(categories: "SPP1+", "SELENOP+", "Other") existing on all
three slices. Goal: find differential marker genes between the

two macrophage subpopulations across patients and draw a
dot plot with gene × (subtype, patient) cells.

```
Build a differential-expression dot plot comparing "SPP1+" and
"SELENOP+" macrophage spots across the three patients. Follow
this recipe exactly — do not skip steps.

Assemble the combined object
1. For each slice in [0, 1, 2]:
   - Select spots where adata.obs['macrophage_subtype'] is in
     {"SPP1+", "SELENOP+"} (i.e. drop "Other").
   - Add two new obs columns on that subset:
       patient = {"P1CRC","P2CRC","P5CRC"}[slice_id]
       subtype = adata.obs['macrophage_subtype']
2. Concatenate the three subset AnnDatas into one object
   `mac_all` using anndata.concat(..., join="inner",
   index_unique="-"). Keep ONLY these obs columns:
   patient, subtype. Ensure `mac_all.X` holds raw integer
   counts — if not, copy counts from layers['counts'] or reload.
3. Print mac_all.shape and a crosstab of subtype x patient.

Normalize + differential expression on the merged object
4. sc.pp.normalize_total(mac_all, target_sum=1e4)
5. sc.pp.log1p(mac_all)
6. Save the log-normalized matrix as mac_all.layers['lognorm'].
7. sc.tl.rank_genes_groups(mac_all, groupby="subtype",
   method="wilcoxon", pts=True, use_raw=False)
8. For each of the two groups ("SPP1+", "SELENOP+"), pull
   genes with pvals_adj < 0.05 and logfoldchanges > 0.1, then
   take the top 10 by logfoldchanges. Deduplicate while
   preserving order. Call this `marker_genes`. Print the two
   per-group top-10 lists and the final marker_genes list.
   Expect ~15-20 genes total.

Per (subtype × patient) dot statistics
9. For every combination of (group in ["SELENOP+","SPP1+"],
   patient in ["P1CRC","P2CRC","P5CRC"]) and every gene in
   marker_genes, compute on mac_all.layers['lognorm']:
     - pct_expressed = 100 * (gene values > 0).mean() over
       the spots in that (group, patient) cell.
     - mean_expr = mean of expm1(lognorm) over those spots
       (i.e. mean of the denormalized per-10k expression).
     - log_mean_expr = log1p(mean_expr).
   Skip any (group, patient) cell that has zero spots (this
   will happen for "SPP1+" in P5CRC). Collect rows in a
   DataFrame `dot_df` with columns
   [gene, group, patient, pct_expressed, log_mean_expr].

10. Apply dot.min filter: drop rows with pct_expressed < 15.
    This removes low-frequency noise and makes the P5CRC SPP1+
    column naturally empty (P5 has no SPP1+ macrophages).

Ordering for the plot
11. Genes on the y-axis: keep the order from marker_genes,
    but place SPP1+ markers first (top of the plot) and
    SELENOP+ markers second (bottom). Concretely: sort genes
    so that genes exclusive to the SPP1+ top-10 come first,
    shared genes go into the SPP1+ block, then SELENOP+
    exclusives last.
12. x-axis columns: six cells in this left-to-right order
      SELENOP+ | P1CRC
      SELENOP+ | P2CRC
      SELENOP+ | P5CRC
      SPP1+    | P1CRC
      SPP1+    | P2CRC
      SPP1+    | P5CRC
    (the P5CRC SPP1+ column will be empty of dots — that is
    expected; still draw the x-tick.)

Plot
13. Single matplotlib axes, figure size ~ (5.5, 7).
```

```
    - x-axis: categorical, the six (group, patient) cells
      above, with tick labels "P1CRC"/"P2CRC"/"P5CRC"
      (rotated 90°) and a top-axis label grouping three cols
      under "SELENOP+ macrophage" and three cols under
      "SPP1+ macrophage". Draw a vertical dashed grey line
      between the two blocks.
    - y-axis: gene names in italics (use fontstyle="italic"),
      ordered as defined above.
    - Scatter each dot at (x, y) with:
        size  = pct_expressed mapped to marker area, e.g.
                s = pct_expressed * 6
        color = log_mean_expr, cmap="viridis",
                vmin=0, vmax=7
    - Two legends on the right: a colorbar labeled
      "log (mean expression)" and a size legend labeled
      "Percentage expressed" with reference dots at 20, 40,
      60, 90.
    - No axes frame on top/right; light horizontal gridlines.
    - plt.tight_layout(); plt.show(). Do not save to disk.

Also store:
- adata.uns['macrophage_subtype_de'] (on each slice, same
  copy) = the full dot_df as a DataFrame.
- adata.uns['macrophage_marker_genes'] = list marker_genes.
```

---

#### Prompt 5 — Pathway enrichment: SPP1+ vs SELENOP+ (Fig 5a)

Run after Prompt 4. Goal: compare which MSigDB Hallmark pathways are enriched in each macrophage subpopulation's marker genes and draw a mirrored bar plot.

Note: the agent's `pathway-enrichment-compare` skill handles the enrichment + plot once gene lists are stored in `adata.uns['enrichment_genes_groups']`. This prompt only needs to prepare the gene lists; the skill does the rest.

```
First, compute differential marker genes between SPP1+ and
SELENOP+ macrophages, then run two-group pathway enrichment.

Step 1: Prepare marker gene lists
- Pool spots with macrophage_subtype in {"SPP1+","SELENOP+"}
  across slices 0, 1, 2 into one AnnData (anndata.concat,
  join="inner"). Add obs columns subtype and patient.
- sc.pp.filter_genes(min_cells=max(3, int(0.05 * n_obs))).
  normalize_total(target_sum=1e4), log1p, save layer 'lognorm'.
- sc.tl.rank_genes_groups(groupby='subtype',
  method='wilcoxon', pts=True, layer='lognorm',
  use_raw=False).
- For each group, take the top 250 genes with
  logfoldchanges > 0.1, sorted by logfoldchanges descending.
  No padj filter.

Step 2: Store on slice 0's adata:
    adata.uns['enrichment_genes_groups'] = {
        "SPP1+": spp1_markers,
        "SELENOP+": selenop_markers,
    }

Step 3: Run two-group pathway enrichment comparison against
MSigDB_Hallmark_2020 (human). Use top_n=10 per side,
p_threshold=1e-3. Colors: SPP1+ orange "#D35D05" (left),
SELENOP+ green "#01702E" (right). Labels on the opposite
blank side from their bars.
```

---

#### Prompt 6 — Tumor expression near macrophage subpopulations (Fig 5b)

Run after Prompt 3. Depends on `adata.obs['macrophage_subtype']` and `adata.obs['region_label']` on all three slices. Goal: for each macrophage subtype, find the densest spatial cluster of that subtype, select nearby tumor spots, run DE between the two tumor groups, and draw violin plots of the top genes.

```
For each macrophage subpopulation (SPP1+ and SELENOP+), find
the tumor spots spatially adjacent to the densest cluster of
that subtype, then compare gene expression between the two
tumor groups. Follow this recipe exactly.

Per slice, per macrophage subtype (independently)

1. Select macrophage spots of the subtype:
     mac_coords = coordinates of spots where
     macrophage_subtype == "SPP1+" (or "SELENOP+").
   If fewer than 5 spots, skip this (slice, subtype) pair.

2. Find the densest macrophage cluster via 2D KDE:
     from scipy.stats import gaussian_kde
   Evaluate the KDE on a 200x200 grid spanning the spot
   coordinate range. The grid cell with the maximum density
   value is the "peak" — the center of the densest niche.

3. Select all spots (any cell type) within 350 micrometers
   of the peak.

4. Among those, keep only TUMOR spots (celltype == the
   per-slice tumor label from adata.uns['region_tumor_label'])
   that are within 50 micrometers of any macrophage spot in
   the 350 µm region. Use a cKDTree on the region macrophage
   coords and query the tumor coords with r=50.

5. Label these tumor spots with:
     CellType = "SPP1" or "SELENOP" (which macrophage niche)
     Patient  = "P1CRC" / "P2CRC" / "P5CRC"
   Collect across all slices into a list of AnnData subsets.

Merge and DE

6. Concatenate all tumor subsets (anndata.concat, join="inner",
   index_unique="-").
7. Exclude mitochondrial genes (var_names starting with "MT-")
   — they reflect cell quality / patient batch, not niche
   biology.
8. sc.pp.filter_genes(min_cells=max(3, int(0.05 * n_obs))).
   normalize_total(target_sum=1e4), log1p, save layer 'lognorm'.
9. sc.tl.rank_genes_groups(groupby='CellType',
   method='wilcoxon', pts=True, layer='lognorm',
   use_raw=False).
10. For each group, take top 5 genes by pvals_adj (ascending)
    with padj < 0.05 and logfoldchanges > 0.1.
    Print both top-5 lists.

Violin plot

11. For the top genes (up to 10 total, deduped), draw a
    violin plot grid:
    - Faceted by gene (5 columns, as many rows as needed).
    - x-axis per facet: two groups "SELENOP" and "SPP1".
    - Within each group, three violins side by side colored
      by Patient: P1CRC="#EE7600" (orange),
      P2CRC="#008B8B" (teal), P5CRC="#483D8B" (purple).
    - Use matplotlib violinplot, set face/edge color per
      patient. scale="width" style.
    - y-axis: "Normalized log expression".
    - Gene name as subplot title in italics.
    - Legend for Patient colors.
    - Figure size ~(15, 3.2 * nrows). plt.show().
      Do not save to disk.
```

```

---

## Prompt 7 — Ligand-receptor analysis: LIANA (Fig 5e/f)

Run after Prompt 3. Depends on `macrophage_subtype`, `region_label`, and `region_tumor_label`. Goal: run LIANA on the TME boundary per patient, draw faceted LR dotplots.

Note: the agent's `cell-communication-liana` skill handles the core `li.mt.rank_aggregate` call and basic dotplot. This prompt adds custom TME subsetting and per-patient execution that the skill does not cover.

```
Run LIANA ligand-receptor analysis on the tumor boundary for
each patient separately, then merge results and draw faceted
dotplots for each macrophage subpopulation as source.

Per slice (independently for slices 0, 1, 2)

Step 1: Build TME subset
- Periphery spots: region_label=="Periphery".
- Boundary Tumor spots: celltype == tumor label AND >= 5
  Periphery neighbors within 50 µm (cKDTree query_ball_point).
- Combine into one AnnData subset.

Step 2: Custom CellType column on the subset
- macrophage_subtype "SPP1+" → "Macrophage_SPP1+"
- macrophage_subtype "SELENOP+" → "Macrophage_SELENOP+"
- macrophage_subtype "Other" AND celltype "Macrophage" → DROP
  (they dilute SELENOP+ signal — express neither anchor gene).
- celltype == tumor_label → "Tumor"
- Everything else → keep original celltype.

Step 3: Subsample cell types > 3000 spots to 3000 (seed=42).

Step 4: Run LIANA per patient (no plot — just store results)
- adata.raw = adata.copy(); normalize_total; log1p.
- li.mt.rank_aggregate(groupby='celltype',
  resource_name='consensus', expr_prop=0.1, min_cells=3,
  n_perms=1000, use_raw=True).
- Compute aggregate_rank = (magnitude_rank +
  specificity_rank) / 2.
- Rename tumor targets to "Tumor".
- Store the FULL result DataFrame (with aggregate_rank and
  Patient columns) in adata.uns['liana_res'].
  Do NOT plot at this step.

After all slices

Step 5: Merge and plot
- Read liana_res from each slice, concatenate.
- FILTER to aggregate_rank <= 0.05 FIRST (keep only
  significant interactions), THEN for each source in
  ["Macrophage_SELENOP+", "Macrophage_SPP1+"], take top 12
  LR pairs by mean aggregate_rank among significant
  interactions only.
- Draw a faceted dotplot per source:
  Targets: ["CD4 T cell","CD8 T cell","Tumor"].
  Faceted by Patient. Dot size = significance,
  color = lr_means (viridis). plt.show().
```

---

## Prompt 8 — Goblet-cell sub-clustering

Standalone: depends only on the three CRC spot slices.

```

```
Goal: uncover heterogeneity within the goblet-cell compartment by
unsupervised clustering of the goblet spots pooled across the 3
patients, then show a marker dotplot and a per-patient spatial map
of the clusters. Follow this recipe exactly.

Slices: 0 (P1CRC), 1 (P2CRC), 2 (P5CRC). Each has raw integer
counts in adata.X.

Step 1: Build a pooled Goblet-only AnnData.
For each slice:
- Keep only spots where adata.obs['celltype'] == "Goblet".
- Tag the subset with a "Patient" obs column
  ("P1CRC"/"P2CRC"/"P5CRC") and make the obs_names unique by
  appending "-<Patient>".
Intersect var_names to common genes across the three slices, then
concatenate (anndata.concat) into one AnnData called `gob`.

Step 2: Seurat-equivalent pipeline on `gob`.
Save the raw counts: gob.layers['counts'] = gob.X.copy().
- sc.pp.normalize_total(gob, target_sum=1e4)
- sc.pp.log1p(gob)
- gob.layers['lognorm'] = gob.X.copy()   # needed for DE/dotplot
- sc.pp.highly_variable_genes(gob, flavor='seurat_v3',
                               n_top_genes=2000, layer='counts')
- sc.pp.scale(gob, max_value=10, zero_center=True)
- sc.tl.pca(gob, n_comps=30, use_highly_variable=True)
- sc.pp.neighbors(gob, n_neighbors=20, n_pcs=12)
- sc.tl.louvain(gob, resolution=0.35, random_state=0,
                key_added='cluster')
  (Note: scanpy's KNN-graph Louvain is coarser than Seurat's
   SNN-graph Louvain at the same nominal resolution, so
   resolution=0.35 here is tuned to produce ~7-8 clusters on this
   data; do NOT change this.)
Restore log-normalized values to X for DE:
    gob.X = gob.layers['lognorm']
Cast gob.obs['cluster'] to an integer-coded categorical.

Step 3: Marker dotplot.
Genes (fixed order, y-axis bottom→top):
    ["MUC2","FCGBP","TFF3","CLCA1","OLFM4",
     "DUOX2","DMBT1","REG1A","REG1B"]
For every (gene, cluster) cell:
- pct = fraction of cluster's cells with X > 0 (percent, 0-100).
- mean expression in linear space: mean(expm1(X)) over the
  cluster's cells. Then z-score across clusters per gene and clip
  to [-2.5, 2.5]. Call this "scaled".
- Hide dots where pct < 5.
Plot: clusters on x-axis, genes on y-axis (italic labels), dot
size ∝ pct, dot color ∝ scaled using RdBu_r (blue→white→red).
Add a "Scaled expression" colorbar and a "Percentage expressed"
size legend with 25 / 50 / 75 % reference dots.

Step 4: Per-patient spatial map of the goblet clusters.
Cluster color palette (index by cluster id, mod len):
    ["#CA3142","#F09235","#FEFF54","#0C00C5",
     "#60B177","#EEE697","#74140C"]
For each of the 3 slices, pull the full-slice obs (x, y,
celltype):
- Plot every spot in lightgrey, s=0.4 (background).
- Overlay Goblet spots coloured by their cluster assignment,
  s=1.2, drawing clusters in ascending order so larger clusters
  sit on top.
- Equal aspect, y-axis inverted, no ticks, no spines, title =
  patient id.
One figure: 1×3 subplot row, shared cluster legend on the right.
plt.show().

Step 5: Print top positive markers per cluster.
Run sc.tl.rank_genes_groups(gob, groupby='cluster',
    method='wilcoxon', use_raw=False, pts=True), filter each
cluster's top genes by padj<0.05, log2FC>=0.25 and
(pts - pts_rest) >= 0.1, and print the top 8 per cluster.
```

```
scanpy stores the full rank_genes table in
gob.uns['rank_genes_groups'].
```

### Supplementary File 2 — CRC analytic prompts, conversational (slide-level) version

Conversational, slide-level prompt set used by STAT to reproduce the same CRC figures with roughly two orders of magnitude fewer input tokens.

````
# CRC Analytic Prompts — Simple Version

Conversational prompts for the STAT agent on the three CRC
Visium HD slices (slice 0 = P1CRC, slice 1 = P2CRC,
slice 2 = P5CRC, 8 µm spot-level).

---

## Prompt 1 — Tumor / Periphery / Tissue

```
On slices 0, 1 and 2, I want to label every spot as
"Tumor", "Periphery", or "Tissue". The dominant tumor
label is different in each slice:
  slice 0 → "Tumor II"
  slice 1 → "Tumor III"
  slice 2 → "Tumor IV"

Tumor spots are the ones whose celltype matches that
slice's tumor label. Periphery spots are non-tumor spots
within 50 µm of a *core* tumor spot — and a tumor spot
only counts as core if it has at least 25 other tumor
spots within 50 µm of it (so isolated tumor bins don't
seed a periphery). Everything else is Tissue.

Save this as a new obs column and show me the regions on
the slices.
```

---

## Prompt 2 — Composition of Periphery vs Tissue

```
What is the cell-type composition of the Periphery vs
the Tissue regions? Row-normalise — i.e. tell me what
fraction of the Periphery is each celltype, and the same
for Tissue. Drop the Tumor region from the comparison.
Combine across the three patients into one plot.
```

---

## Prompt 3 — Macrophage SPP1+ / SELENOP+ subtyping

```
Within the Periphery, split the Macrophage spots into
two populations based on exclusive raw-count expression
of their two anchor genes:

  SPP1 > 0 AND SELENOP == 0  →  SPP1+
  SELENOP > 0 AND SPP1 == 0  →  SELENOP+
  otherwise                  →  Other

Store the call in obs['macrophage_subtype']. If a slice
has fewer than 5 % of its qualifying macrophages in one
of the subtypes, just call all of that slice's
macrophages the majority subtype. Print and plot the
per-slice counts.
```

---

## Prompt 4 — SPP1+ vs SELENOP+ marker dotplot

```
Pool the SPP1+ and SELENOP+ macrophage spots across the
three slices into one object (drop "Other"), keep
````
````

patient and subtype on each spot, and find the
differentially expressed marker genes between the two
subtypes (Wilcoxon, top ~10 per side by log fold-change,
padj<0.05).

Show those markers on a dotplot with one column per
(subtype, patient) — six columns in total — and one row
per gene. Note: P5CRC has no SPP1+ macrophages, so its
SPP1+ column will be empty.
```

---

## Prompt 5 — Pathway enrichment, SPP1+ vs SELENOP+

```
Take the Wilcoxon positive markers of Macrophage-SPP1+
and Macrophage-SELENOP+ (Periphery spots, padj<0.05,
log2FC≥0.25, pooled across the three slices) as two gene
lists and run a two-group hallmark-pathway enrichment
comparison between them.
```

---

## Prompt 6 — Tumor expression near macrophage niches

```
For each slice, find where the SPP1+ macrophages are
most spatially concentrated, and the same for the
SELENOP+ macrophages. Take a 350 µm circular region
around each of those hotspots and, inside each region,
keep only the tumor spots that sit within 50 µm of any
macrophage in that region (i.e. tumor immediately
abutting the niche).

I want to see whether tumor near SPP1+ macrophages
behaves differently from tumor near SELENOP+ macrophages.
Pool these two niche-tumor groups across patients and
run differential expression between them; show the top
markers as one violin per gene, faceted by niche, with
patients shown side by side within each facet. Drop
mitochondrial genes (MT-) before reporting.
```

---

## Prompt 7 — Cell-cell communication at the TME boundary

```
Per slice, build a TME AnnData with:
- all Periphery spots, plus
- tumor spots that have at least 5 Periphery spots within
  50 µm of them (the boundary tumor only — drop deep
  tumor).

Use the SPP1+/SELENOP+ macrophage labels from before for
those spots; the included tumor spots are just "Tumor";
every other spot keeps its original celltype. Drop
Macrophage-Other, and also drop any Macrophage spot that
didn't get a SPP1+/SELENOP+ label.

Run a ligand-receptor / cell-cell communication
analysis per slice and store the full per-pair table

After all three slices: merge, filter to
aggregate_rank ≤ 0.05 first, then for each macrophage
source pick the top 12 LR pairs by mean aggregate_rank.
Show the result faceted by patient, with targets =
CD4 T cell, CD8 T cell, Tumor.
```
```

---

#### Prompt 8 — Goblet-cell sub-clustering

Standalone — no prior prompt needed.

```
Pool every Goblet spot across slices 0/1/2 into one
AnnData (tag obs['Patient']) and run standard scanpy
clustering on the pooled object — using the first 12 PCs
when building the neighbor graph, and pick a Louvain
resolution that lands at about 7-8 clusters (~0.35 works
on this data).

Then show me:
1. A dotplot of canonical goblet / colonocyte markers
   across clusters — including the inflammatory markers
   REG1A, REG1B, DUOX2, DMBT1.
2. A per-patient spatial scatter coloured by cluster,
   non-Goblet spots drawn grey.
3. Top Wilcoxon markers per cluster.
```
